# The Injured Sciatic Nerve Atlas (iSNAT), Insights into the Cellular and Molecular Basis of Neural Tissue Degeneration and Regeneration

**DOI:** 10.1101/2022.06.26.497651

**Authors:** Xiao-Feng Zhao, Lucas D. Huffman, Hannah Hafner, Mitre Athaiya, Matthew Finneran, Ashley L. Kalinski, Rafi Kohen, Corey Flynn, Ryan Passino, Craig Johnson, David Kohrman, Riki Kawaguchi, Lynda Yang, Jeff Twiss, Daniel H. Geschwind, Gabriel Corfas, Roman J. Giger

## Abstract

Upon trauma, the adult murine PNS displays a remarkable degree of spontaneous anatomical and functional regeneration. To explore extrinsic mechanisms of neural repair, we carried out single cell analysis of naïve mouse sciatic nerve, peripheral blood mononuclear cells, and crushed sciatic nerves at 1-day, 3-days, and 7-days following injury. During the first week, monocytes and macrophages (Mo/Mac) rapidly accumulate in the injured nerve and undergo extensive metabolic reprogramming. Proinflammatory Mo/Mac in the injured nerve show high glycolytic flux compared to Mo/Mac in blood and dominate the early injury response. They subsequently give way to inflammation resolving Mac, programmed toward oxidative phosphorylation. Nerve crush injury causes partial leakiness of the blood-nerve-barrier, proliferation of endoneurial and perineurial stromal cells, and accumulation of select serum proteins. Micro-dissection of the nerve injury site and distal nerve, followed by single-cell RNA-sequencing, identified distinct immune compartments, triggered by mechanical nerve wounding and Wallerian degeneration, respectively. This finding was independently confirmed with *Sarm1^-/-^* mice, where Wallerian degeneration is greatly delayed. Experiments with chimeric mice showed that wildtype immune cells readily enter the injury site in *Sarm1-/-* mice, but are sparse in the distal nerve, except for Mo. We used CellChat to explore intercellular communications in the naïve and injured PNS and report on hundreds of ligand-receptor interactions. Our longitudinal analysis represents a new resource for nerve regeneration, reveals location specific immune microenvironments, and reports on large intercellular communication networks. To facilitate mining of scRNAseq datasets, we generated the injured sciatic nerve atlas (iSNAT): https://cdb-rshiny.med.umich.edu/Giger_iSNAT/

## Introduction

Axonal injury caused by trauma or metabolic imbalances triggers a biochemical program that results in axon self-destruction, a process known as Wallerian degeneration (WD). In the peripheral nervous system (PNS), WD is associated with nerve fiber disintegration, accumulation of myelin ovoids, reprogramming of Schwann cells (SC) into repair (r)SC, and massive nerve inflammation. Fragmented axons and myelin ovoids are rapidly cleared by rSC and professional phagocytes, including neutrophils and macrophages (Mac) (Jang et al., 2016; Klein and Martini, 2016; Perry and Brown, 1992; Rotshenker, 2011). In the mammalian PNS, timely clearance of fiber debris during WD stands in stark contrast to the central nervous system (CNS), where upon injury, clearance of degenerated axons and myelin is protracted and accompanied by prolonged inflammation (Bastien and Lacroix, 2014; Vargas and Barres, 2007).

Successful PNS regeneration depends upon the coordinated action and communication among diverse cell types, including fibroblast-like structural cells, vascular cells, SC, and different types of immune cells (Cattin et al., 2015; Chen et al., 2021; Girouard et al., 2018; Pan et al., 2020; Stratton et al., 2018). Nerve trauma not only results in fiber transection, but also causes necrotic cell death and release of intracellular content at the site of nerve injury. Depending on severity, physical nerve trauma results in vasculature damage, endoneurial bleeding, breakdown of the blood-nerve-barrier (BNB), and tissue hypoxia (Cattin et al., 2015; Rotshenker, 2011). Distal to the injury site, where mechanical damage is not directly experienced, severed axons undergo WD. The vast majority of immune cells in the injured PNS are blood-borne myeloid cells, including neutrophils, monocytes (Mo), and Mac (Kalinski et al., 2020; Ydens et al., 2020). Entry into the injured nerve causes rapid activation and acquisition of specific phenotypes. A comparative analysis between circulating immune cells in peripheral blood, and their descends in the injured nerve, however, has not yet been carried out.

It is well established that the immune system plays a key role in the tissue repair response (Bouchery and Harris, 2019). Following PNS injury, the complement system (Ramaglia et al., 2007) and Natural killer cells (NK) promote WD of damaged axons (Davies et al., 2019). Mac and neutrophils phagocytose nerve debris, including degenerating myelin and axon remnants (Chen et al., 2015; Kuhlmann et al., 2002; Lindborg et al., 2017; Rosenberg et al., 2012). Mac protect the injured tissue from secondary necrosis by clearing apoptotic cells through phagocytosis, a process called efferocytosis (Boada-Romero et al., 2020; Greenlee-Wacker, 2016; Lantz et al., 2020). In the injured sciatic nerve, efferocytosis readily takes place and is associated with inflammation resolution (Kalinski et al., 2020). The highly dynamic nature of intercellular communications, coupled with spatial differences in the nerve microenvironment, make it difficult to untangle the immune response to nerve trauma from the immune response to WD. While previous studies have employed single-cell RNA sequencing (scRNAseq) to describe naïve or injured PNS tissue (Carr et al., 2019; Kalinski et al., 2020; Wang et al., 2020; Wolbert et al., 2020; Ydens et al., 2020), a longitudinal study to investigate transcriptomic changes and cell-cell communication networks, has not yet been carried out. Moreover, a comparative analysis of cell types and transcriptional states at the nerve injury site versus the distal nerve, does not yet exist.

To better understand cellular functions, immune cell trafficking, and intercellular communication, we carried out a longitudinal study using scRNAseq and validated gene expression data by flow cytometry, ELISA, *in situ* hybridization, and immunofluorescence labeling of nerve sections. Peripheral blood mononuclear cells from naïve mice were analyzed by scRNAseq to identify leukocytes that enter the nerve upon injury and to assess their metabolic profile. To distinguish between the immune response to nerve trauma and WD, we employed *Sarm1* null (*Sarm1^-/-^)* mice in which WD is greatly delayed (Osterloh et al., 2012). We show that in injured *Sarm1^-/-^* mice blood-borne Mac rapidly accumulate at the nerve crush site. However, in the absence of WD, inflammation in the distal nerve is greatly reduced. While monocytes are found in the distal nerve of *Sarm1^-/-^* mice at 7days following injury, they fail to mature into Mac. In sum, we report temporal changes in the cellular and metabolic landscape of the injured mammalian PNS, identify spatially distinct, yet overlapping immune responses to nerve wounding and WD, and provide insights into WD elicited nerve inflammation.

## Results

### The cellular and molecular landscape of naïve sciatic nerve

To gain insights into extrinsic mechanisms associated with neural degeneration and regeneration, we carried out a longitudinal analysis of injured mouse sciatic nerve tissue using single cell transcriptomics. We first captured CD45^+^ immune cells from sciatic nerve trunk, and in parallel, CD45^-^ non-immune cells for scRNAseq analysis (**Fig. 1A**). The resulting datasets were combined, high-quality cells identified, and subjected to dimensional reduction using the top 30 principal components (**Table. S1**). Uniform manifold approximation and projection (UMAP) was used for visualization of cell clusters and clusters determined using the Louvain algorithm with a resolution of 0.5, revealing 24 clusters (**Fig. 1B**). Marker gene expression analysis was used for cell type identification (**Fig. 1C**). Stromal cells, often referred as structural cells, are abundant in the naïve nerve. They include epineurial fibroblasts (Fb1-Fb3, clusters 7-9), identified by their strong expression of *Pcolce2*/procollagen C-endopeptidase enhance 2 (**Fig. 1D**), endoneurial mesenchymal cells (eMES1-eMES3, clusters 10-12), expressing the tumor suppressor *Cdkn2a*/cyclin-dependent kinase inhibitor 2A (**Fig. 1E**), and perineurial (p)MES in clusters 13 and 14, expressing the tight junction molecule *Cldn1*/claudin 1 (**Fig. 1F**).

**Table. S1.**
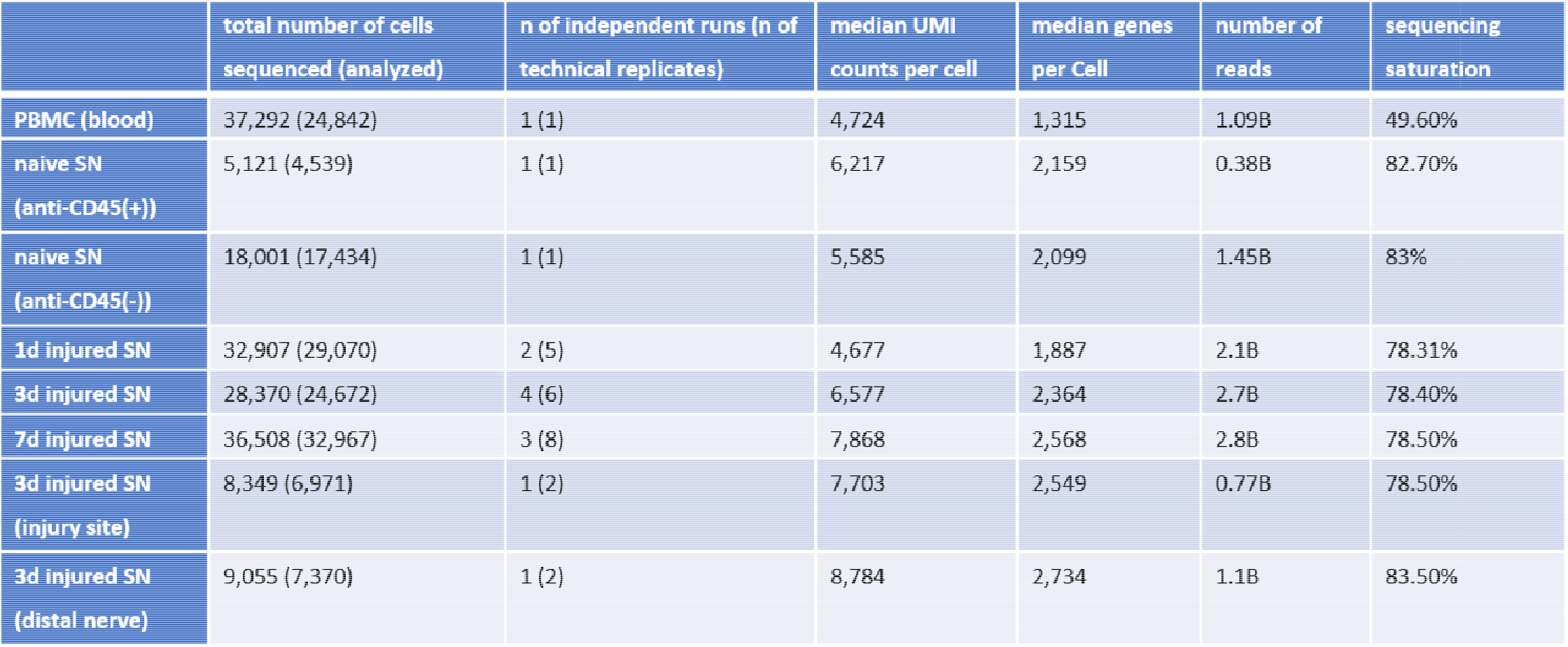
Summary of scRNAseq datasets generated and analyzed in this study. A total of 147,865 high-quality single cell transcriptomes were generated from naïve mouse sciatic nerve, injured sciatic nerve, and peripheral blood mononuclear cells (PBMC). Some of the 3-day (3d) injured nerves were divided into injury site and distal nerve and sequenced separately. The number of independent runs, biological replicates, is shown and defined as cells sequenced from different cohorts of mice. The number of technical replicates is shown in parenthesis and indicates how many times cells from the same cohort of mice was sequenced.

**Figure 1.**
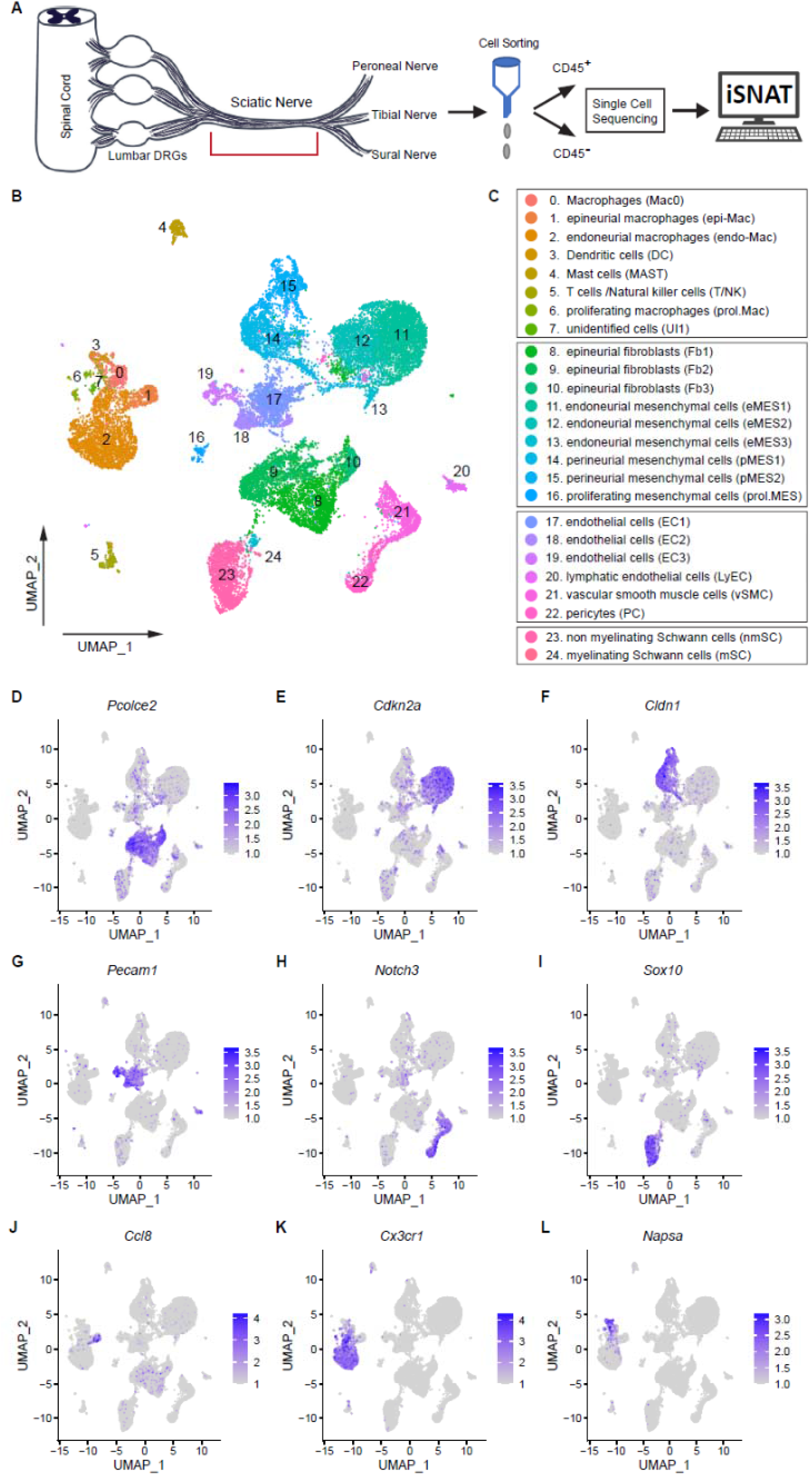
The cellular and molecular landscape of naïve mouse sciatic nerve. **(A)** Workflow for peripheral nervous tissue analysis. Cartoon of a mouse lumbar spinal cord with dorsal root ganglia (DRGs), sciatic nerve trunk and major branches. The sciatic nerve trunk is marked with a red bracket and was harvested for further analysis. Immune cells were captured using anti-CD45 magnetic beads. The flow through containing non-immune (CD45^-^) cells collected. In separate scRNAseq runs, CD45^+^ and CD45^-^ single-cell transcriptomes were determined. A total of 21,973 high-quality transcriptomes, including 4,539 CD45^+^ cells and 17,434 CD45^-^ cells were generated and used for downstream analysis. **(B)** UMAP embedding of naïve sciatic nerve cells. Unsupervised Seurat-based clustering identified 24 clusters. **(C)** List of cell types identified in the naïve sciatic nerve, grouped into immune cells (clusters 0-7), structural/stromal cells (clusters 8-16), cells associated with the nerve vasculature (clusters 17-22), and Schwann cells (clusters 23 and 24). **(D-L)** Feature plots of canonical markers used to identify major cell types, including epineurial fibroblasts (*Pcolce2/*procollagen C-endopeptidase enhancer 2), endoneurial MES (*Cdkn2a/*cyclin dependent kinase inhibitor 2A), perineurial MES (*Cldn1/*claudin-1), endothelial cells (*Pecam1*/CD31), vascular smooth muscle cells and pericytes (*Notch3/*notch receptor 3), Schwann cells (*Sox10/*SRY-box transcription factor 10), epineurial Mac (*Ccl8/*C-C motif chemokine ligand 8), endoneurial Mac (*Cx3cr1/*C-X3-C motif chemokine receptor 1), and dendritic cells (*Napsa/*napsin A aspartic peptidase). Expression levels are color coded and calibrated to average gene expression.

The nerve vasculature is comprised of three clusters with endothelial cells (EC1-EC3, clusters 16-18), strongly expressing *Pecam1/*CD31 (**Fig. 1G**). EC express high levels of adherent junction (*Cldn5*/claudin 5), tight junction (*Tjp1*), and junctional adhesion molecules (*Jam2*), key components of the BNB (**Fig. 1, Suppl. 1A**) (Maiuolo et al., 2019). Clusters 20 and 21 are connected and harbor *Notch3*^+^ cells (**Fig. 1H**) that can be further subdivided into vascular smooth muscle cells (vSMC, *Acta2^+^*) and pericytes (PC, *Pdgfrb^+^*) (**Fig. 1, Suppl. 1A**). A small island (cluster 19) with lymphatic endothelial cells (LyEC, *Prox1*) was detected (**Fig. 1, Suppl. 1A**). Schwann cells (SC, *Sox10*) are mainly comprised of non-myelinating (nm)SC (cluster 22) and few myelinating (m)SC (cluster 23) (**Fig. 1I**) (Gerber et al., 2021). The low number of mSC indicates that many are lost during the cell isolation process from naïve nerve (**Fig. 1, Suppl. 1A**).

**Figure 1, Suppl. 1.**
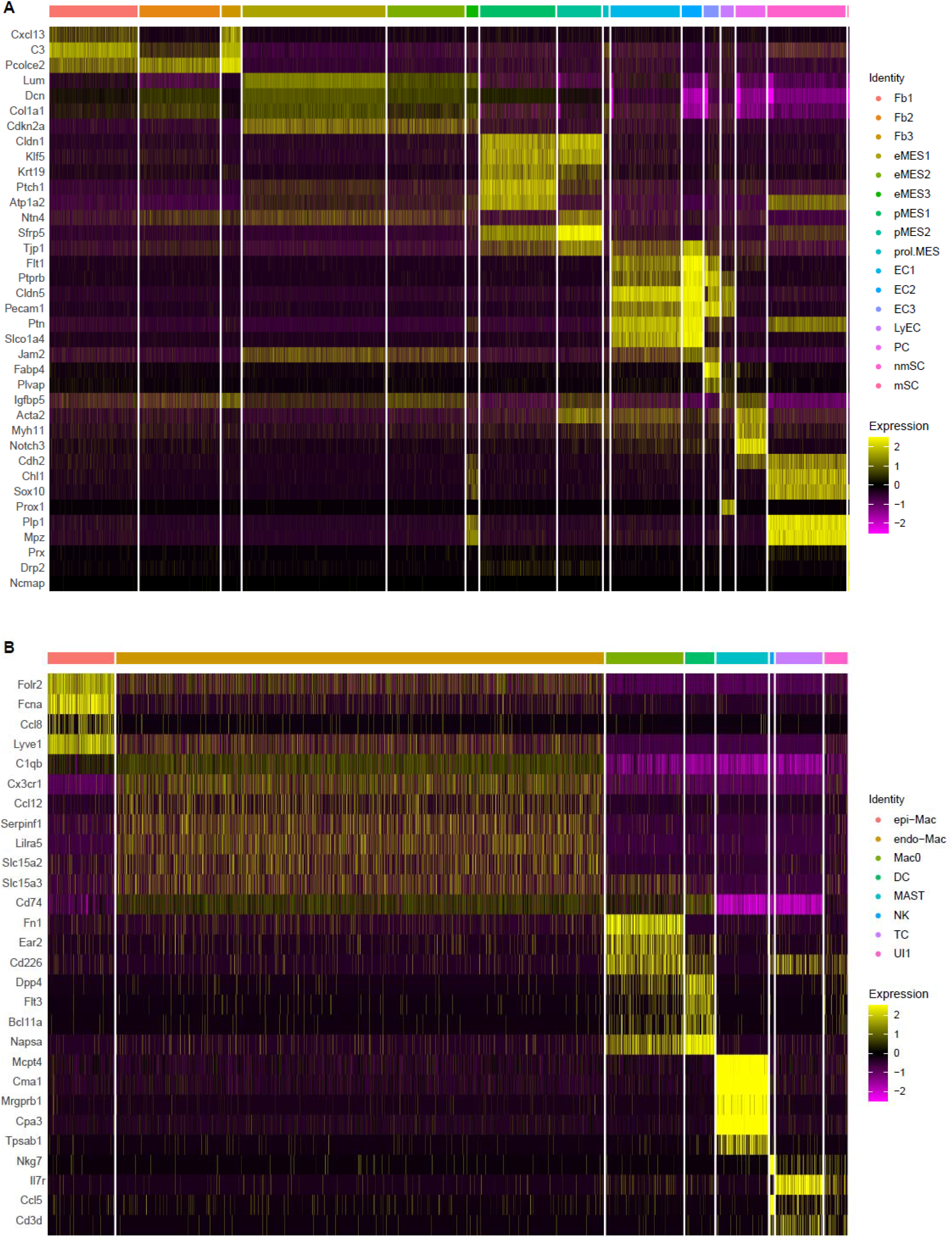
Heatmaps of top cluster enriched gene products in the naïve mouse sciatic nerve trunk for **(A)** non-immune cells, and **(B)** immune cells. Expression levels are calibrated to median gene expression. Abbreviations: Fb, fibroblasts; eMES, endoneurial mesenchymal cells; pMES, perineurial mesenchymal cells; prol.MES, proliferating mesenchymal cells; EC, endothelial cells; LyEC, lymphatic endothelial cells; PC, pericytes; nmSC, non-myelinating Schwann cells; mSC, myelinating Schwann cells; epiMac, epineurial macrophages; endoMac, endoneurial macrophages; DC, dendritic cells; MAST, mast cells; NK, natural killer cells; TC, T cells; UI, unidentified cells.

Mac (clusters 0, 1, and 2) are the most abundant immune cell type in the naïve nerve, representing 75% of all immune cells. Smaller immune cell clusters with dendritic cells (DC, 3%), mast cells (Mast, 6%), T cells (TC) and Natural killer cells (NK) 5% are detected (**Fig. 1B, Suppl. 1B**). Less than 1% of immune cells express markers for granulocytes (GC, *Cxcr2*/C-X-C motif chemokine receptor 2), monocytes (Mo, *Chil3*/chitinase-like protein 3), or B cells (BC, *Cd79a/*B cell antigen receptor-associated protein), indicating these cell types are sparse in the naïve PNS of healthy mice. Tissue resident Mac in the naïve PNS strongly express the scavenger receptor SCARI1, encoded by *Cd163*. A small group of *Fn1*(fibronectin) expressing Mac (cluster 0) was detected and these may represent leukocytes that recently entered the nerve (**Fig. 1, Suppl. 2A, 2B**). Previous scRNA-seq studies employed cell sorting to isolate Mac from naïve PNS, using CD45^+^CD64^+^F4/80^+^ (Ydens et al., 2020) or CD45^int^CD64^+^ sorting (Wang et al., 2020), and reported transcriptionally distinct epineurial (epi) Mac and endoneurial (endo) Mac. When compared to our dataset, we find that cells in cluster 1 express *Fcna*/Ficolin-A and *Ccl8/*monocyte chemotactic protein 2, and thus, represent epiMac (**Fig. 1J** and **Fig. 1, Suppl. 2C**). The expression of *Ccl8* by epiMac was validated by *in situ* hybridization using RNAscope. *Ccl8* labeling was highest in Mac located in the epineurium (**Fig. 1, Suppl. 2G,G’**). Cells in cluster 2 preferentially express *Cx3cr1, Trem2, Lilra5*, and the small peptide transporter *Slc15a2* and represent endoMac (**Fig. 1K** and **Fig. 1 Suppl. 2D-2F**). Longitudinal sciatic nerve sections of *Cx3cr1-GFP* reporter mice confirmed labeling of endoMac (**Fig. 1, Suppl 2H,H’)**. Mac are educated by the local nerve microenvironment and acquire niche specific phenotypes. In the naïve PNS, epiMac (*Lyve1^hi^, Cx3cr1^lo^*) are located in the heavily vascularized epineurium and resemble perivascular Mac (Chakarov et al., 2019), while endoMac exhibit neural niche gene signatures reminiscent of Mac in the enteric plexus (De Schepper et al., 2019) and microglia (Wang et al., 2020). There are few DC in the naïve nerve, they function as professional antigen presenting cells and can be identified by their strong expression of *Napsa/*aspartic peptidase (**Fig. 1L**). Mast cells in the naïve nerve are readily identified by their preferential expression of *Fcer1a* (the immunoglobulin epsilon receptor for IgE) and *Cpa3* (carboxypeptidase A) (**Fig. 1, Suppl 1B**).

**Figure 1, Suppl. 2.**
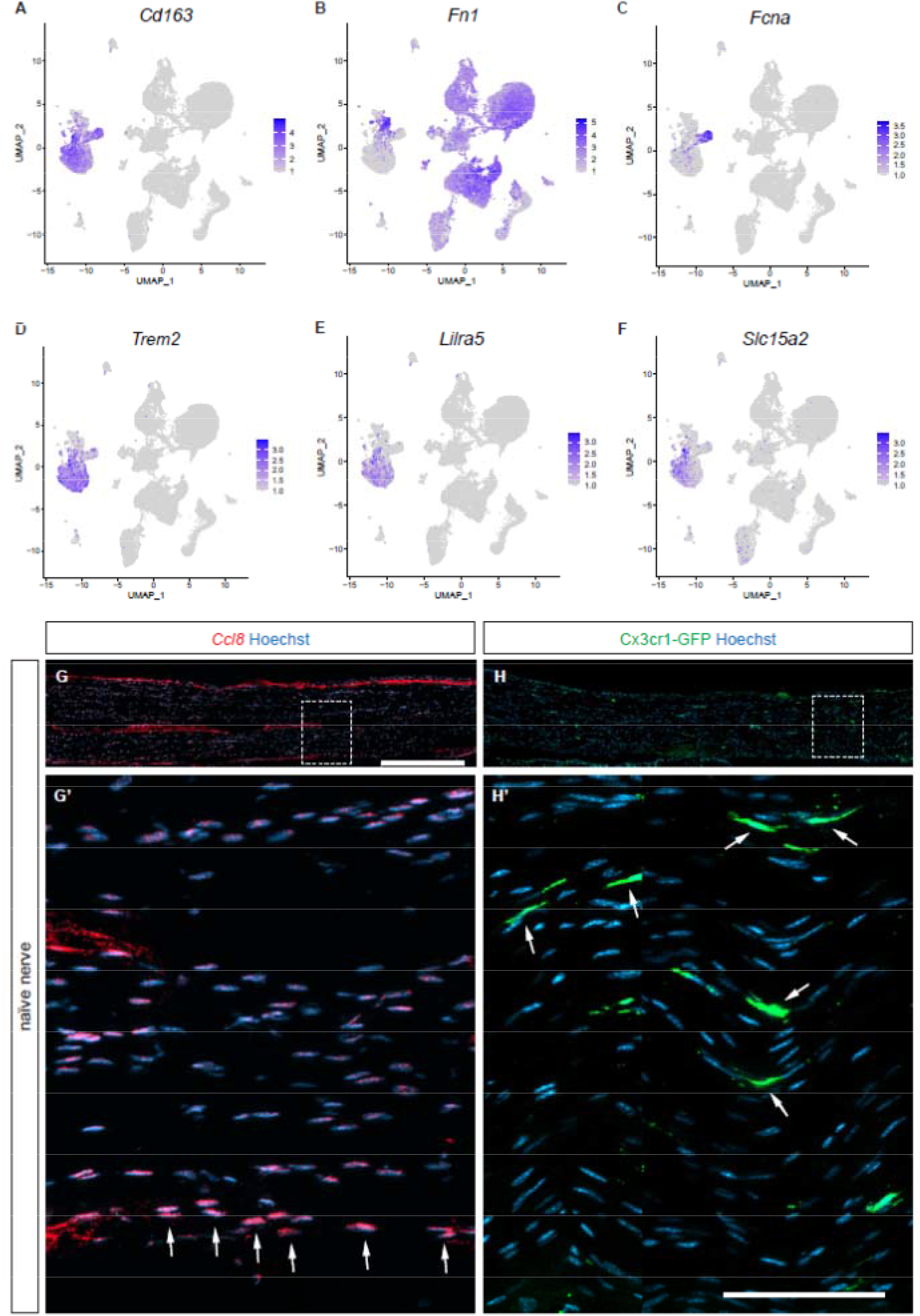
**(A-F)** Feature plots of naïve sciatic nerve. (**A**) *Cd163* is expressed by epi-Mac and endo-Mac. (**B**) A small subpopulation of Mac, expressing very high levels of *Fn1* (fibronectin), is observed in the naïve nerve. In addition, *Fn1* is expressed by most non-immune cells. (**C**) *Fcna* (Ficolin A) is strongly and selectively expressed by epi-Mac. (**D-F**) Show expression of the endo-Mac markers *Trem2* (triggering receptor expressed on myeloid cells 2), *Lilra5* (leukocyte immunoglobulin like receptor A5), and *Slc15a1* (solute carrier family 15 member 1). Expression levels are projected onto the UMAP with a minimum expression cutoff of 1. **(G,G’)** Longitudinal sections of naïve mouse sciatic nerve. *In situ* hybridization of *Ccl8* with RNAscope revealed preferential staining of epi-Mac, labeled with arrows. (**H,H’**) Longitudinal sciatic nerve section of *Cx3cr1-GFP* reporter mice revealed preferential labeling of endo-Mac. Scale bar (**G,H**), 500 μm; (**G’,H’**),100 μm.

To independently validate naïve nerve scRNAseq datasets, we examined the presence of some of the corresponding proteins in naïve sciatic nerve lysates by ELISA. We used an ELISA kit that allows simultaneous profiling of 111 extracellular proteins, including cytokines, chemokines, growth factors, proteases, and protease inhibitors (**Fig. 1, Suppl. 3A**). Proteins abundantly detected include, FGF1 (*Fgf1*), IGF binding protein 6 (*Igfbp6*), the anti-angiogenic factor endostatin (*Col18a1*), interleukin 33 (*ll33*), coagulation factor III (*F3*), cystatin C (*Cst3*), retinoic acid binding protein 4 (*Rbp4*), matrix metallopeptidase 2 (*Mmp2*), intercellular adhesion molecule 1 (*Icam1*), fetuin A/alpha 2-HS glycoprotein (*Ahsg*), resistin/adipose tissue-specific secretory factor (*Retn*), low-density lipoprotein receptor (*Ldlr*), serpin F1/neurotrophic protein (*Serpinf1*), and adiponectin (*Adipoq*), a regulator of fat metabolism (**Fig. 1, Suppl. 3B**). For many of these proteins, the corresponding transcripts are present in our scRNAseq data (**Fig. 1, Suppl. 3C**). Gene products abundantly detected by ELISA, but not by scRNAseq, include fetuin A, resistin, and adiponectin (**Fig. 1, Suppl. 3C**). Resistin and adiponectin are both secreted by adipocytes, and fetuin A is secreted by hepatocytes. All three proteins are known serum components, and thus, may have entered the nerve via the circulatory system. Indeed, ELISA for serum proteins from naïve mice, revealed high levels of fetuin A, resistin, and adiponectin (**Fig. 1, Suppl. 3D**). Importantly, some of the most abundant serum proteins, including E-selectin (*Sele*), IGFBPs (*Igfbp1, 2, 3* and *5*), proprotein convertase 9 (*Pcsk1*), soluble C1qR1 (*Cd93)*, angiopoietin-2 (*Angpt2*), C-C motif chemokine ligand 21 (*Ccl21*), and CRP/C-reactive protein (*Crp*) are not present in nerve lysates, providing confidence that mice were perfused properly, and nerve samples not contaminated with serum (**Fig. 1, Suppl. 3E**). These experiments also identify serum proteins that can or cannot cross the intact BNB. Collectively, our analysis establishes a baseline of the cellular landscape and molecular milieu of naïve PNS. These data will be used as reference to determine how a crush injury alters the nerve microenvironment.

**Figure 1, Suppl. 3.**
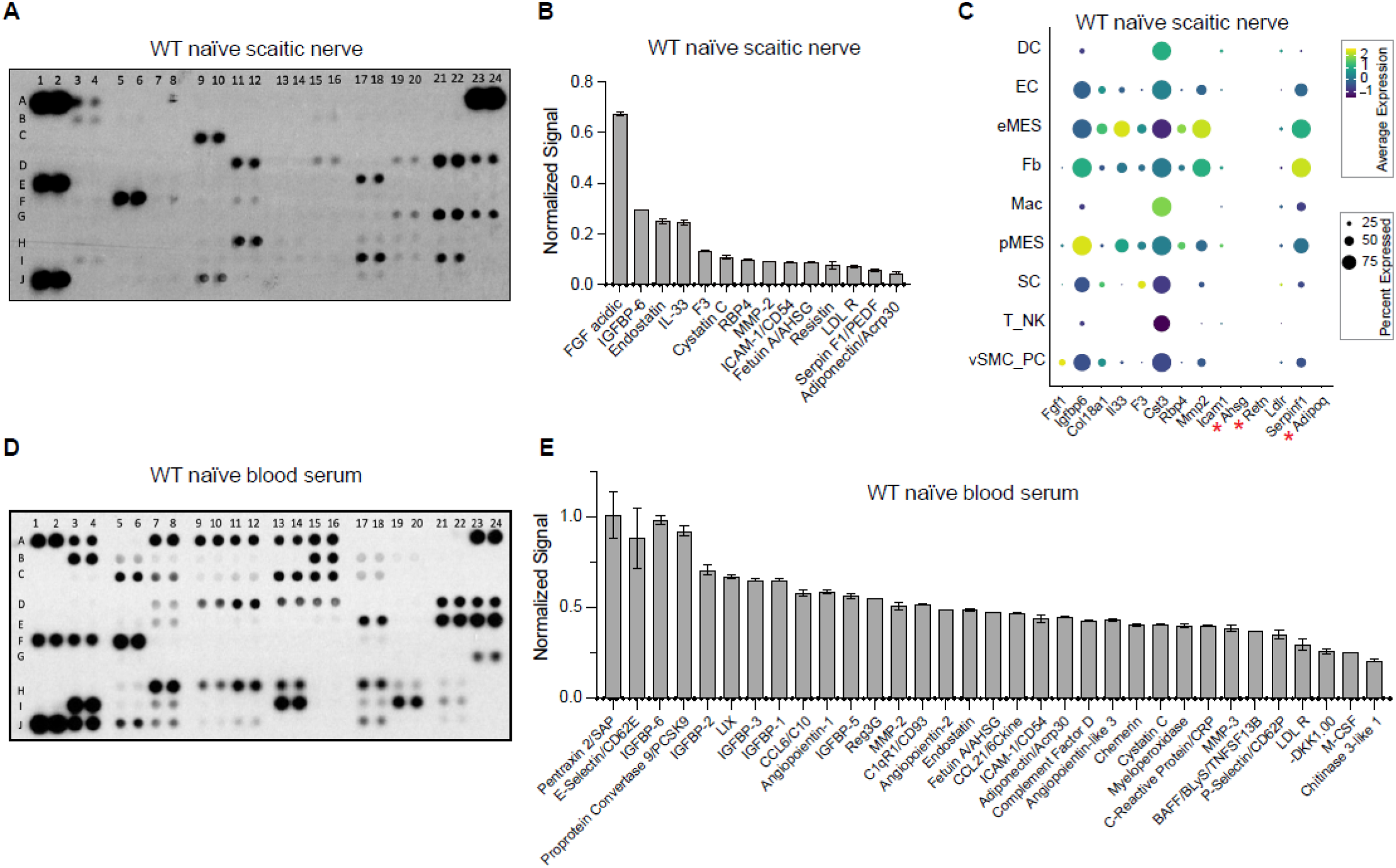
**(A)** ELISA membrane probed with naïve sciatic nerve lysate. **(B)** List of top proteins detected. For quantification, ELISA signals were normalized to reference spots shown at coordinates (A1,A2), (A23,A24), and (J1,J2). **(C)** Dotplot analysis of the corresponding gene products of naïve sciatic nerve cells, as identified by scRNAseq. Relative expression levels, normalized to average gene expression (color coded) are shown. For each cell cluster, the percentile of cells expressing a specific gene product is indicated by the dot size. **(D)** ELISA membrane probed with serum from naïve mice. (**E**) List of top serum proteins detected. For quantification, ELISA signals were normalized to reference spots shown at coordinates (A1,A2), (A23,A24), and (J1,J2). Number of biological replicates, n =1. Coordinates of proteins that can be detected by the ELISA, (A1, A2) Reference spots, (A3, A4) Adiponectin [*Adipoq*], (A5, A6) Amphiregulin [*Areg*], (A7, A8) Angiopoientin-1 [*Angpt1*], (A9, A10) Angiopoientin-2 [*Angpt2*], (A11, A12) Angiopoientin-like 3 [*Angptl3*], (A13, A14) BAFF [*Tnfrsf13b*], (A15, A16) C1qR1 [*Cd93*], (A17, A18) CCL2 [*Ccl2*], (A19, A20) CCL3 [*Ccl3*], (A21, A22) CCL5 [*Ccl5*], (A23, A24) Reference spots, (B3, B4) CCL6 [*Ccl6*], (B5, B6) CCL11 [*Ccl11*], (B7, B8) CCL12 [*Ccl12*], (B9, B10) CCL17 [*Ccl17*], (B11, B12) CCL19 [*Ccl19*], (B13, B14) CCL20 [*Ccl20*], (B15, B16) CCL21 [*Ccl21*], (B17, B18) CCL22 [*Ccl22*], (B19, B20) CD14 [*Cd14*], (B21, B22) CD40 [*Cd40*], (C3, C4) CD160 [*Cd160*], (C5, C6) Chemerin [*Rarres2*], (C7, C8) Chitinase 3-like 1 [*Chil3*], (C9, C10) Coagulation Factor III [*F3*], (C11, C12) Complement Component C5 [*C5*], (C13, C14) Complement Factor D [*Cfd*], (C15, C16) C-Reactive Protein [*Crp*], (C17, C18) CX3CL1 [*Cx3cl1*], (C19, C20) CXCL1 [*Cxcl1*], (C21, C22) CXCL2 [*Cxcl2*], (D1, D2) CXCL9 [*Cxcl9*], (D3, D4) CXCL10 [*Cxcl10*], (D5, D6) CXCL11 [*Cxcl11*], (D7, D8) CXCL13 [*Cxcl13*], (D9, D10) CXCL16 [*Cxcl16*], (D11, D12) Cystatin C [*Cst3*], (D13, D14) DKK-1 [*Dkk1*], (D15, D16) DPPIV [*Dpp4*], (D17, D18) EGF [*Egf*], (D19, D20) Endoglin [*Eng*], (D21, D22) Endostatin [*Col18a1*], (D23, D24) Fetuin A [*Ahsg*], (E1, E2) FGF acidic [*Fgf1*], (E3, E4) FGF-21 [*Fgf21*], (E5, E6) Flt-3 Ligand [*Flt3l*], (E7, E8) Gas 6 [*Gas6*], (E9, E10) G-CSF [*Csf3*], (E11, E12) GDF-15 [*Gdf15*], (E13, E14) GM-CSF [*Csf2*], (E15, E16) HGF [*Hgf*], (E17, E18) ICAM-1 [*Icam1*], (E19, E20) IFN-gamma [*Ifng*], (E21, E22) IGFBP-1 [*Igfbp1*], (E23, E24) IGFBP-2 [*Igfbp2*], (F1, F2) IGFBP-3 [*Igfbp3*], (F3, F4) IGFBP-5 [*Igfbp5*], (F5, F6) IGFBP-6 [*Igfbp6*], (F7, F8) IL-1alpha [*Il1a*], (F9, F10) IL-1Beta [*Il1b*], (F11, F12) IL-1ra [*Il1rn*], (F13, F14) IL-2 [*Il2*], (F15, F16) IL-3 [*Il3*], (F17, F18) IL-4 [*Il4*], (F19, F20) IL-5 [*Il5*], (F21, F22) IL-6 [*Il6*], (F23, F24) IL-7 [*Il7*], (G1, G2) IL-10 [*Il10*], (G3, G4) IL-11 [*Il11*], (G5, G6) IL-12 p40 [*Il12*], (G7, G8) IL-13 [*Il13*], (G9, G10) IL-15 [*Il15*], (G11, G12) IL-17A [*Il17a*], (G13, G14) IL-22 [*Il22*], (G15, G16) IL-23 [*Il23*], (G17, G18) IL-27 p28 [*Il27*], (G19, G20) IL-28 [*Ifnl3*], (G21, G22) IL-33 [*Il33*], (G23, G24) LDL R [*Ldlr*], (H1, H2) Leptin [*Lep*], (H3, H4) LIF [*Lif*], (H5, H6) Lipocalin-2 [*Lcn2*], (H7, H8) LIX [*Cxcl5*], (H9, H10) M-CSF [*Csf1*], (H11, H12) MMP-2 [*Mmp2*], (H13, H14) MMP-3 [*Mmp3*], (H15, H16) MMP-9 [*Mmp9*], (H17, H18) Myeloperoxidase [*Mpo*], (H19, H20) Osteopontin [*Spp1*], (H21, H22) Osteoprotegerin [*Tnfrsf11b*], (H23, H24) PD-ECGF [*Tymp*], (I1, I2) PDGF-BB [*Pdgfb*], (I3, I4) Pentraxin 2 [*Nptx2*], (I5, I6) Pentraxin 3 [*Ptx3*], (I7, I8) Periostin [*Postn*], (I9, I10) Pref-1 [*Dlk1*], (I11, I12) Proliferin [*Prl2c2*], (I13, I14) Proprotein Convertase 9 [*Pcsk9*], (I15, I16) RAGE [*Ager*], (I17, I18) RBP4 [*Rbp4*], (I19, I20) Reg3G [*Reg3g*], (I21, I22) Resistin [*Retn*], (J1, J2) Reference spots, (J3, J4) E-Selectin [*Sele*], (J5, J6) P-Selectin [*Selp*], (J7, J8) Serpin E1 [*Serpine1*], (J9, J10) Serpin F1 [*Serpinf1*], (J11, J12) Thrombopoietin [*Thpo*], (J13, J14) TIM-1 [*Havcr1*], (J15, J16) TNF-alpha [*Tnf*], (J17, J18) VCAM-1 [*Vcam1*], (J19, J20) VEGF [*Vegf*], (J21, J22) WISP-1 [*Ccn4*], (J23, J24) negative control.

### The injured sciatic nerve atlas (iSNAT)

To describe the dynamic nature of the cellular and molecular landscape of injured PNS tissue, we subjected adult mice to sciatic nerve crush injury (SNC) and carried out scRNAseq at 1-day post-SNC (1dpc), 3dpc, and 7dpc (**Fig. 2A)**. **Table S1** shows the number of cells analyzed at each time point. The longitudinal study revealed marker genes that allow reliable identification of major cell types in the naïve and injured PNS (**Fig. 2, Suppl. 1A-D**). SNC triggers massive infiltration of blood-borne immune cells (Kalinski et al., 2020; Ydens et al., 2020). In the 1dpc UMAP plot we identified granulocytes (GC, *Cxcr2*) in cluster 11 (Fig. 2D), Mo (*Ly6c2/* lymphocyte antigen 6C2 and *Chil3/*Ym1) in cluster 0 (**Fig. 2F, Fig. 2, Suppl. 1B**), and five Mac subpopulations (Mac-I to Mac-V) in clusters 1-5, expressing *Adgre1*(F4/80) (**Fig. 2G**). GC are primarily composed of neutrophils (*Retnlg/*resistin-like gamma*, Grina/*NMDA receptor associated protein 1), intermingled with a smaller population of eosinophils (*Siglecf*) (**Fig. 2E**). Mo that recently entered the nerve can be identified by high levels of *Cd177* (data not shown) a known surface molecule that interacts with the EC adhesion molecule *Pecam1*/CD31 and promotes transmigration. Mo strongly express *Ly6c2*, suggesting they represent Ly6C^hi^ proinflammatory cells. In the UMAP plot of 1dpc nerves, cluster 0 (Mo) has a volcano-like shape connected at the top to cluster 1, harboring Mac-I (*Ccr2^hi^, Ly6c2^low^, Cx3cr1*^-^), indicating these are maturing monocyte-derived Mac (**Fig. 2B**). Four smaller Mac clusters include, Mac-II (*C1qa*, complement factor C1q), Mac-III (*Ltc4s*/leukotriene C4 synthase, *Mmp1*9/matrix metallopeptidase 19), Mac-IV expressing MHCII genes (*H2-Aa, H2-Eb1, H2-Ab1, Cd74*), and Mac-V (*Il1rn/*IL1 receptor antagonist, *Cd36/*ECM receptor, coagulation factors *F7* and *F10*), suggesting roles in opsonization, matrix remodeling, antigen presentation, and stopping of bleeding (**Fig. 2B**, and data not shown). Cluster 6 harbors monocyte-derived dendritic cells (MoDC: *Cd209a*/DC-SIGN, *H2-Aa*/MHC class II antigen A, alpha), professional antigen presenting cells (**Fig. 2H, 2I**). A small island (cluster 13) harbors a mixture of TC and NK (**Fig. 2B**). The 1dpc dataset contains five clusters with structural cells (Fb, differentiating (d)MES, eMES, and pMES), including a small number of proliferating (*Mki67^+^/*Ki67) cells, vasculature cells, EC (cluster 20), vSMC and PC (cluster 21) (**Fig. 2B and Fig. 2, Suppl. 2B**).

**Figure 2.**
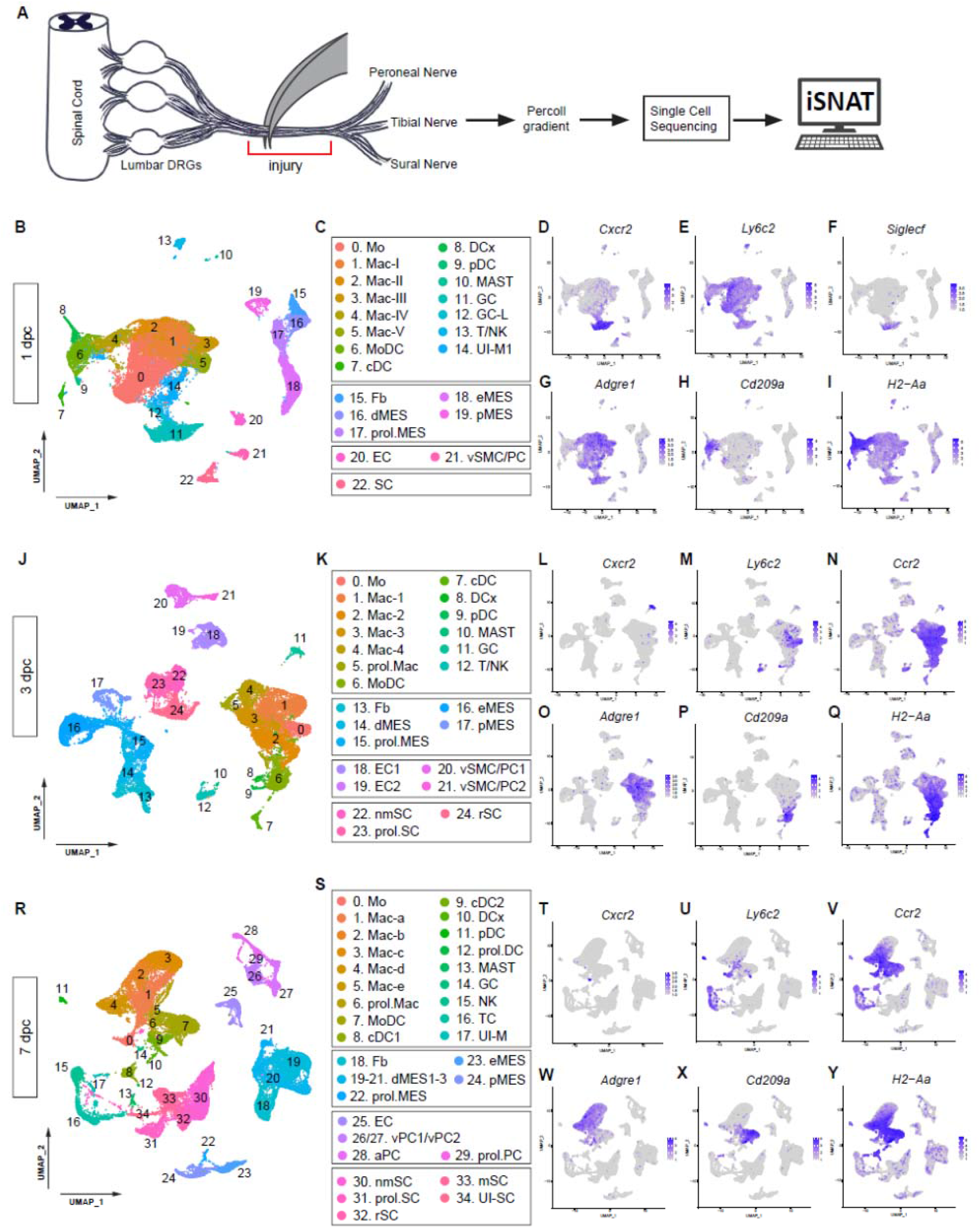
Longitudinal analysis of single-cell transcriptomes of injured PNS. **(A)** Workflow for single cell analysis of injured mouse sciatic nerve trunk. Cartoon of lumbar spinal cord with DRGs and sciatic nerve. The injury site is shown and the segment marked with the red bracket was harvested at different post-injury time points and analyzed by scRNAseq. **(B)** UMAP plot embedding of sciatic nerve cells at 1dpc. A total of 29,070 high-quality cells (n= 2 biological replicates, 5 technical replates) were subjected to unsupervised Seurat-based clustering resulting in 22 cell clusters. **(C)** List of cell types identified, grouped into immune cells (clusters 0-14), structural cells (clusters 15-19), cells associated with the nerve vasculature (clusters 20 and 21), and Schwann cells (cluster 22). **(D-I)** Feature plots of canonical immune cell markers to identify clusters with GC (*Cxcr2*), including a subset of eosinophils (*Siglecf/*sialic acid binding Ig-like lectin F), Mo (*Ly6c2/*Ly6C), Mac (*Adgre1/*F4/80), MoDC (*Cd209a/*DC-SIGN), and antigen presenting cells (*H2-Aa*/histocompatibility 2, class II antigen A, alpha). **(J)** UMAP plot embedding of sciatic nerve cells at 3dpc. A total of 24,672 high-quality cells (n= 4 biological replicates, 6 technical replates) were subjected to unsupervised Seurat-based clustering resulting in 24 cell clusters. **(K)** List of cell types in the 3-day injured nerve, grouped into immune cells (clusters 0-12), structural cells (clusters 13-17), cells associated with the nerve vasculature (clusters 18-21), and Schwann cells (clusters 22-24). **(L-Q)** Feature plots of canonical markers for immune cells to identify clusters with GC (*Cxcr2*), Mo (*Ly6c2*), Mo/Mac (*Ccr2*), Mac (*Adgre1*), MoDC (*Cd209a*), and antigen presenting cells (*H2-Aa*). **(R)** UMAP plot embedding of sciatic nerve cells at 7dpc. A total of 32,976 high-quality cells (n= 3 biological replicates, 8 technical replates) were subjected to unsupervised Seurat-based clustering resulting in 34 cell clusters. **(S)** List of cell types in the nerve at 7dpc, grouped into immune cells (clusters 0-17), structural cells (clusters 18-24), cells associated with the nerve vasculature (clusters 25-29), and Schwann cells (clusters 30-34). **(T-Y)** Feature plots of canonical markers for immune cells to identify clusters with GC (*Cxcr2*), Mo (*Ly6c2*), Mo/Mac (*Ccr2*), Mac (*Adgre1*), MoDC (*Cd209a*), and antigen presenting cells (*H2-Aa*). Expression levels are projected onto the UMAP with a minimum expression cutoff of 1. Abbreviations for immune cells: Mo, monocytes; Mac, macrophages; prol.Mac, proliferating macrophages; MoDC, monocyte-derived dendritic cells; cDC, conventional dendritic cells; DCx mature/migrating dendritic cells; pDC, plasmacytoid dendritic cells; MAST, mast cells; GC, granulocytes (including neutrophils and eosinophils), GC-L, granule cell-like; TC, T cells; NK, natural killer cells. Abbreviations for structural cells: Fb, fibroblast; dMES, differentiating mesenchymal cells; prol.MES, proliferating mesenchymal cells; eMES, endoneurial mesenchymal cells; pMES, perineurial mesenchymal cells. Abbreviations for vascular cells: EC, endothelial cells, vSMC, vascular smooth muscle cells; PC, pericytes; vPC, venous pericytes; aPC, arterial pericytes; prol.PC, proliferating pericytes. Abbreviations Schwann cells, nmSC, non-myelinating Schwann cells; mSC myelinating Schwann cells; rSC, repair Schwann cells; prol.SC, proliferating Schwann cells. UI, unidentified cells.

**Figure 2, Suppl Figure 1.**
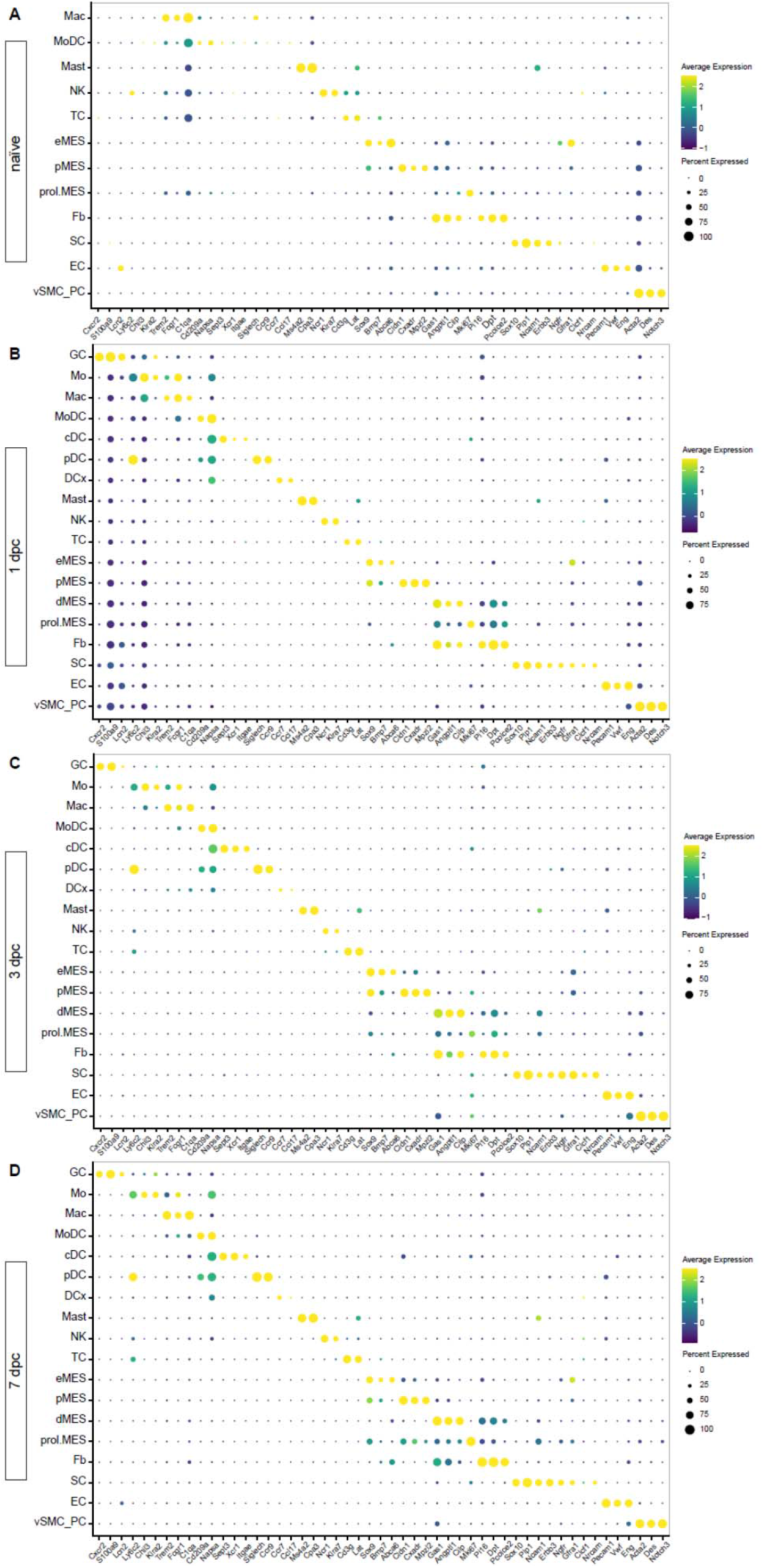
Identification of marker genes for cell type identification in the naïve and injured PNS. Dotplot analysis of scRNAseq datasets of **(A)** naïve nerve, **(B)** 1dpc nerve, **(C)** 3dpc nerve, and **(D)** 7dpc nerve. Expression levels are normalized to average gene expression (color coded). For each cell cluster, the percentile of cells expressing a specific gene product is indicated by the dot size.

**Figure 2, Suppl Figure 2.**
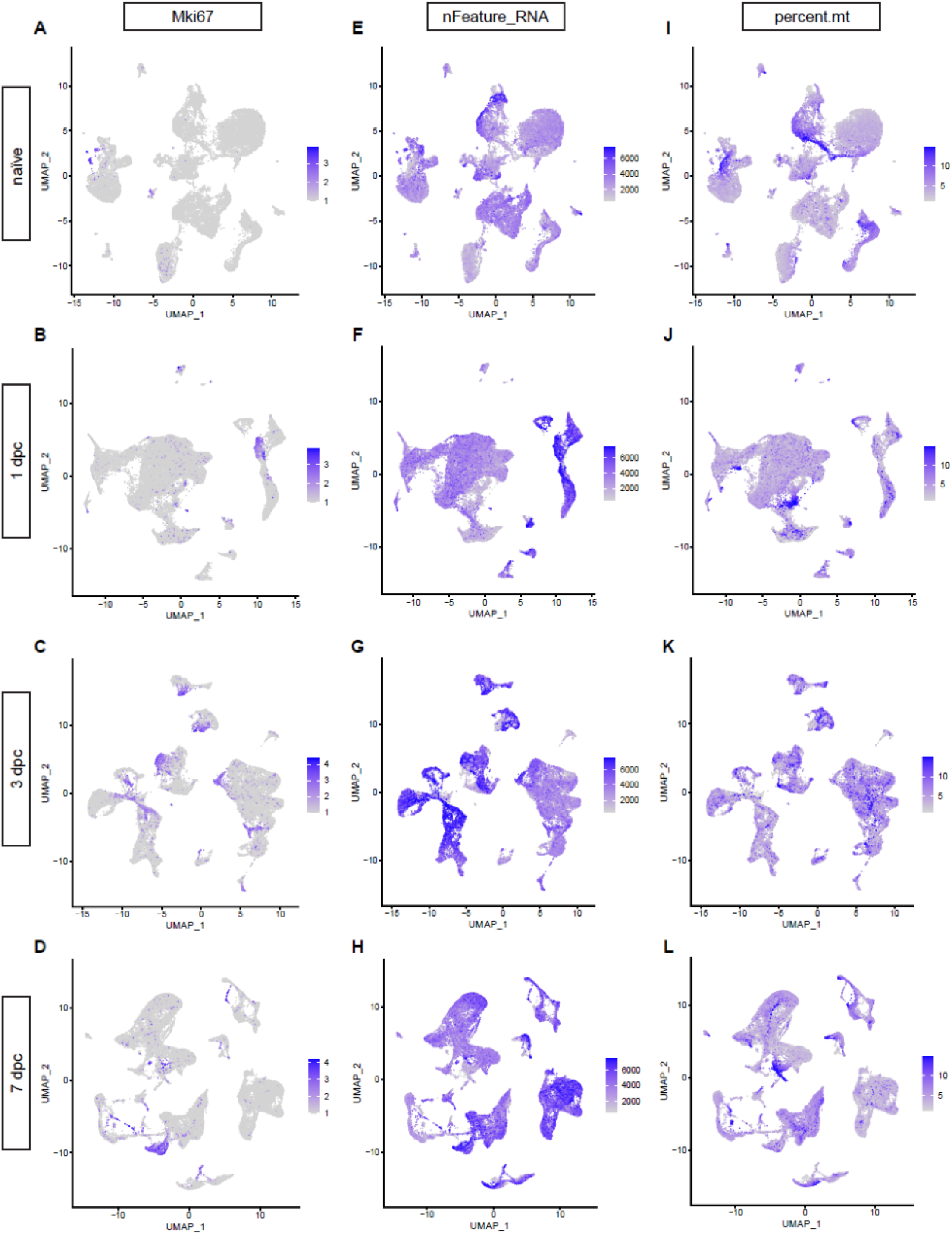
Feature plots for *Mki67* reveals proliferating cells in **(A)** naïve nerve, **(B)** 1dpc, **(C)** 3dpc, and **(D)** 7dpc. The expression levels are projected onto the UMAP with a minimum expression cutoff of 1. **(E-H)** Feature plot showing the number of unique transcripts detected in naïve and injured sciatic nerve cells. Color coded calibration is shown. Note, cells with less than 500 unique features or more than 7500 were excluded from the study. **(I-L)** Feature plots showing the mitochondrial content of cells in the naïve and injured nerves. Color coded calibration is shown. Note, cells with more than 15% mitochondrial content were excluded from the analysis.

In the UMAP plot of 3dpc nerve (**Fig. 2J**), far fewer GC (*Cxcr2*) (**Fig. 2L**) and Mo (*Ly6c2*) (**Fig. 2M**) are present when compared to 1dpc nerve. *Adgre1*^+^ Mac are the most dominant immune cell type at 3dpc, comprised of five subpopulations, (clusters 1-5) designated Mac1-Mac4, and prol. Mac (**Fig. 2O**). *Ccr2* labels most Mac and DC, except for Mac4 and a subset of Mac1 cells (**Fig. 2N**). As discussed below, there is no one-to-one match of Mac subclusters identified in the 1dpc and 3dpc nerve, likely owing to their high degree of transcriptional plasticity. To emphasize this observation, we labeled Mac in the 1dpc nerve as MacI-MacV and Mac in the 3dpc nerve as Mac1-Mac4 and prol.Mac (**Fig. 2C and 2K**). MoDC (*Cd209a*) are readily detected in the 3dpc nerve (**Fig. 2P**). Compared to the 1dpc nerve, the fraction of MHCII^+^ (*H2-Aa*) cells is increased, including four subpopulations of dendritic cells (MoDC, cDC, DCx, and pDC, see below for details), as well as subpopulations of Mac (**Fig. 2Q**). Structural cells begin to proliferate heavily at 3dpc, suggesting that a nerve crush injury causes substantial damage to the epineurium, perineurium, and endoneurium (**Fig. 2, Suppl. 2C**). SNC is known to inflict vascular damage and breach in the BNB. This is underscored by the strong proliferative response of vascular cells, including EC, vSMC, and PC (**Fig. 2, Suppl. 2C**).

In the 7dpc UMAP plot (**Fig. 2R**), GC and Mo have further declined (**Fig. 2T, 2U**), and *Adgre1^+^* Mac remain highly prevalent (**Fig. 2W**). Mac form a connected continuum of clusters, harboring subpopulations Mac-a to Mac-d (**Fig. 2S**). *Ccr2* remains high in most Mac subpopulations, except for Mac-c and is also observed in Mo and DC (**Fig. 2V**). Similar to earlier timepoints, there is no clear one-to-one match of Mac clusters at 7dpc to Mac at 3dpc and are thus labeled Mac-a to Mac-e (**Fig. 2R**). At 7dpc, the number of *Cd209a^+^*DC has increased compared to earlier time points (**Fig. 2X**). MHCII (*H2-Aa*) expressing Mac and DC are readily detected (**Fig. 2Y**). Lymphoid cells, identified by *Il2rb*/IL2 receptor subunit β expression, include NK (cluster 15) and TC (cluster 16) (**Fig. 2R**). At 7dpc, the proliferation of structural and vascular cells is reduced compared to 3dpc (**Fig. 2, Suppl. 2C, 2D**), suggesting that cells required for the repair process and wound healing are in place.

The sequelae of successful tissue repair requires extensive communication between nerve resident cells and hematogenous immune cells. For a detailed description of the cellular and molecular changes that occur during the first week following PNS injury, we generated and analyzed more than 180,000 high-quality single-cell transcriptomes (**Table S1**). These datasets provide a new resource for understanding cell function and molecules in the degenerating and regenerating mammalian PNS. For widespread distribution of scRNAseq data, we created the injured sciatic nerve atlas (iSNAT), a user-friendly and freely available web-based resource. The “*Expression Analysis*” function in iSNAT provides a readily accessible platform for comparative analysis of gene expression in naïve and injured nerve at single-cell resolution, https://cdb-rshiny.med.umich.edu/Giger_iSNAT/.

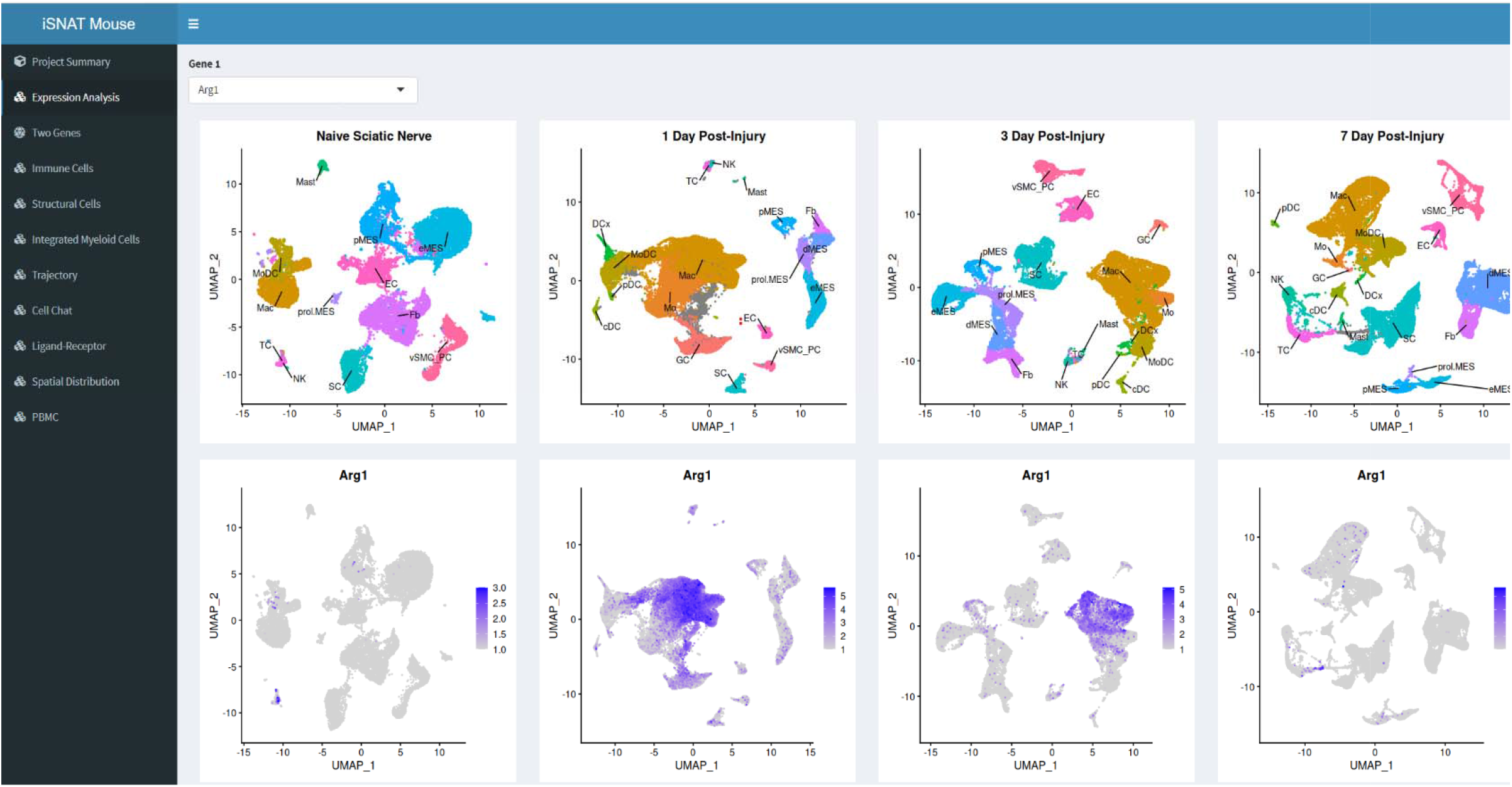
Screenshot of iSNAT webpage showing the menu with the different tabs/functions. As an example, the longitudinal expression changes of arginase-1 (Arg1) are shown. Time points include naïve sciatic nerve, 1-day post injury, 3-days post injury, and 7-days post injury. Top row shows UMAP plots with major cell types labeled. The bottom row shows expression of Arg1 during the first week post nerve injury. Note, Arg1 shows highest expression in macrophage (Mac) subpopulations in the 1-day and 3-day injured nerves.

### Myeloid cells in the injured nerve undergo rapid metabolic reprogramming

To demonstrate the application of iSNAT toward understanding how cellular function may change during the repair process, we focused on energy metabolism. Tissue repair is an energetically demanding process, suggesting cells must efficiently compete for limited resources, at the same time, the repair process requires highly coordinated action among diverse cell types. Nerve injury results in strong upregulation of the transcription factor *Hif1a*/hypoxia-induced factor 1α (Hif1α) in immune cells, including Mo, maturing Mac, and GC (**Fig. 3A-3D, 3I**). After a sharp increase at 1dpc, *Hif1a* levels remain elevated in Mo/Mac at 3dpc, before declining to pre-injury levels at 7dpc (**Fig. 3I**). Hif1α is a master regulator of cellular metabolism and the molecular machinery for glycolytic energy production (Nagao et al., 2019; Pearce and Pearce, 2013; Schuster et al., 2021). A metabolic shift, away from oxidative phosphorylation (OXPHOS) and toward aerobic glycolysis for the conversion of glucose into lactate, is known as Warburg effect (**Fig. 3J**). In immune cells the Warburg effect is of interest because it not only regulates metabolic pathways for energy production, but also gene expression to drive Mac toward a proinflammatory, glycolytic state (McGettrick and O’Neill, 2020; Palsson-McDermott et al., 2015).

**Figure 3.**
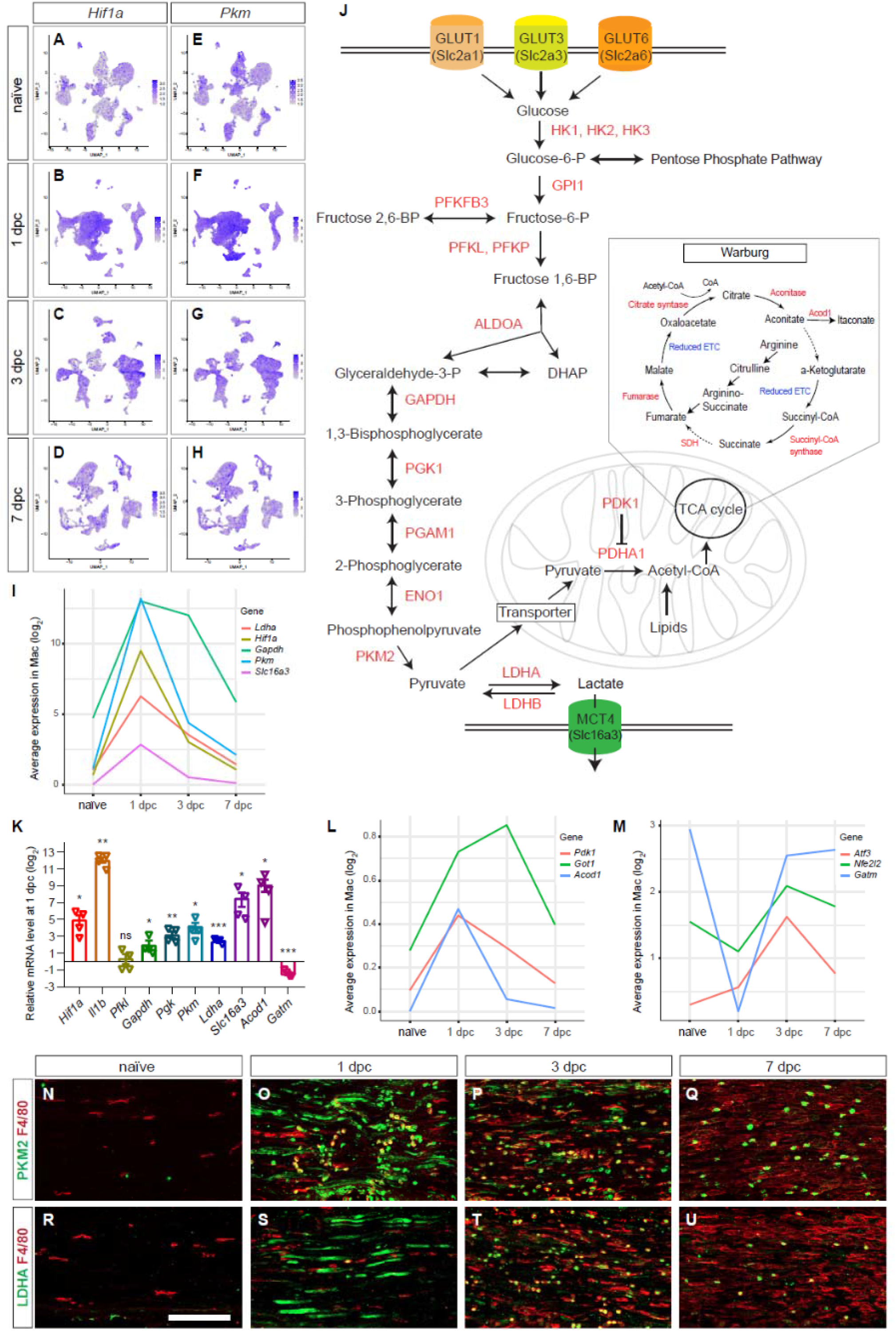
Application of iSNAT reveals metabolic reprogramming of immune cells. **(A-D)** Feature plots of *Hif1a* expression in naïve and injured sciatic nerve during the first week. **(E-H)** Feature plots of *Pkm* expression in naïve and injured sciatic nerve during the first week. Expression levels are projected onto the UMAP with a minimum expression cutoff of 1. **(I)** Injury regulated gene products associated with glycolysis, as inferred by scRNAseq data. Log2 average expression of genes for cells classified as Mac. **(J)** Metabolic pathways: glycolysis, the catabolism of glucose into pyruvate, and synthesis of nucleotides through the pentose phosphate pathway occur in the cytosol. The tricarboxylic acid cycle (TCA) takes place in mitochondria. The Warburg Effect allows for rapid ATP production through aerobic glycolysis and lactate production. Mitochondrial function is limited because of TCA fragmentation at steps marked with dotted arrows **(K)** Quantification of gene expression by qRT-PCR in the 1dpc nerve relative to naïve nerve. Log2-fold changes relative to naïve nerve are shown. Per gene product, n= 4 replicates. P-values, * <0.05, ** <0.001, *** < 0.0001, Student’s *t* test. ns, not significant. **(L, M)** Injury regulated gene products associated with inhibition of mitochondrial energy synthesis (**L**) and inflammation resolution (**M**), as inferred by scRNAseq data. Log2 average expression of genes for cells classified as Mac. (**N-U**) Longitudinal sections of naïve and injured sciatic nerves stained for macrophages (anti-F4/80, in red). Nerve sections were co-stained with (**N-Q**) anti-PKM2 (green) and (**R-U**) anti-LDHA (green). Proximal is to the left. Scale bar (**N-U**), 100 µm.

Analysis of gene products implicated in glucose metabolism, revealed that during the early injury response, GC and Mo/Mac express transporters for glucose (*Slc2a1/*GLUT1, *Slc2a3/*GLUT3) and hexose (*Slc2a6/*GLUT6) to import carbohydrates as a means of energy production. Moreover, there is rapid, injury-induced upregulation of most glycolytic enzymes, including hexokinases (*Hk1, Hk2, Hk3*), phospho-fructokinases (*Pfkp, Pfkl*), glyceraldehyde-3-phosphate dehydrogenase (*Gapdh*), and pyruvate kinase (*Pkm*) (**Fig. 3E-3H, 3I**). Injury regulated expression of *Hif1a* and key glycolytic enzymes was validated by qRT-PCR (**Fig. 3K**). The *Pkm* gene products, PKM1 and PKM2, convert phosphoenol pyruvate into pyruvate, the rate-limiting enzymes of glycolysis (**Fig. 3J**). PKM2 is of interest because of its nuclear role and interaction with Hif1α to promote expression of glycolytic enzymes and proinflammatory cytokines, including *Il1b* (Palsson-McDermott et al., 2015). Analysis of 1dpc nerve by qRT-PCR revealed a strong upregulation of *Il1b* when compared to naïve nerve (**Fig. 3K**). Mac in the injured nerve express elevated levels of *Ldha*, the enzyme that converts pyruvate to lactate (**Fig. 3I**). Evidence for intracellular lactate build-up, is the sharp increase in *Slc16a3*, encoding the monocarboxylate transporter 4 (MCT4) shuttling lactate out of cells (**Fig. 3I, 3K**). To validate the increase in glycolytic enzymes in injured nerve tissue, longitudinal sections of naïve, 1, 3, and 7dpc mice were stained with anti-PKM2 and anti-LDHA (**Fig. 3N-3U**). In naïve nerve, very few cells stain for PKM2 and LDHA. At 1dpc, PKM2 and LDHA are elevated in Mo/Mac at the nerve injury site. At 3dpc, PKM2 and LDHA are most abundant and preferentially detected in F4/80^+^ Mac. At 7dpc, staining is reduced and largely confined to a subset of F4/80^+^ Mac.

Lactate is far more than a metabolic waste product, since extracellular lactate has been shown to exert immunosuppressive functions, promote angiogenesis, axonal growth, and neuronal health (Chen et al., 2018; Funfschilling et al., 2012; Hayes et al., 2021; Kes et al., 2020). Lactate released by SC has axon protective effects and elevated lactate may be particularly important during the early injury response (Babetto et al., 2020). Evidence for inhibition of OXPHOS in the nerve during the early injury response is the upregulation of *Acod1* (**Fig. 3K, 3L**), an enzyme that converts aconitate into itaconate, thereby disrupting the TCA (**Fig. 3J**). Moreover, itaconate functions as an inhibitor of succinate dehydrogenase (SDH) leading to further inhibition of the TCA (Lampropoulou et al., 2016). Similarly, *Pdk1* (pyruvate dehydrogenase kinase 1) and *Got1* (glutamic-oxaloacetic transaminase), inhibitors of OXPHOS, are high at 1 and 3dpc, but low at 7dpc (**Fig. 3L**). Fragmentation of the TCA is a key feature of pro-inflammatory Mac and a hallmark of the Warburg effect (**Fig. 3J**) (Eming et al., 2021). During the inflammation resolution phase, Mac undergo metabolic reprogramming away from glycolysis toward OXPHOS, a switch that coincides with upregulation of the anti-inflammatory transcription factors *Atf3/*activating transcription factor 3 and *Nfe2l2*/Nrf2 (Mills et al., 2018). In the injured nerve, *Atf3* and *Nfe2l2* are low at 1dpc and upregulated at 3dpc (**Fig. 3M**). The mitochondrial enzyme *Gatm* (glycine amidinotransferase) is important for the biosynthesis of creatine, a molecule that facilitates ATP production from ADP. *Gatm* is transiently downregulated in Mo/Mac at 1dpc and increases in Mac at 3dpc and 7dpc (**Fig. 3M**).

Taken together, application of iSNAT provides multiple lines of evidence that during the early injury response Mo/Mac undergo rapid metabolic reprogramming to increase glycolytic flux and acquire a proinflammatory state. The proinflammatory state is short-lived as Mac rewire their metabolism toward OXPHOS and this is paralleled by a transition toward a pro-resolving phenotype.

### Myeloid cells in peripheral blood are not programed for glycolytic energy production

Because large numbers of blood-borne immune cells enter the injured PNS (Kalinski et al., 2020; Ydens et al., 2020), this prompted a deeper analysis of peripheral blood mononuclear cells (PBMC) before they enter the injured nerve. One important goal was to determine which PBMC enter the nerve upon injury and to assess how this impacts gene expression and metabolic profiles. The UMAP plot of 24,842 high-quality PBMC reveled 20 clusters (**Fig. 4A,4B**). Four clusters with B cells (BC1-BC4) expressing *Cd79a* (38% of all PBMC); plasmablasts (PB, 1.1%) expressing *Jchain/*joining chain of multimeric IgA and IgM*, Tnfrsf17/*TNF receptor superfamily member 17, and *Derl3/*derlin-3; platelets/megakaryocytes (P/M, 2%) expressing, *Pf4/*platelet factor 4, *Gp5/*glycoprotein V platelet, and *Plxna4/*PlexinA4; four clusters with T cells (TC1-TC4, 24.2%), expressing *Cd3e* with subsets of *Cd8b1* and *Cd4* cells, and NK cells (8.6%), expressing *Nkg7/*natural killer cell granule protein 7 (**Fig. 4C-4H**). Two clusters with granulocytes, including mature (GCm) (*Ly6g^hi^, Lcn2^hi^*, *Il1b*^-^) and immature (GCi) (*Cxcr2^hi^, Ly6g^low^, Il1b^hi^*) make up 10.6% of PBMC (**Fig. 4I,4J**). In addition, there are two Mo subpopulations, Mo1 (2.8%) expressing *Chil3^hi^, Ccr2^hi^, Cx3cr1^int^ Ly6c2^hi^, Csf1r^low^,* and *Ear2^-^*; Mo2 (5.3%) expressing *Cx3cr1^hi^, Csf1r^hi^, Ear2^hi^, Ccr2^-^, Ly6c2^-^,* and *Chil3^-^*, and a cluster with Mac (1.2%) expressing *Adgre1, Ccl24, Ltc4s,* and *Fn1* (**Fig. 4B, 4K-4O**). Mast cells (*Fcer1a, Cpa3*) make up 0.7% and DC (*Cd209a, Clec10a*) make up 1.8% of PBMC (**Fig. 4B, 4P,4Q**).

**Figure 4.**
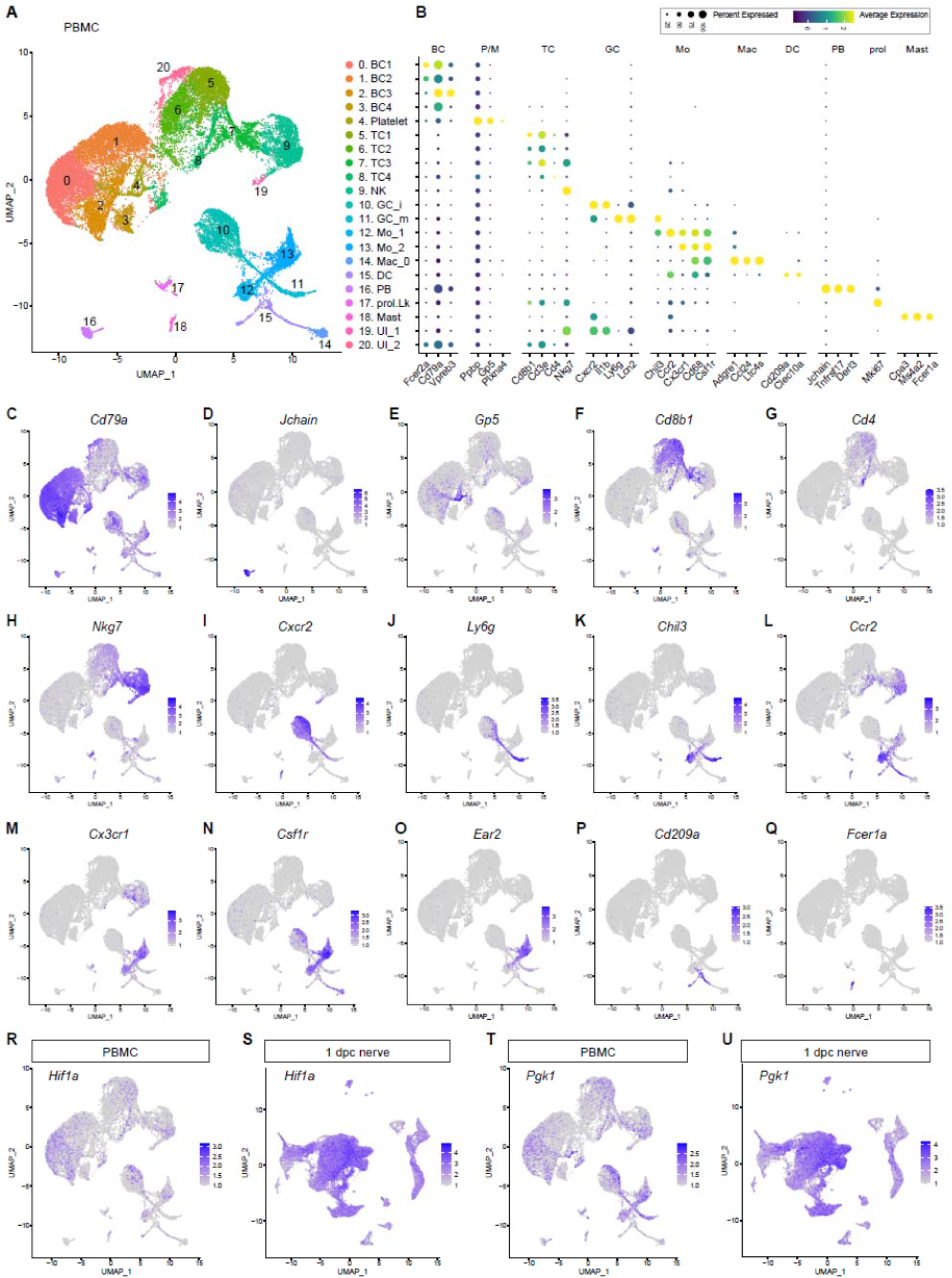
Cellular composition and peripheral blood mononuclear cells. (**A**) UMAP plot embedding of naïve mouse PBMC collected by cardiac puncture. Cell clusters identified harbor B cells (BC1-BC4), Platelets/megakaryocytes (P/M), T cells (TC1-TC4), Natural killer cells (NK), immature and mature granulocytes (mGC and iGC), two subpopulations of monocytes (Mo1 and Mo2), peripheral blood macrophages (MacPB), dendritic cells (DC), plasmablasts (PB), proliferating leukocytes (prol.Lk), mast cells (MAST), and two clusters with unidentified cells (UI1 and UI2). (**B**) A dotplot with marker genes for identification of PBMC. Color coded expression levels are shown. The dot size reflects the percentage of cells that express the gene. (**C-Q**) Feature plots of marker gene expression in PBMC. (**R-U**) Feature plots of *Hif1a* and *Pgk1* (phosphoglycerate kinase 1) expression in PBMC and the injured nerve at 1dpc. Expression levels are projected onto the UMAP with a minimum expression cutoff of 1 Abbreviations: *Cd79a*, CD79A antigen (immunoglobulin-associated alpha); *Gp5*, glycoprotein 5 (platelet); *Cd8b1*, CD8 antigen beta chain 1; *Cd4*, CD4 antigen; *Nkg7*, natural killer cell group 7 sequence; *Cxcr2*, chemokine (C-X-C motif) receptor 2; *Ly6g*, lymphocyte antigen 6 complex locus G; *Chil3*, chitinase-like 3 (Ym1); *Ccr2*, chemokine (C-C motif) receptor 2; *Cx3cr1*, chemokine (C-X3-C motif) receptor 1; *Csf1r*, colony stimulating factor 1 receptor; *Ear2*, eosinophil-associated, ribonuclease A family member 2; *Cd209a*, DC-SIGN (C-type lectin); *Jchain*, joining chain of multimeric IgA and IgM; *Fcer1a*, Fc receptor IgE high-affinity 1 alpha polypeptide.

Next, we used computational methods to extract blood myeloid cells (GCi, GCm, Mo1, Mo2, Mac, and DC) from the PBMC dataset for comparison with myeloid cells in the injured sciatic nerve at 1dpc, 3dpc, and 7dpc. Interestingly, Mo1 (*Ccr2^+^, Chil3^+^*) are transcriptionally more similar to 1dpc Mo in the nerve than Mo2 (*Ccr2^-^, Chil3^-^)*, suggesting Mo1 are the primary source of Mo entering the injured nerve. The small population of Mac in blood, is most similar to Mac-III at 1dpc (**Fig. 2B**), suggesting they may enter the injured nerve. GC in the 1dpc nerve are more similar to GCi than GCm in blood, suggesting preferential entry of GCi. Because Mo/Mac in the 1dpc nerve are metabolic programmed for glycolytic energy production (**Fig. 3**), we ask whether their precursors in blood exhibit a similar metabolic profile. Strikingly, when compared to Mo/Mac in the injured nerve, blood Mo1, Mo2, and Mac show either low levels or lack expression of *Hif1a* (**Fig. 4R, 4S**), the glycolytic enzymes *Pgk1*/phosphoglycerate kinase-1 (**Fig. 4T.4U**), *Pfkl*/phosphofructokinase liver type (**Fig. 4, Suppl. 1A-1E**), and the lactate exporter *Slc16a3* (**Fig. 4, Suppl. 1F-1J**). This suggests that upon nerve entry, circulating Mo/Mac become activated and undergo metabolic reprogramming toward glycolytic energy production. *Arg1,* an important regulator of innate and adaptive immune responses, is not detected in peripheral blood Mo/Mac, however upon nerve entry, Mo/Mac rapidly upregulate *Arg1*. Regulation of *Arg1* expression is very dynamic, levels are highest at 1dpc and 3dpc, then rapidly decline, reaching baseline at 7dpc (**Fig. 4, Suppl. 1K-1O**). Thus, our studies show that upon nerve entry, circulating Mo/Mac undergo extensive transcriptional remodeling of glucose metabolism.

**Figure 4, Suppl. 1.**
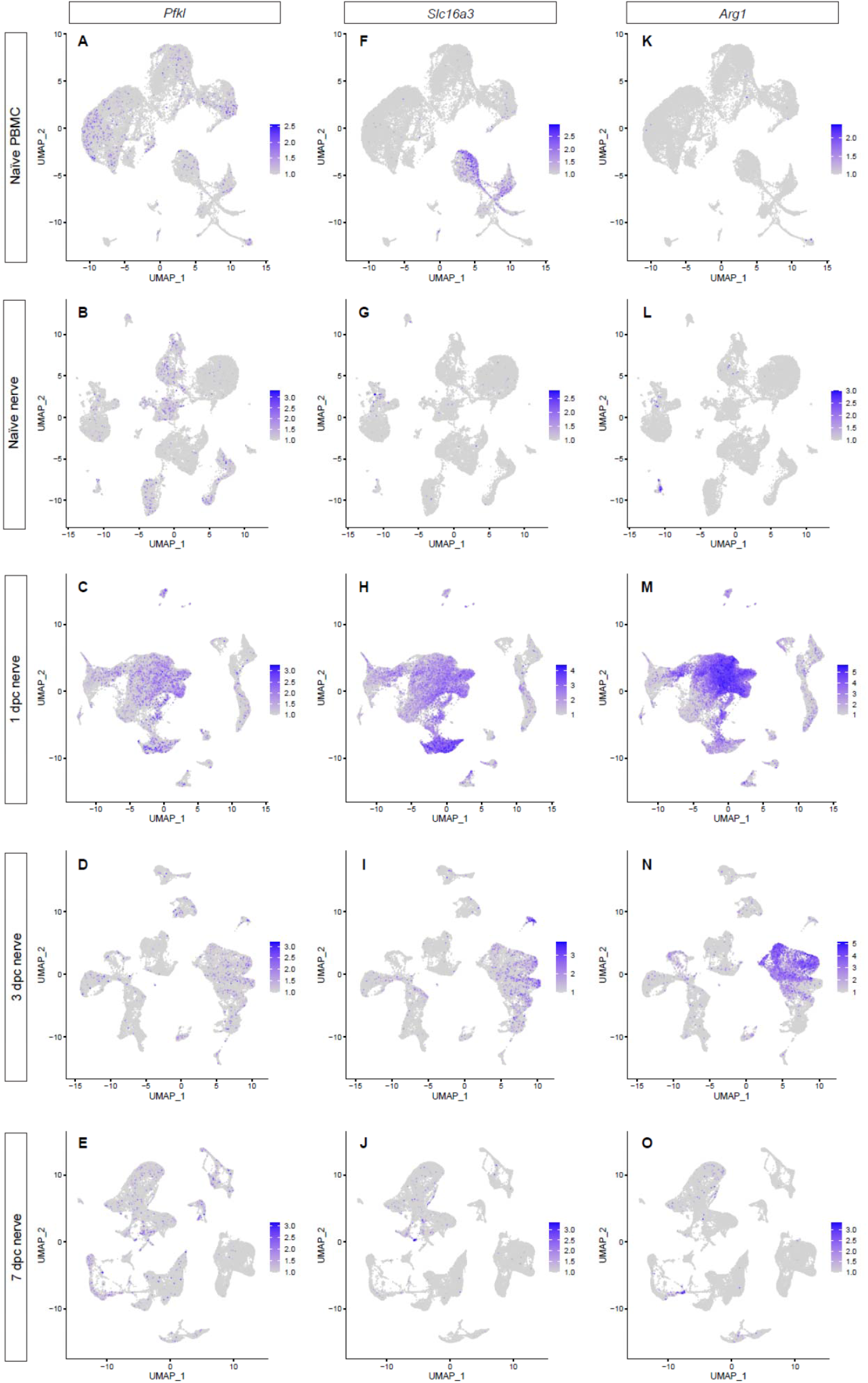
Activation of circulating immune cells upon nerve entry. Feature plots for the glycolytic enzyme *Pfkl* (**A-E**), the monocarboxylate transporter *Slc16a3* (**F-J**), and the arginine degrading enzyme *Arg1* (**K-O**), in peripheral blood monocytes (PBMC), naïve nerve, 1dpc, 3dpc, and 7dpc nerve. Expression levels are projected onto the UMAP with a minimum expression cutoff of 1.

### CellChat identifies large cell-cell communication networks activated during nerve repair

To understand cell-cell communication networks during the repair process, we interrogated cell surface protein interactions using CellChat (Jin et al., 2021). To facilitate identification and mining of predicted protein-protein interactions, we added CellChat as a feature to iSNAT. The output of CellChat is the probability for cell-cell communication to occur via specific ligand-receptor systems. In naïve and injured nerve, CellChat identified hundreds of ligand-receptor pairs among different cell groups, which were further categorized into 64 major signaling networks. Prominent examples of injury regulated networks include CXCL- (**Fig. 5A-5H**), CCL-(**Fig. 5I-5P**), CX3C-type chemokine, serum amyloid A (SAA), progranulin (GRN), and osteopontin/secreted phosphoprotein 1 (SPP1). Many of the ligand-receptor pairs in these networks function in leukocyte chemotaxis (SenGupta et al., 2019), and thus, can be mined to determine cellular sources and receptor mechanisms that promote GC and Mo infiltration into the injured nerve. For example, at 1dpc, Mac-II and Mac-III express *Cxcl2, Cxcl3, Cxcl5*, and *Ppbp*/pro-platelet basic protein/CXCL7, encoding activators and chemoattractants for GC that signal through the receptor *Cxcr2*. Mac-I show highest levels of *Ccl2*, Mac-II express *Pf4/*CXCL4*, Grn/*progranulin, and Mac-V express *Ccl24, Spp1*, and *Il1rn*. Analysis of chemotactic receptors further revealed that GC express *Cxcr2*, the formyl peptide receptors *Fpr1* and *Fpr2*, *Tnfrsf1a/*TNF receptor 1*, Ltb4r1*/leukotriene B4 receptor 1, and *Ccr1*. Mo strongly express the chemotactic receptors *Ccr2*, *Ccr5,* and *Ltb4r1,* indicating that GC and Mo use overlapping, yet distinct mechanisms for entering the injured nerve. Many of the chemotactic molecules including *Ccl2, Ccl7*, *Ccl9*, and *Ccl12* show highest expression at 1dpc. At 3dpc, chemokines are reduced compared to peak levels, and by 7dpc have declined further, approaching steady state levels comparable to naïve nerves (**Fig. 5A-5P**).

**Figure 5.**
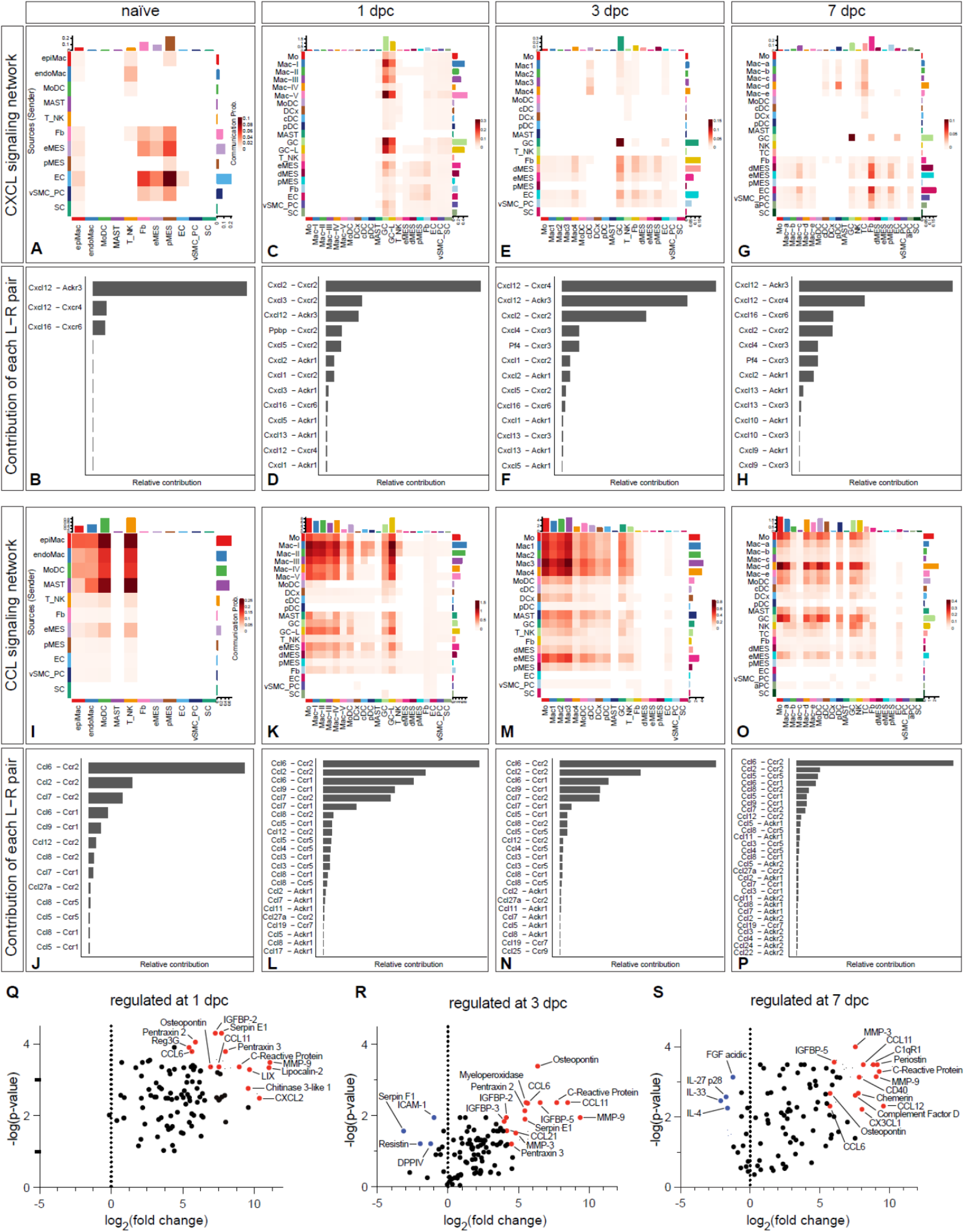
CellChat reveals chemotactic cell-cell communication networks in the injured PNS. Hierarchical plots of CellChat analysis showing the inferred intercellular communication network for (**A-H**) CXCL-chemokines and (**I-P**) CCL-cytokines in naïve nerve and during the first week following injury. The sender cells (ligand sources) are shown on the y-axis and receiving cells on the x-axis. The communication probabilities for cells that communicate with each other are indicated. (**B,D,F,H**) The bar graphs show the contributions of CXCL ligand receptor pairs for each time point. (**J,L,N,P**) The bar graphs show the contributions of CCL ligand receptor pairs for each time point. (**Q-S**) Volcano plot of extracellular proteins detected by ELISA compared to naïve nerve. The most abundant and strongly upregulated proteins in the 1dpc nerve (**Q**), the 3dpc nerve (**R**), and the 7dpc nerve (**S**) are shown. The normalized signal on the x-axis shows the log2 fold-change and the y-axis shows the -log(p-value), normalized to naïve nerve.

In addition to immune cells, CellChat identifies structural cells as a major hub for chemotactic factors, providing evidence for prominent immune-stroma crosstalk. In particular, eMES rapidly increase the production of *Ccl2, Ccl7, Il6, Il11, Cxcl1, Cxcl5, Cxcl12, Cx3cl1, Spp1,* and *Lif*. In addition, injured eMES show elevated expression of serum amyloid A apolipoproteins (*Saa1, Saa2, Saa3*), chemotactic molecules for GC and Mo/Mac. *Ptx3*/pentraxin 3, a factor implicated in wound healing (Erreni et al., 2017), is regulated by injury and rapidly increases in eMES and dMES at 1dpc and 3dpc. Distinct immune molecules are expressed by pMES, including *Serping1* (complement C1 inhibiting factor), *Cfh* (complement factor H), *Lbp* (lipopolysaccharide binding protein). Upon injury, dMES upregulate *Ccl2, Ccl7, Cxcl1, Cxcl5, Cxcl12,* underscoring the importance of diverse stromal cells in orchestrating the immune response (**Fig. 5, Suppl 1**). Because SNC inflicts vascular damage, we searched CellChat for injury induced angiogenic signaling pathways, top hits include VEGF, ANGPT (angiopoietin), ANGPTL (angiopoietin-like proteins), SEMA3 (semaphorin 3), EPHA, and EPHB (ephrin receptors). While the importance of angiogenesis and angiogenic factors in PNS repair has been appreciated, CellChat informs on specific ligand-receptor systems and shows which cell types produce these factors.

**Figure 5, Suppl. 1.**
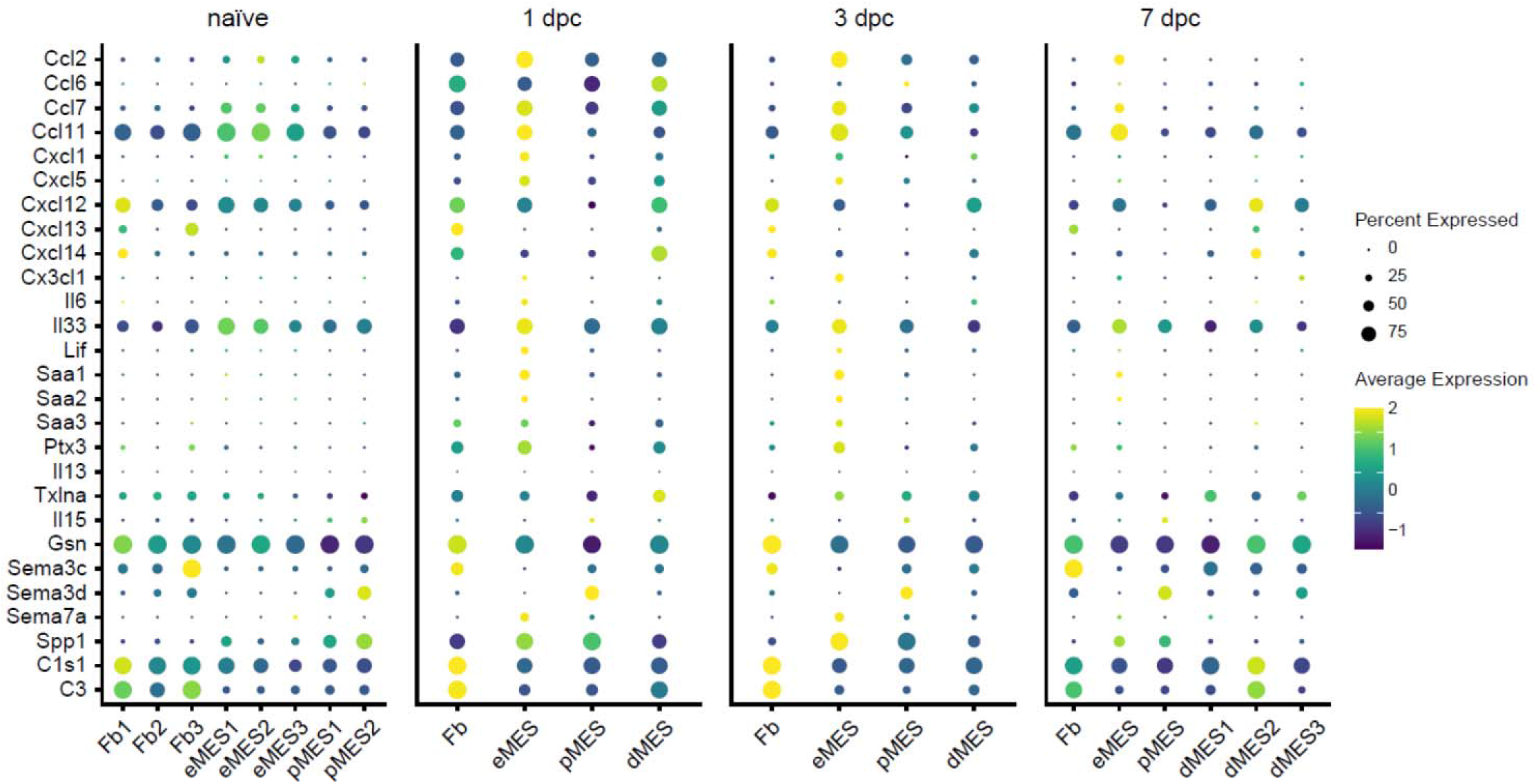
Structural cells in the injured nerve directly shape the immune microenvironment. Dotplots of structural cells, including Fb (fibroblasts), eMES (endoneurial mesenchymal cells), pMES (perineurial mesenchymal cells), and dMES (differentiating mesenchymal cells). Expression of extracellular immune molecules, as detected by scRNAseq, during the first week following nerve injury are shown. Color coded gene expression levels, normalized to average gene expression is shown. The dot size reflects the percentage of cells that express the gene.

### Identification of sciatic nerve injury regulated extracellular proteins

While CellChat predicts a large number of ligand-receptor interactions to occur in the injured nerve, we used ELISA to probe nerve lysates to ask whether some of the corresponding extracellular proteins are present at 1, 3, and 7dpc (**Fig. 5, Suppl. 2**). ELISA data obtained from naïve nerve and serum were used for comparison (**Fig. 1, Suppl. 3**). Proteins strongly upregulated by SNC include chemokines (CCL6, CCL11, CCL12, CXCL2, LIX, and osteopontin), serpin family members (Serpin E1/PAI-1, Serpin F1/PEDF), pentraxins (NPTX2, PTX3, CRP), ECM degrading matrix metallopeptidases (MMP9, MMP3, MMP2), and IGF binding proteins (IGFBP2, IGFBP3, IGFBP5, IGFBP6) (**Fig. 5 Suppl. 2**). In the 1dpc nerve, prominently detected proteins include MMP9, CCL6, NPTX2, PTX3, CST3, MPO, Osteopontin, SerpinF1, CXCL5, FGF1, AHSG, CCL11, MMP3, IGFBP2, (**Fig. 5Q, Fig 5, Suppl. 2D**). Top proteins detected in the nerve at 3dpc include CCL11, CCL6, FGF1, NPTX2, PTX3, IGFBP6, Osteopontin, MPO, IGFBP5, MMP9, CST3, MMP3, Endostatin (**Fig. 5R, Fig 5, Suppl. 2E**). Top proteins detected in the nerve at 7dpc include CCL6, CCL11, IL1α, FGF21, Endostatin, F3, AHSG, ENG, CST3, MMP2, Adipoq, INFγ, IL33, and LDLR (**Fig. 5S, Fig 5, Suppl. 2F).**

**Figure 5, Suppl. 2.**
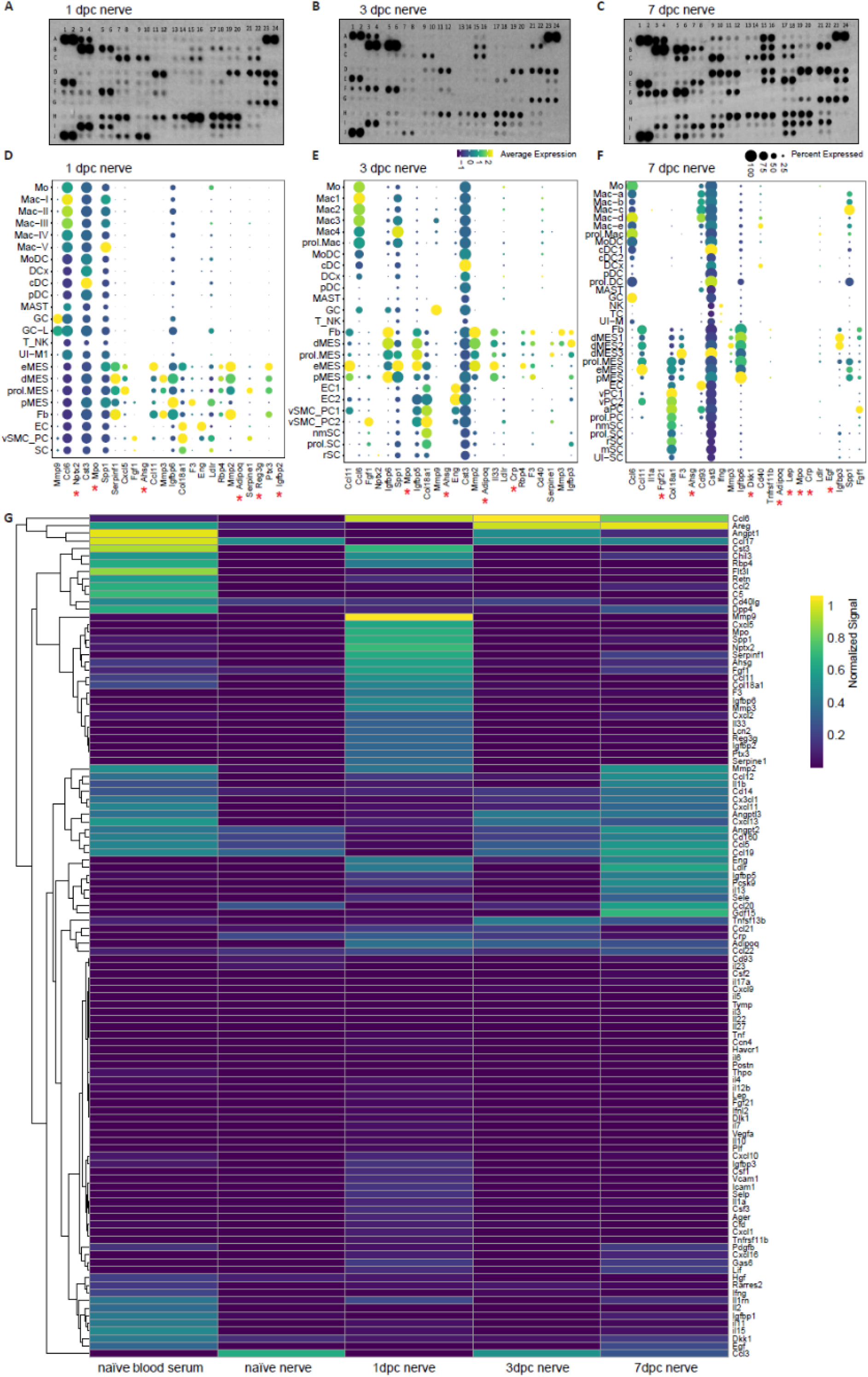
Identification of extracellular proteins in the injured sciatic nerve by ELISA. (**A-C**) ELISA membranes probed with injured nerve lysates prepared at (A) 1dpc, (B) 3dpc, and (C) 7dpc. (**D-F**) Dotplots of scRNAseq data from 1dpc, 3dpc, and 7dpc, for the top 25 gene products detected by ELISA. Relative RNA expression levels, normalized to average gene expression (color coded) are shown. For each cell cluster, the percentile of cells expressing a specific gene product is indicated by the dot size. Gene products marked with a red star are detected by ELISA but not by scRNAseq. (**G**) Heatmap shows proteins detected by ELISA in serum of naïve mice, naïve sciatic nerve trunk, and injured sciatic nerve trunk at 1dpc, 3dpc, and 7dpc. Number of biological replicates for ELISA, n = 1. Relative protein levels were normalized to internal controls on ELISA membranes shown at coordinates (A1,A2), (A23,A24), (J1,J2). Coordinates of proteins that can be detected by the ELISA: (A3, A4) Adiponectin [*Adipoq*], (A5, A6) Amphiregulin [*Areg*], (A7, A8) Angiopoientin-1 [*Angpt1*], (A9, A10) Angiopoientin-2 [*Angpt2*], (A11, A12) Angiopoientin-like 3 [*Angptl3*], (A13, A14) BAFF [*Tnfrsf13b*], (A15, A16) C1qR1 [*Cd93*], (A17, A18) CCL2 [*Ccl2*], (A19, A20) CCL3 [*Ccl3*], (A21, A22) CCL5 [*Ccl5*], (A23, A24) Reference spots, (B3, B4) CCL6 [*Ccl6*], (B5, B6) CCL11 [*Ccl11*], (B7, B8) CCL12 [*Ccl12*], (B9, B10) CCL17 [*Ccl17*], (B11, B12) CCL19 [*Ccl19*], (B13, B14) CCL20 [*Ccl20*], (B15, B16) CCL21 [*Ccl21*], (B17, B18) CCL22 [*Ccl22*], (B19, B20) CD14 [*Cd14*], (B21, B22) CD40 [*Cd40*], (C3, C4) CD160 [*Cd160*], (C5, C6) Chemerin [*Rarres2*], (C7, C8) Chitinase 3-like 1 [*Chil3*], (C9, C10) Coagulation Factor III [*F3*], (C11, C12) Complement Component C5 [*C5*], (C13, C14) Complement Factor D [*Cfd*], (C15, C16) C-Reactive Protein [*Crp*], (C17, C18) CX3CL1 [*Cx3cl1*], (C19, C20) CXCL1 [*Cxcl1*], (C21, C22) CXCL2 [*Cxcl2*], (D1, D2) CXCL9 [*Cxcl9*], (D3, D4) CXCL10 [*Cxcl10*], (D5, D6) CXCL11 [*Cxcl11*], (D7, D8) CXCL13 [*Cxcl13*], (D9, D10) CXCL16 [*Cxcl16*], (D11, D12) Cystatin C [*Cst3*], (D13, D14) DKK-1 [*Dkk1*], (D15, D16) DPPIV [*Dpp4*], (D17, D18) EGF [*Egf*], (D19, D20) Endoglin [*Eng*], (D21, D22) Endostatin [*Col18a1*], (D23, D24) Fetuin A [*Ahsg*], (E1, E2) FGF acidic [*Fgf1*], (E3, E4) FGF-21 [*Fgf21*], (E5, E6) Flt-3 Ligand [*Flt3l*], (E7, E8) Gas 6 [*Gas6*], (E9, E10) G-CSF [*Csf3*], (E11, E12) GDF-15 [*Gdf15*], (E13, E14) GM-CSF [*Csf2*], (E15, E16) HGF [*Hgf*], (E17, E18) ICAM-1 [*Icam1*], (E19, E20) IFN-gamma [*Ifng*], (E21, E22) IGFBP-1 [*Igfbp1*], (E23, E24) IGFBP-2 [*Igfbp2*], (F1, F2) IGFBP-3 [*Igfbp3*], (F3, F4) IGFBP-5 [*Igfbp5*], (F5, F6) IGFBP-6 [*Igfbp6*], (F7, F8) IL-1alpha [*Il1a*], (F9, F10) IL-1Beta [*Il1b*], (F11, F12) IL-1ra [*Il1rn*], (F13, F14) IL-2 [*Il2*], (F15, F16) IL-3 [*Il3*], (F17, F18) IL-4 [*Il4*], (F19, F20) IL-5 [*Il5*], (F21, F22) IL-6 [*Il6*], (F23, F24) IL-7 [*Il7*], (G1, G2) IL-10 [*Il10*], (G3, G4) IL-11 [*Il11*], (G5, G6) IL-12 p40 [*Il12*], (G7, G8) IL-13 [*Il13*], (G9, G10) IL-15 [*Il15*], (G11, G12) IL-17A [*Il17a*], (G13, G14) IL-22 [*Il22*], (G15, G16) IL-23 [*Il23*], (G17, G18) IL-27 p28 [*Il27*], (G19, G20) IL-28 [*Ifnl3*], (G21, G22) IL-33 [*Il33*], (G23, G24) LDL R [*Ldlr*], (H1, H2) Leptin [*Lep*], (H3, H4) LIF [*Lif*], (H5, H6) Lipocalin-2 [*Lcn2*], (H7, H8) LIX [*Cxcl5*], (H9, H10) M-CSF [*Csf1*], (H11, H12) MMP-2 [*Mmp2*], (H13, H14) MMP-3 [*Mmp3*], (H15, H16) MMP-9 [*Mmp9*], (H17, H18) Myeloperoxidase [*Mpo*], (H19, H20) Osteopontin [*Spp1*], (H21, H22) Osteoprotegerin [*Tnfrsf11b*], (H23, H24) PD-ECGF [*Tymp*], (I1, I2) PDGF-BB [*Pdgfb*], (I3, I4) Pentraxin 2 [*Nptx2*], (I5, I6) Pentraxin 3 [*Ptx3*], (I7, I8) Periostin [*Postn*], (I9, I10) Pref-1 [*Dlk1*], (I11, I12) Proliferin [*Prl2c2*], (I13, I14) Proprotein Convertase 9 [*Pcsk9*], (I15, I16) RAGE [*Ager*], (I17, I18) RBP4 [*Rbp4*], (I19, I20) Reg3G [*Reg3g*], (I21, I22) Resistin [*Retn*], (J1, J2) Reference spots, (J3, J4) E-Selectin [*Sele*], (J5, J6) P-Selectin [*Selp*], (J7, J8) Serpin E1 [*Serpine1*], (J9, J10) Serpin F1 [*Serpinf1*], (J11, J12) Thrombopoietin [*Thpo*], (J13, J14) TIM-1 [*Havcr1*], (J15, J16) TNF-alpha [*Tnf*], (J17, J18) VCAM-1 [*Vcam1*], (J19, J20) VEGF [*Vegf*], (J21, J22) WISP-1 [*Ccn4*], (J23, J24) negative control.

To examine whether the corresponding transcripts are expressed in the injured nerve, and to identify the cellular sources, we generated dotplots from scRNAseq datasets (**Fig. 5, Suppl. 2D-2F**). For the top 25 proteins detected by ELISA, there is a close correlation at the transcriptional level with many gene products strongly expressed by structural cells. Some of the top gene products detected by ELISA but not by scRNAseq, are serum proteins (**Fig. 1, Suppl. 3D)**. Specifically, they include fetuin A (AHSG), CRP (C reactive protein), IGFBP-2, Reg3g (regenerating family member 3γ), and CFD (complement factor D). Because mice were transcardially perfused before nerves were harvested, this shows that numerous systemic factors accumulate in the injured nerve. CRP is of interest, because it binds to the surface of dead cells and activates complement C1q (Alnaas et al., 2017). Collaboration between CRP and C1q is supported by the strong expression of C1q components (*C1qa, C1qb, and C1qc*) in most Mac subpopulations. CFD is a serine protease required for the formation of C3 convertase. Soluble C1qR1/CD93 functions as a bridging molecule that aids apoptotic cell binding to professional phagocytes for removal by efferocytosis (Blackburn et al., 2019). Taken together, we provide validation for many injury-regulated gene products identified by scRNA-seq and show that serum proteins that function in opsonization feature prominently in the PNS following nerve crush injury. Interestingly, several abundantly detected serum proteins (Angpt1, Angpt2, Angptl3, IGFBP-1) seem not to enter the injured nerve parenchyma (**Fig. 5, Suppl. 1**). This suggests that these proteins are either rapidly degraded or that SNC causes a partial breakdown of the BNB, allowing only some serum proteins to enter the injured nerve.

### The injured sciatic nerve harbors distinct dendritic cell populations

Only few DC are present in the naïve nerve, and they can be distinguished from resident Mac by their prominent expression of *Flt3* (FMS-like tyrosine kinase 3) and *Napsa* (**Fig. 1L**). During the first week following nerve injury, MoDC, identified by *Cd209a* expression, gradually increase (**Fig. 2H, 2P, 2X**). Gene expression analysis in iSNAT identifies additional DC subpopulations, including conventional dendritic cells (cDC, *Sept3/*septin 3 GTPase, *Clec9a*/C-type lectin domain containing 9A), plasmacytoid dendritic cells (pDC*, Ccr9* and *Siglech*), and DCx (*Ccr7/*chemokine receptor 7, *Ccl17, Ccl22*, *Fscn1*/fascin an actin-binding protein) (**Fig. 5, Suppl. 3A**). DCx are reminiscent of Langerhans cells and express *Ccr7*, suggesting they are mature cells destined for homing, the migration from injured nerve to draining lymph nodes (Bros et al., 2011; Liu et al., 2021). At 7dpc, two subpopulations of conventional DC (cDC1 and cDC2) are detected. Previous work has established that cDC1 are specialized in cross presentation and activation of cytotoxic TC, and cDC2 are specialized in driving helper TC responses (Steinman, 2012). The two populations can be distinguished based on preferential expression of marker genes, cDC1 (*Itgam^-^*/CD11b, *Itgae^+^*/integrin αE, *Clec9a^+^, Tlr3^+^/*toll-like receptor 3*, Xcr1^+^/*X-C motif chemokine receptor 1*^+^*) and cDC2 (*Itgam^+^, Clec9a^-^*, *Xcr1^-^, Irf4^+^/*interferon regulatory factor 4, *Slamf7*/CD2-like receptor activating cytotoxic cells).

**Figure 5, Suppl. 3.**
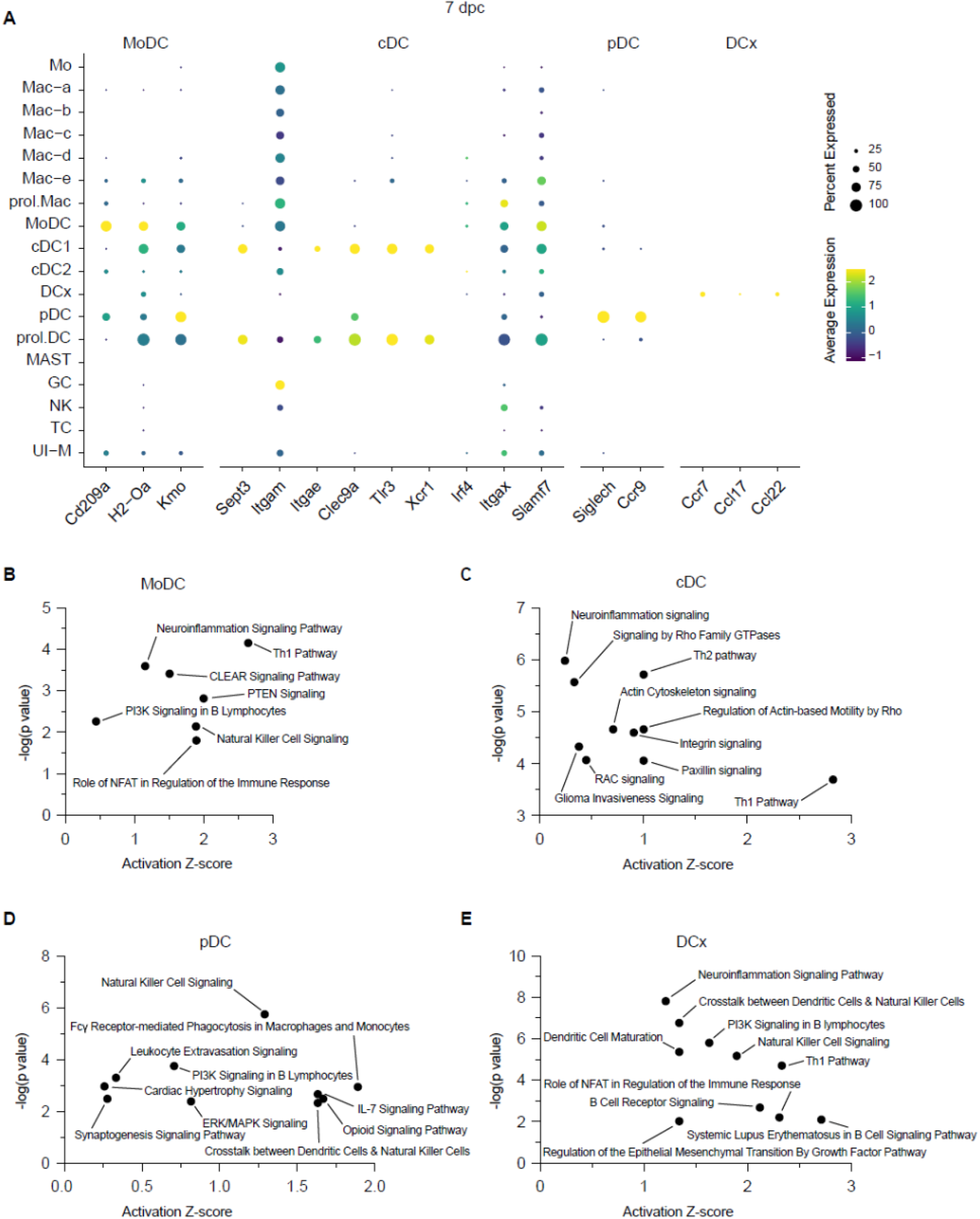
Identification of distinct dendritic cell populations in the injured PNS. (**A**) Dotplot for DC marker genes at 7dpc identifies MoDC (monocyte-derived dendritic cells), cDC (conventional dendritic cells), pDC (plasmacytoid dendritic cells), and DCx (mature dendritic cells destined for homing to draining lymph nodes). Relative RNA expression levels, normalized to average gene expression (color coded) are shown. For each cell cluster, the percentile of cells expressing a specific gene product is indicated by the dot size. Ingenuity pathway analysis at 7dpc for (**B**) MoDC, (**C**) cDC, (**D**) pDC, and (**E**) DCx. Top-scoring enriched canonical pathways in each cell type are represented by activation z-scores and p-values.

CellChat identifies an FLT3 signaling network in the injured nerve and predicts communication between TC, NK, and all four DC populations (data not shown). Nerve injury triggers strong expression of interferon-inducible genes in MoDC (*Ifi30, Ifitm1, Ifitm6*), cDC1 (*Ifi30, Ifi205, Irf8*), and cDC2 (*Ifi30*). MoDC express high levels of macrophage galactose-type lectin, encoded by the C-type lectin receptor *Clec10a*, typically found on tolerogenic antigen-presenting cells (van Kooyk et al., 2015). The *Clec9a* gene product is expressed by cDC and binds to filamentous actin exposed by damaged or ruptured cells (Zhang et al., 2012). The strong expression of *Clec9a* by cDC1 at 7dpc suggests a role in antigen uptake and presentation. While DC are superior to Mac in presenting antigen and express high levels of *MHC class II* components (*H2-Aa, H2-Ab1, H2-DMa, H2-Eb1, Cd74*), most Mac in the naïve nerve, and some Mac in the injured nerve, express *MHC class II* and *Cd74*, suggesting they are endowed with antigen presenting capabilities. Mac in the injured nerve can be distinguished from DC by their preferential expression of complement C1q (*C1qa, C1qb, C1qc*), *C3ar1/*complement C3a receptor 1, *Fcgr1/*CD64*, Trem2, Slc11a1*, *Adgre1*, and *Apoe*/apolipoprotein E.

To infer potential functions for DC in the injured nerve, we carried out ingenuity pathway analysis (IPA) (**Fig. 5, Suppl. 3B-3E**). At 7dpc, the top positively regulated pathways include, *Th1 pathway* for MoDC and *Th2 pathway* for cDC1, suggestive of proinflammatory and inflammation resolving functions, respectively. The top pathways for pDC are *NK cell signaling* and *Fcγ receptor-mediated phagocytosis* and for DCx *neuroinflammation signaling* and *crosstalk between DC and NK cells*. Taken together, we identify different DC populations in the injured PNS and pathway analysis predicts extensive crosstalk between DC-NK and DC-TC.

### PNS injury triggers a gradual T cell response

For TC classification, we used the “*Expression Analysis*” function in iSNAT. Most TC express the T cell receptor (TCR) α-chain constant (*Trac*) and β-chains (*Trbc1* and *Trbc2*), suggesting they are αβ TC (**Fig. 5, Suppl. 4A-4D**). Only few TC are present in the naïve sciatic; however, TC gradually increase upon injury, and by 7dpc make up ∼10% of immune cells. Few *Mki67*/Ki67^+^ TC are observed in the injured nerve and the majority expresses *Ms4a4b*, a negative regulator of cell proliferation (**Fig. 2, Suppl. 2F-2H**). This suggests that TC expansion is primarily due to nerve infiltration, rather than local proliferation. In support of this, TC strongly express *Cxcr6*, a chemotactic receptor for *Cxcl16* expressed by Mac and DCx in the injured nerve. Nearly all TC express CD3 (*Cd3g, Cd3d, Cd3e,* and *Cd3z/Cd247*), a key component of the TCR-CD3 complex (**Fig. 5, Suppl. 4E-4H**). At 7dpc, the TC population is comprised of CD4^+^ T helper cells (Th), expressing *Cd4*, (**Fig. 5, Suppl. 4I-4L**) and CD8 cells expressing *Cd8a* and *Cd8b1,* suggesting they are CD8αβ^+^ cells (data not shown). Differentiation of CD4^+^ Th into pro-inflammatory Th1 or anti-inflammatory Th2 effector cells is controlled by the transcription regulators T-bet (*Tbx21*) and GATA3 (*Gata3*), respectively (Jenner et al., 2009). *Gata3*^+^ Th2 cells are observed in naïve and injured nerves, while *Tbx21^+^* Th1 are largely absent from naïve nerve but increase following injury and express the killer cell lectin like receptors *Klrd1* and *Klrc1*, markers of an activated pro-inflammatory state. Pathway analysis of TC at 7dpc, identified *Th2 pathway* and *Th1 pathway* as top hits (**Fig. 5, Suppl. 4Q**). It is well established that the Th1 response plays a key role in neuropathic pain development and persistence (Davies et al., 2020; Moalem and Tracey, 2006). At 7dpc, there is a small population of γδ TC (*Trdc*), and some T regulatory cells (Tregs), a specialized subpopulation of TC that acts to suppress the immune response. In the PNS, Tregs (*Cd4*, *Il2ra*/CD25, *Foxp3)* are of interest, because they may function in self-tolerance and pain mitigation (Davoli-Ferreira et al., 2020).

**Figure 5, Suppl. 4.**
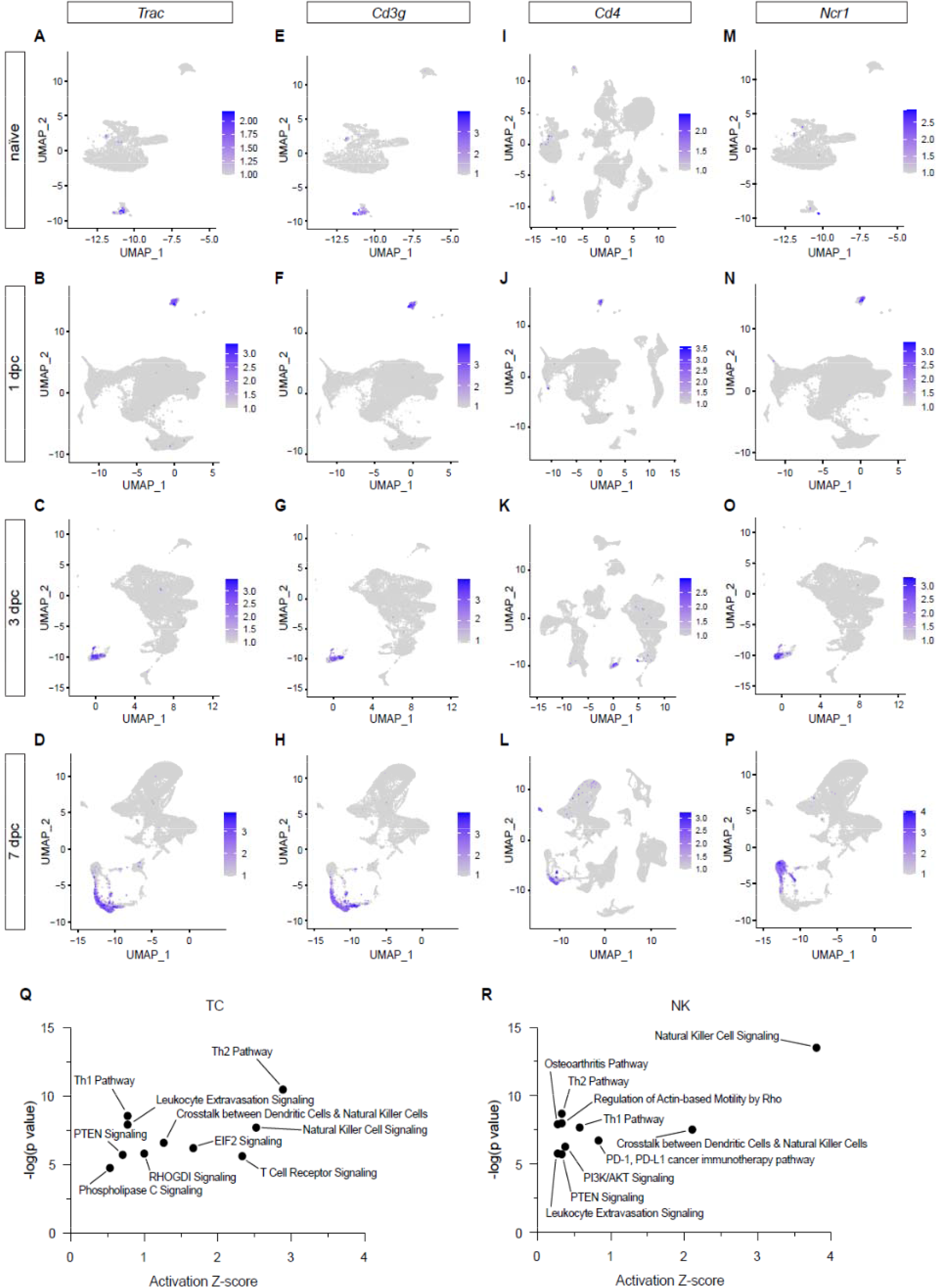
Description of TC and NK populations in the injured PNS. Feature plots for (**A-D**) *Trac/*T cell receptor alpha constant, (**E-H**) *Cd3g/*CD3 gamma subunit of T cell receptor complex, (**I-L**) *Cd4/*T cell surface glycoprotein CD4, and (**M-P)** *Ncr1*/Natural cytotoxicity triggering receptor-1 of naïve, 1, 3, and 7dpc nerve identify TC subpopulations and NK cells. Expression levels are projected onto the UMAP with a minimum expression cutoff of 1. (**Q, R**) Ingenuity pathway analysis for TC and NK at 7dpc. Top-scoring enriched canonical pathways are represented by activation z-scores and p-values.

### Natural Killer cells increase following nerve injury

In contrast to TC, NK do not express *Cd3g, Cd3d, Cd3e*, but strongly express natural cytotoxicity receptor (*Ncr1*), killer cell lectin-like receptor subfamily B member 1C (*Klrb1c/*NK1.1), the pore-forming glycoprotein perforin (*Prf1*), and the granzyme family serine proteases (*Gzma, Gzmb*) (**Fig. 5, Suppl. 4M-4P**). Granzymes are delivered into target cells through the immunological synapse to cause cell death. While the full spectrum of cells targeted by NK has yet to be determined, NK cytotoxic factors have been shown to accelerate degeneration of damaged axons (Davies et al., 2019). Compared to NK, CD8^+^ TC express low levels of granulysin (*Gnly*), perforin (*Prf1*), and granzymes (*Gzma, Gzma*) suggesting limited cytotoxic activity. However, similar to NK, many TC express *Nkg7*, a natural killer cell granule protein, and the killer cell lectin like receptor K1 (*Klrk1*/NKG2D), indicative of some cytotoxic abilities. NK (and some TC) strongly express *Klrc1*/NKG2A (CD94), an immune inhibitory checkpoint gene. The ligand for NKG2A is the non-classical MHC1 molecule Qa-2 (encoded by *H2-q7*), expressed by NK and TC. CellChat predicts a high probability for an MHC-I signaling network between CD8^+^ TC, NK, and pDC, and an MHC-II signaling network between CD4^+^ TC and with DC and Mac in the injured nerve. IPA for NK at 7dpc, identifies *NK signaling*, *Th2 pathway*, and *crosstalk between DC and NK* as top hits (**Fig. 5, Suppl. 4R**). The majority of NK and TC in the injured nerve produce interferon γ (*Ifng*), some TC produce TNFα (*Tnf*), and few produce GM-CSF (*Csf2*). In addition, NK and TC produce chemoattractants and survival signals for DC, including *Flt3l*, *Ccl5, Xcl1/*lymphotactin, suggesting they directly regulate DC migration and function.

### Tracking myeloid cell transcriptional states and maturation in the injured nerve

Mo/Mac are often described as highly “plastic” cells, educated by the local tissue environment. While Mac subpopulations in the injured PNS have been cataloged (Kalinski et al., 2020; Ydens et al., 2020), tracking them over time, as they mature, has not yet been attempted. To better understand Mo/Mac maturation in the injured nerve, we took advantage of our longitudinal scRNAseq datasets. Computational methods were used to extract myeloid cells from PBMC, naïve nerve, and the three post-injury time points to carry out an integrated data analysis (**Fig. 6A-E**). When comparing the UMAP plots of naïve sciatic nerve and PBMC, there is little overlap, indicative of largely distinct immune compartments (**Fig. 6A,6B**). When comparing myeloid cells in blood with 1dpc nerve, the rapid entry of GC into the nerve is readily detected and GC are largely absent from the 7dpc nerve (**Fig. 6, Suppl. 1F-1J**). Interestingly, 1dpc GC are more similar to *Il1b^+^* GCi than *Il1b^-^* GCm in blood (**Fig. 6, Suppl. 1K-1O**). In a similar vein, Mo in the 1dpc nerve are transcriptionally more similar to *Chil3^+^*Mo1 than to *Ear2^+^* Mo2 in blood. This suggests that following SNC, select GC and Mo subpopulations preferentially extravasate and enter the injured nerve. Moreover, the integrated analysis clearly shows the rapid increase of *Chil3^+^*Mo in the 1dpc nerve, followed by a decline over the next six days (**Fig. 6F-6J**). Conversely, few *Gpnmb^+^* Mac are detected in blood or naïve nerve, however *Gpnmb^+^* gradually increase following SNC. Only a small number of *Gpnmb^+^* Mac is detected at 1dpc, they have increased at 3dpc, and further increased at 7dpc, suggesting these are more mature Mac (**Fig. 6K-6O**).

**Figure 6.**
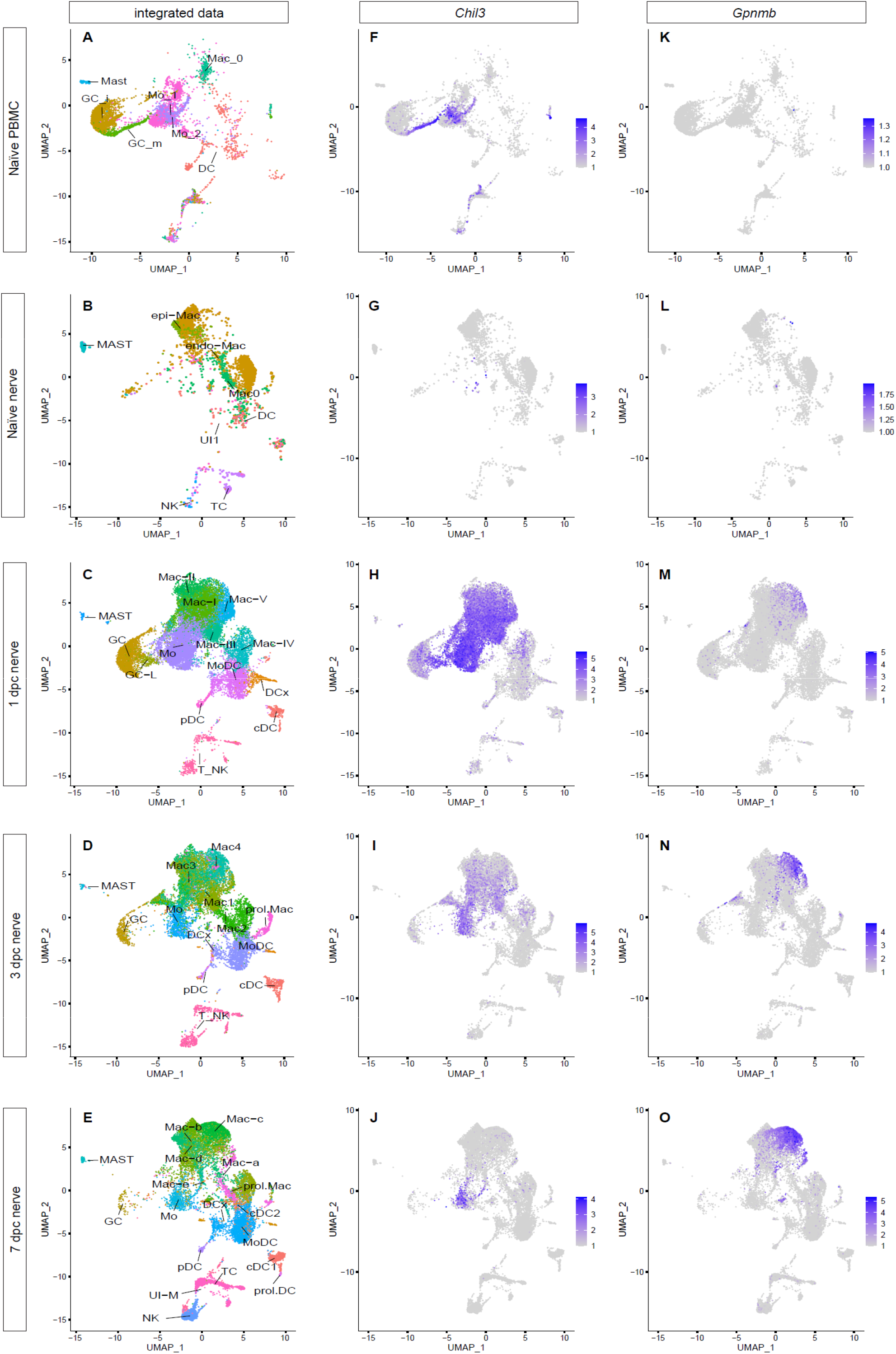
Integrated analysis of single cell transcriptomes generated from PBMC, naïve nerve, and injured nerve. (**A-E**) UMAP plots of integrated myeloid cells split into (**A**) PBMC (of naïve mice), (**B**) naïve sciatic nerve trunk, (**C**), 1dpc nerve, (**D**) 3dpc nerve, and (**E**) 7dpc nerve. (**F-J**) Integrated analysis of *Chil3^+^* Mo in (**F**) PBMC, (**G**) naïve nerve, (**H**) 1dpc nerve, (**I**) 3dpc nerve, (**J**) 7dpc nerve. (**K-D**) Integrated analysis of *Gpnmb^+^* Mac in (**K**) PBMC, (**L**) naïve nerve, (**M**) 1dpc nerve, (**N**) 3dpc nerve, (**O**) 7dpc nerve. For feature plots (**F-O**), expression values are projected onto the integrated UMAP with a minimum expression cutoff of 1. Abbreviations: GC, granulocytes; Mo, monocytes; Mac, macrophages: DC, dendritic cells; MAST, mast cells; T_NK, T cells and natural killer cells; UI, unidentified cells.

**Figure 6, Suppl. 1.**
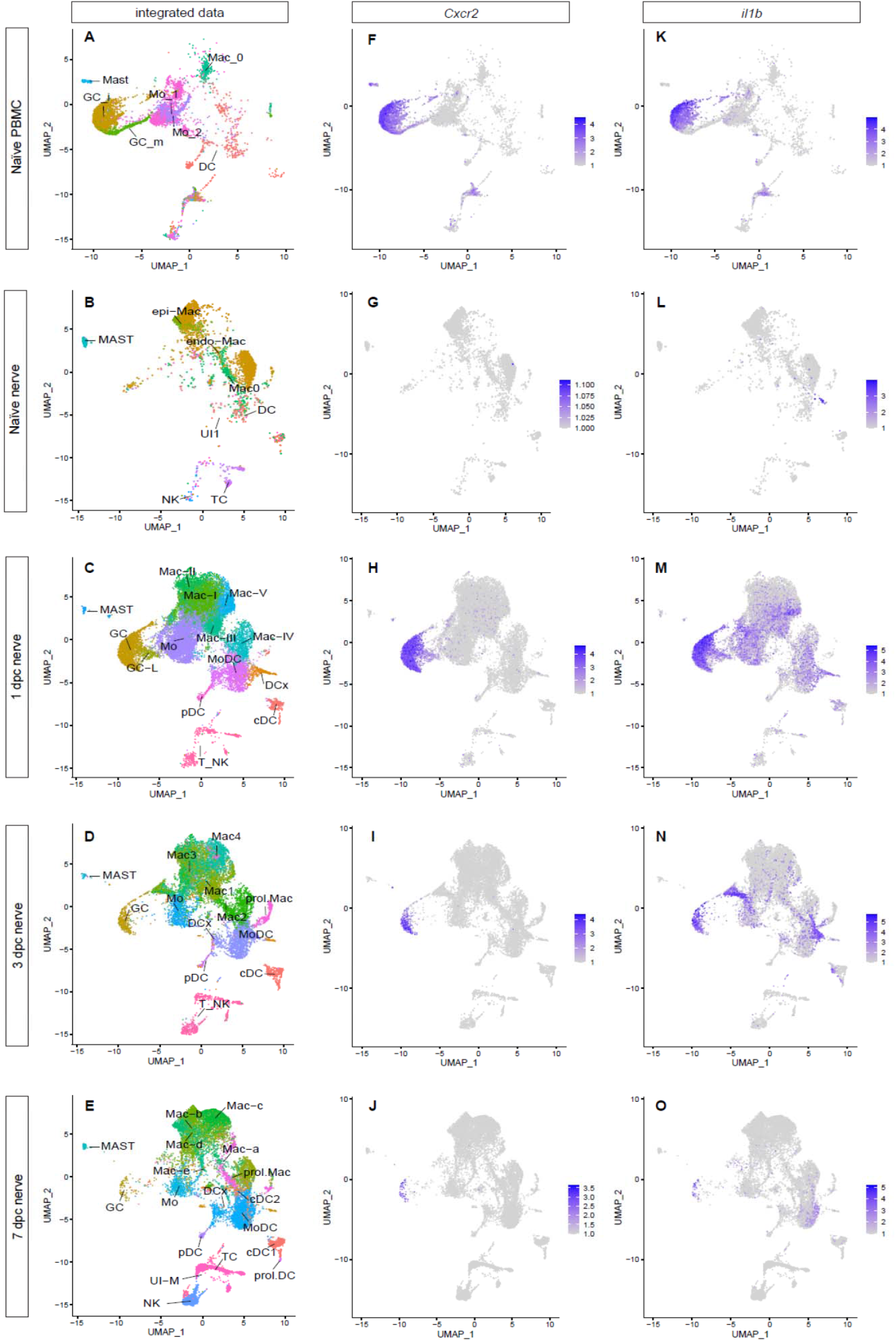
(**A-E**) UMAP plots of integrated myeloid cells split into (**A**) PBMC (of naïve mice), (**B**) naïve sciatic nerve trunk, (**C**) 1dpc nerve, (**D**) 3dpc nerve, and (**E**) 7dpc nerve. Integrated analysis to track *Cxcr2^+^* GC (**F-J**) and expression of *Il1b* (**K-O**) in myeloid cells of PBMC, naïve nerve, 1dpc, 3dpc, and 7dpc nerve. For feature plots (**F-O**), expression values are projected onto the integrated UMAP with a minimum expression cutoff of 1 Abbreviations: GC, granulocytes; Mo, monocytes; Mac, macrophages: DC, dendritic cells; MAST, mast cells; T_NK, T cells and natural killer cells; UI, unidentified cells.

To predict trajectories of Mo maturation into their descendants, Mac and MoDC, we carried out slingshot, pseudotime analysis of integrated myeloid cells. Using Mo as starting cells, revealed a bifurcated differentiation trajectory and indicates that Mo give rise to Mac and MoDC in the injured nerve **(Fig. 7; Fig. 7, Suppl. 1-3).** The high degree of Mo/Mac plasticity is evident when cell cluster identified at 1dpc (Mac-I to Mac-V) (**Fig. 2A)**, 3dpc (Mac1-Mac4) (**Fig. 2J**), and 7dpc (Mac-a to Mac-d) (**Fig. 2R**) are visualized in the integrated dataset (**Fig. 7, Suppl. 1**). To facilitate access to integrated immune cell datasets, we added these data to iSNAT.

**Figure 7.**
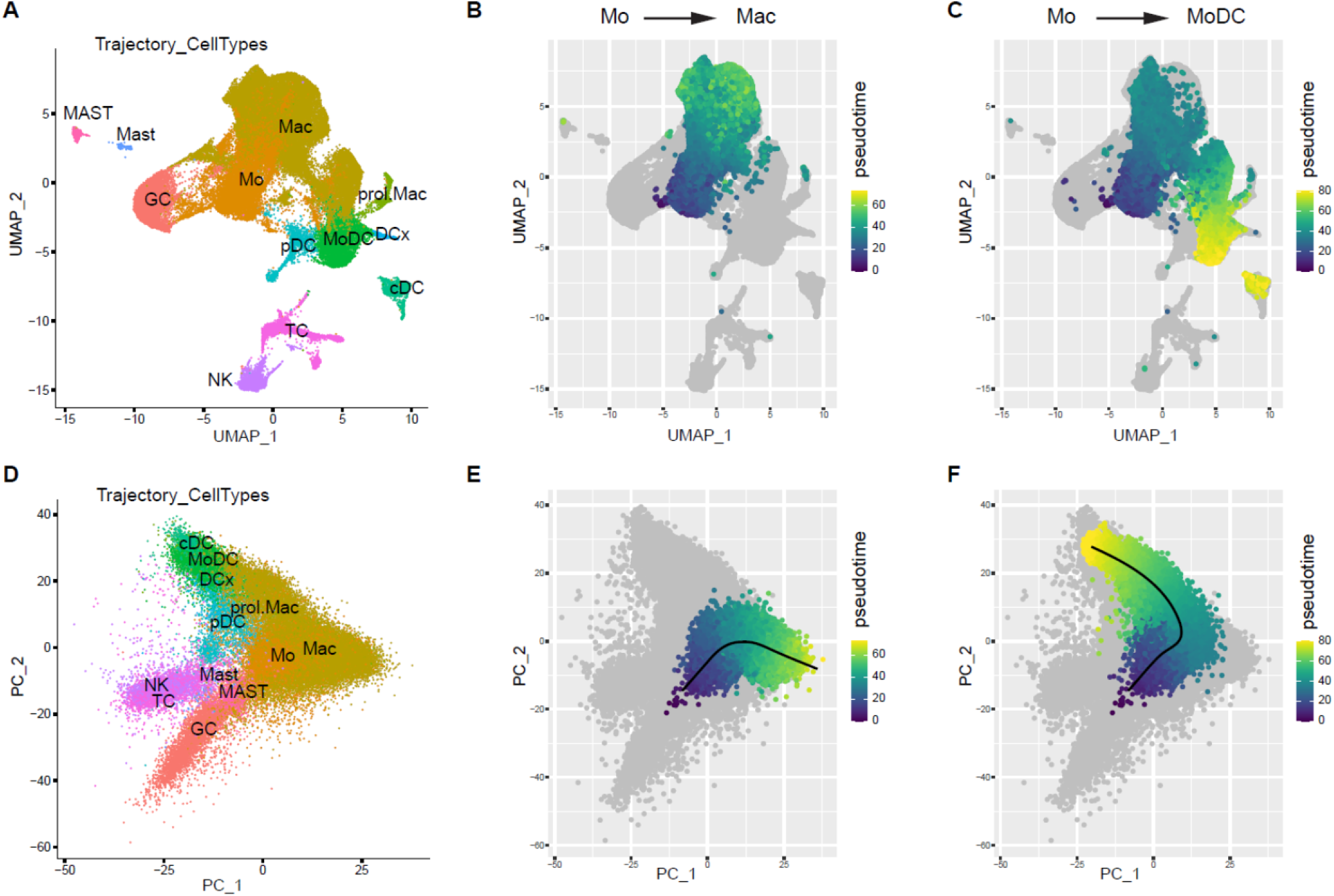
Tracking of myeloid cells before and after entering the injured nerve. **(A)** UMAP of integrated immune cells with simplified cluster labels **(D)** PCA of integrated immune cells using the same cluster labels as A. The first four principal components were used as input to slingshot pseudotime analysis (**B**) Slingshot pseudotime, for Mo to Mac trajectory, projected onto the UMAP. **(E)** Pseudotime projected onto the PCA showing the predicted trajectory, starting from Mo and differentiating toward Mac (**C**) A separate trajectory, starting from Mo shows differentiation toward MoDC; psuedotime projected on UMAP. (**F**) Mo to MoDC pseudotime projected on PCA.

**Figure 7, Suppl 1.**
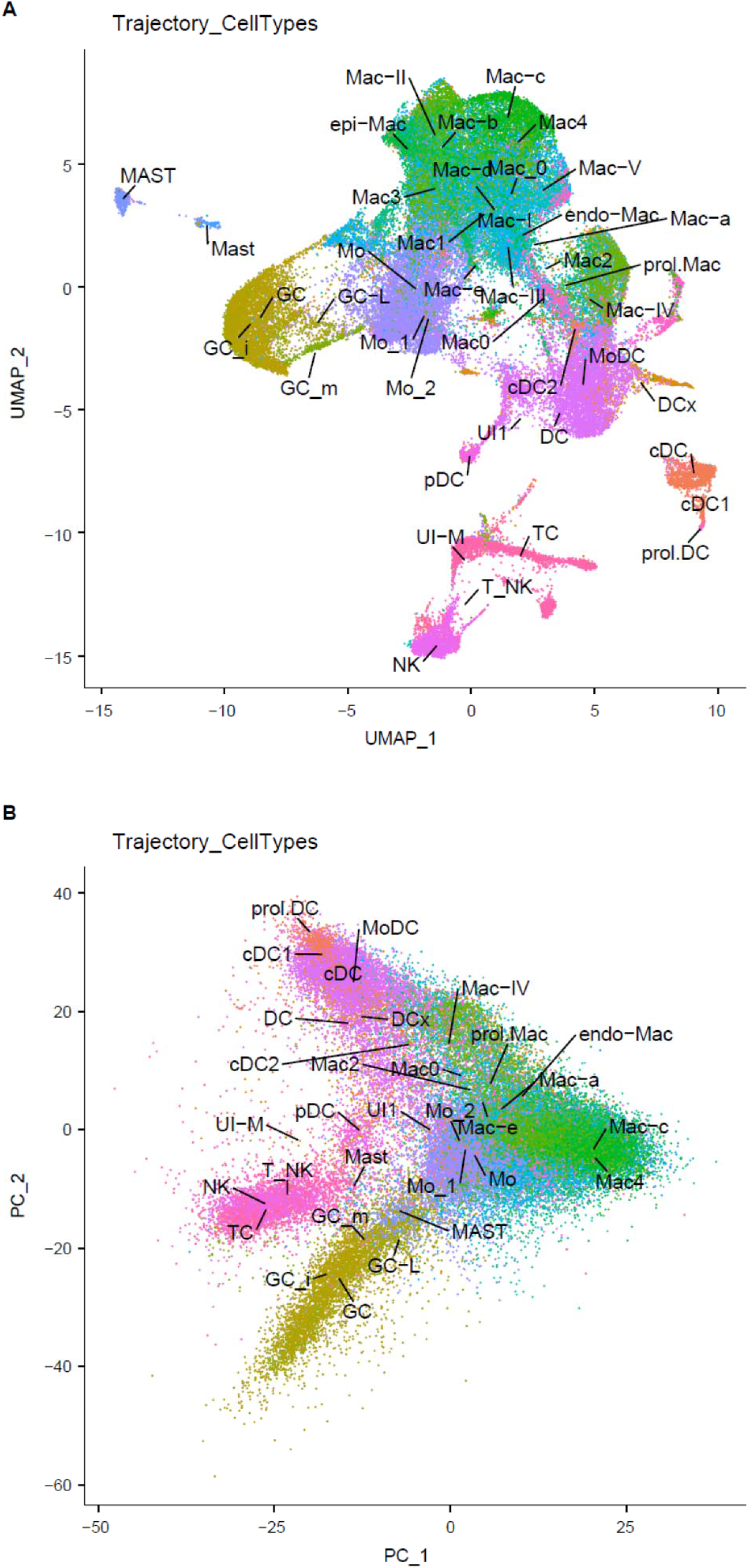
(A) Integrated immune cells UMAP with all cluster labels from each time points individual analysis. (B) Integrated immune cells PCA with all cluster labels from each time points individual analysis. Abbreviations: GC, granulocytes [GC-i and GC-m (blood), GC-L (1dpc nerve), GC (1dpc, 3dpc and 7dpc nerve)], Mo, monocytes (Mo_1 and Mo_2 (blood), Mo (1dpc, 3dpc, and 7dpc nerve)], Mac, macrophages [endo-Mac and epi-Mac (naïve nerve), (Mac-I, Mac-II, Mac-III, Mac-IV, Mac-V (1dpc nerve), Mac0, Mac1, Mac2, Mac3, Mac4 (3dpc nerve), Mac-a, Mac-b, Mac-c, Mac-d, Mac-e (7dpc nerve)], MoDC (monocyte-derived dendritic cells), cDC (conventional dendritic cells), pDC, plasmacytoid dendritic cells, DCx (mature dendritic cells), Mast (mast cells), TC (T cells), NK (natural killer cells), UI (unidentified).

**Figure 7, Suppl 2.**
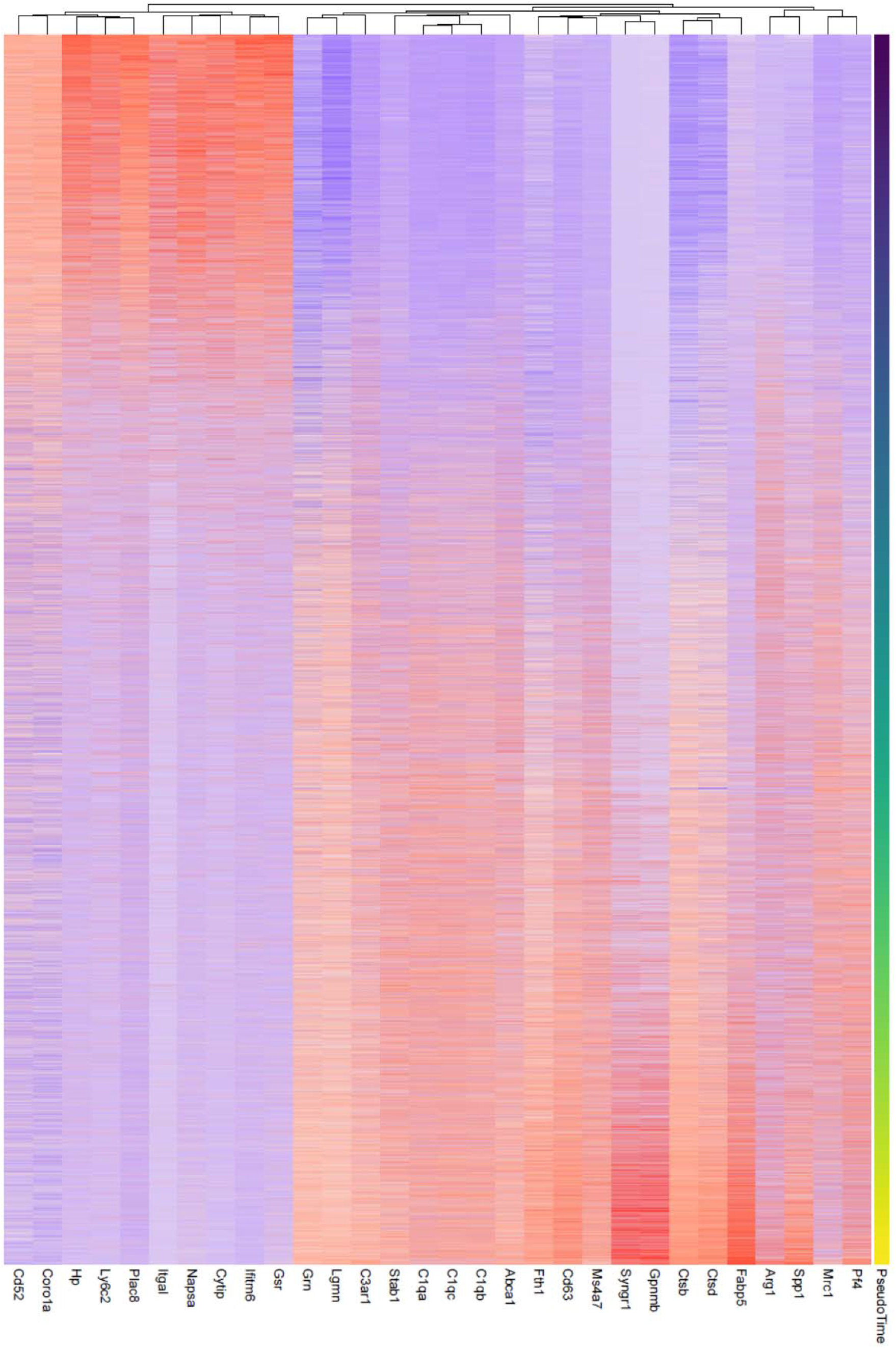
Heatmap showing the top 30 genes by importance in predicting pseudotime (Figure 7E) as determined by random forest analysis. Colors represent the Z-score calculated across genes.

**Figure 7, Suppl 3.**
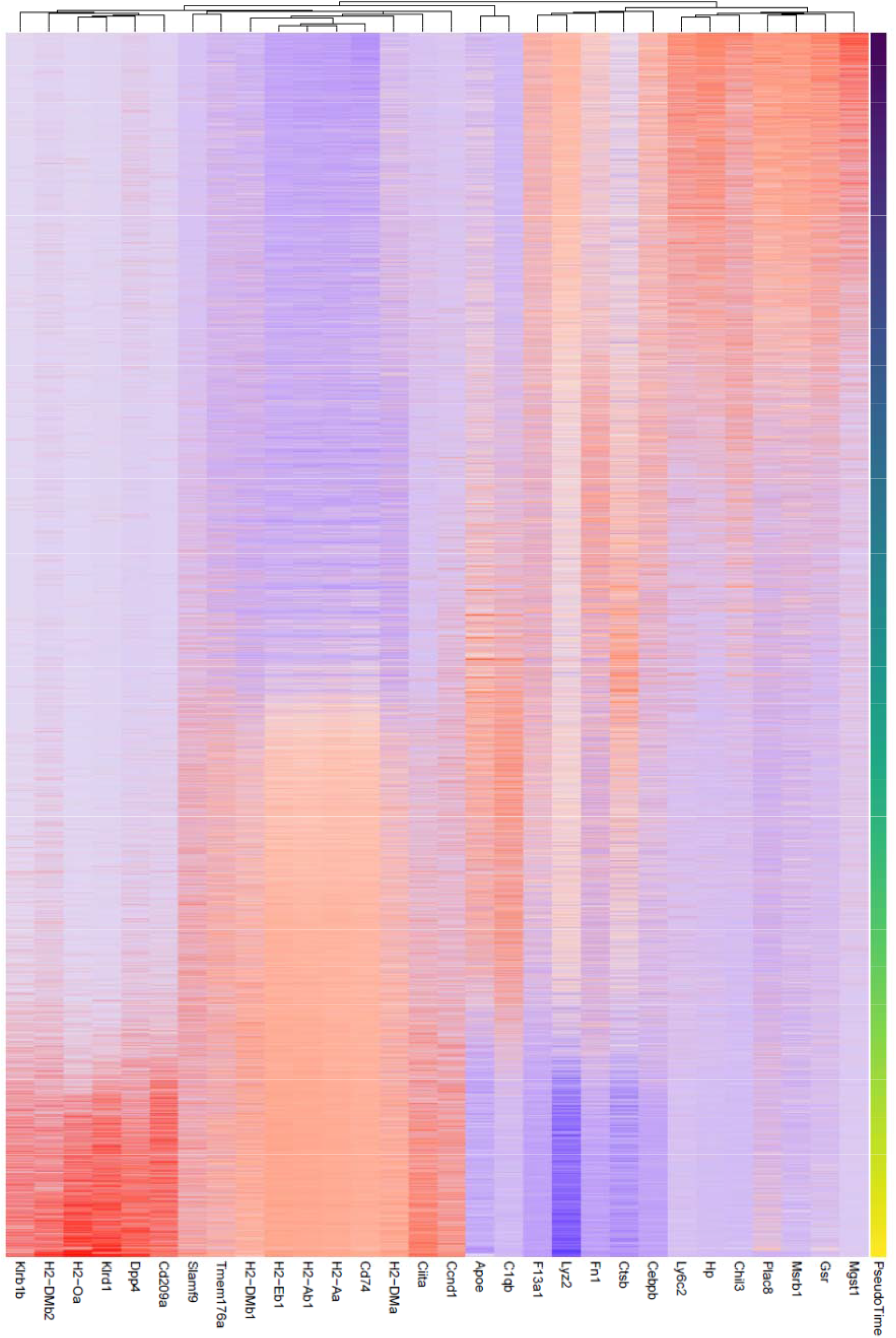
Heatmap showing the top 30 genes by importance in predicting pseudotime (Figure 7F) as determined by random forest analysis. Colors represent the Z-score calculated across genes.

### Nerve trauma generates spatial differences in immune cell composition

A crush injury divides a nerve into three distinct compartments: the proximal nerve, the injury site, and the distal nerve (**Fig. 8A**). While specialized immune cells have been identified at the site of nerve injury (Cattin et al., 2015; Kalinski et al., 2020; Shin et al., 2018), a comparative analysis of cells at the injury site and the distal nerve has not yet been carried out. Here we harvested sciatic nerves at 3dpc and microdissected ∼ 3mm segments that either harbor the injury site or distal nerve. Innate immune cells were then captured with anti-CD11b and further analyzed by scRNAseq (**Fig. 8A**). The resulting UMAP plots revealed location specific enrichment of select immune cell populations (**Fig. 8B**). For example, GC (cluster 10) are more abundant at the nerve injury site, than in the distal nerve and Mo (cluster 0) are more abundant in the distal nerve (**Fig. 8C**). Most notably, is the location specific enrichment of select Mac subpopulations (**Fig. 8C**). For example, *Arg1* expressing Mac4 (cluster 2) and Mac1 (cluster 3) are enriched at the nerve injury site (**Fig. 8D, 8E**), while *Cd38^+^* Mac3 (cluster 1) are more abundant in the distal nerve (**Fig. 8F, 8G**). The number of proliferating Mac is comparable between the injury site and distal nerve (**Fig. 8C**). Similarly, the distribution of MoDC, DCx, cDC, and T/NK is comparable between the injury site and the distal nerve (**Fig. 8C**). However, NK are more abundant in the distal nerve (**Fig. 8C**).

**Figure 8.**
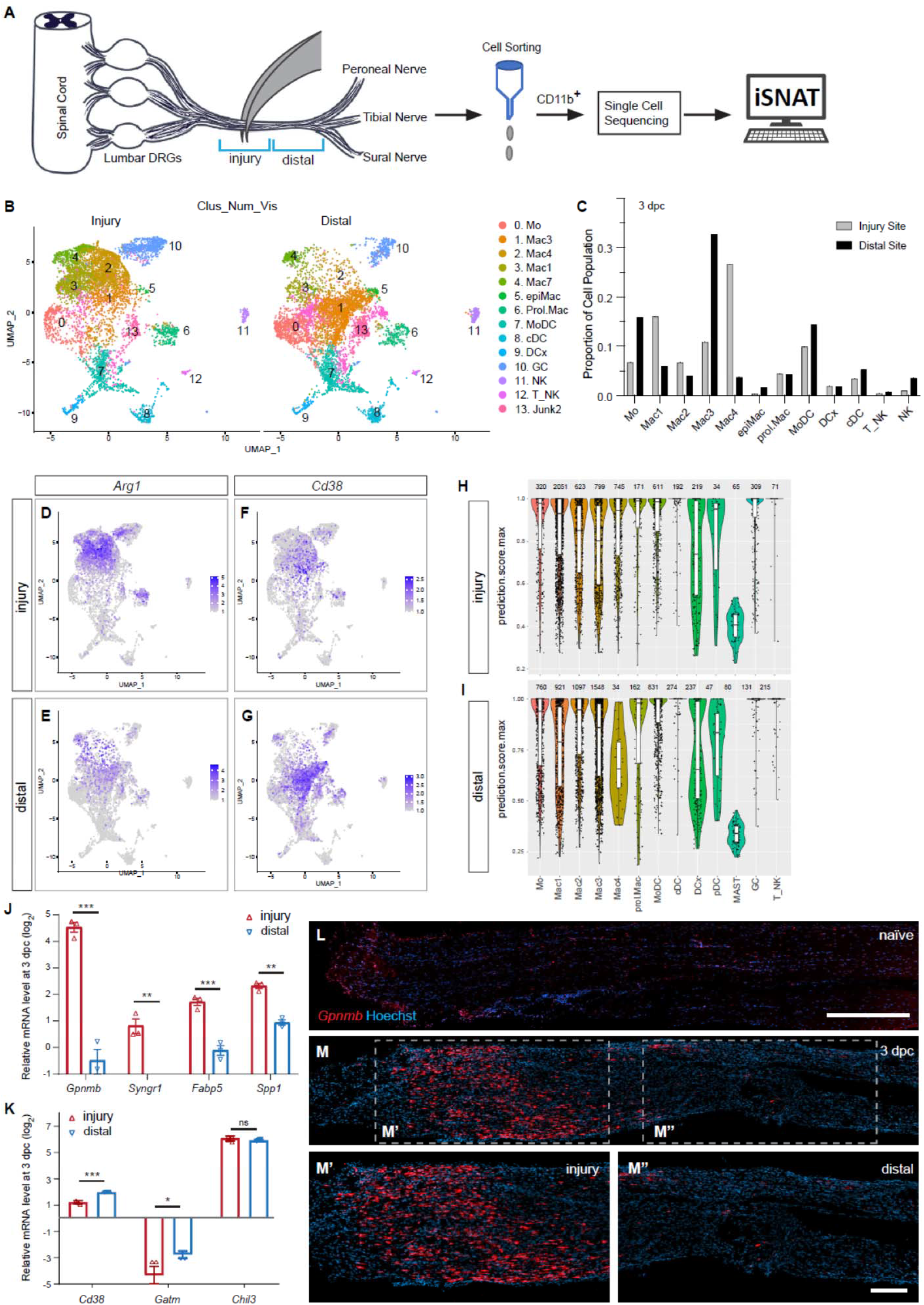
Spatial differences in the immune landscape of the injured sciatic nerve. **(A)** Cartoon of a mouse lumbar spinal cord, dorsal root ganglia (DRGs) and major branches of the sciatic nerve. Nerve segments ∼ 3 mm in length, with the injury site and distal nerve are marked with blue brackets. Immune cells from the injury site and distal nerve were captured with anti-CD11b magnetic beads and analyzed in separate scRNAseq runs. **(B)** UMAP plot of sciatic nerve myeloid cells captured at the injury site (left) and the distal nerve (right) at 3dpc. A total of 17,404 high quality cells were subjected to unsupervised Seurat-based clustering resulting in 13 cell clusters. (**C**) Bar graph of population size at the injury site versus distal nerve for 3d injured nerve immune cells. (**D, E**) Feature plots for *Arg1* expressing immune cells at the 3d injury site and distal nerve (**F, G**). Feature plots for *Cd38* expressing immune cells at the injury site and distal nerve. Expression levels are color coded and calibrated to average gene expression. (**H, I**) Projection of the 3d “whole nerve” reference data onto cells at the 3d injury site (**H**) and 3d distal nerve (**I**) onto 3d “whole nerve” reference data. The y-axis shows the prediction score for each cell’s top predicted cell population. The number of cells identified to each population is shown on top. (**J, K**) Quantification of gene expression by qRT-PCR in the 3d injured nerve injury site versus distal nerve (n= 3). P-values, * <0.05, ** <0.001, *** < 0.0001, Student’s *t* test. ns, not significant. (**L**) Longitudinal sections of naïve sciatic nerve stained for *Gpnmb* expression by RNAscope. (**M**) Longitudinal sections of 3d injured nerve stained for *Gpnmb* expression by RNAscope, proximal is to the left. Sections were counterstained with Hoechst. High power images of injury site (**M’**) and distal nerve (**M’’**) are shown. Scale bar: 200 μm (**L**, **M’’**).

For an unbiased cell cluster identification at the injury site and in the distal nerve, we compared the scRNAseq datasets generated from the 3dpc injury site and distal nerve to the “whole nerve” 3dpc reference scRNAseq data (**Fig. 2J**). We projected the whole nerve principal component analysis (PCA) structure onto “query” scRNA-seq data generated from the injury site of distal nerve, implement through Seurat v3 (Stuart et al., 2019). Our ‘TransferData’ pipeline finds anchor cells between the 3dpc whole nerve reference data and the query dataset, then uses a weighted vote classifier based on the known reference cell labels to yield a quantitative score for each cell’s predicted label in the query dataset. A prediction score of 1 means maximal confidence, all votes, for the predicted label, and a score of 0 means no votes for that label (**Fig. 8H, 8I**). Most notable is the strong enrichment of Mac4 at the 3dpc injury site (**Fig. 8H**), when compared to distal nerve (**Fig. 8I**). Similarly, distribution of the Mac1 population is skewed toward the injury site, however to a lesser extent than Mac4. Conversely, Mac3 cells are enriched in the distal nerve (**Fig. 8H,8I**). To confirm the location specific distribution of different Mac subpopulations at 3dpc, we analyzed the injury site and distal nerve for Mac subpopulation enriched transcripts using qRT-PCR. Because *Gpnmb/*glycoprotein non-metastatic melanoma protein B, and *Syngr1*/synaptogyrin 1*, Fabp5/*fatty binding protein 5, and *Spp1* are highly enriched in Mac4, we assessed their expression by qRT-PCR and found significantly higher expression at the injury site (**Fig. 8J**). Conversely, scRNAseq reveled preferential expression of *Cd38*/ADP-ribosyl cyclase 1 in Mac3 in the distal nerve and this was independently confirmed by qRT-PCR (**Fig. 8K**). Expression of the Mo marker *Chil3* is not significantly different between the injury site and distal nerve. Compared to naïve nerve, *Gatm* (creatine biosynthesis) is reduced in the nerve at 3dpc, both at the injury site and the distal nerve (**Fig. 8K**).

To assess the spatial distribution of Mac4 in the 3dpc nerve, we used *in situ* hybridization with an RNAscope probe specific for *Gpnmb.* Very few *Gpnmb^+^* cells are detected in naïve sciatic nerve (**Fig. 8L**), however at 3dpc, *Gpnmb^+^* cells are enriched in the endoneurium at the site of nerve injury site and far fewer labeled cells are detected in the distal nerve (**Fig. 8M-M’’**). Together, these studies reveal spatial differences in Mac subpopulation distribution within the injured nerve.

### Nerve trauma causes a strong inflammatory response independently of Wallerian degeneration

To separate the immune response to mechanical nerve wounding from the immune response to WD, we employed *Sarm1-/-* mice, in which WD is greatly delayed (Osterloh et al., 2012). WT and *Sarm1-/-* mice were subjected to SNC and nerves harvested at 1, 3, and 7dpc. Longitudinal nerve sections were stained with anti-F4/80 to assess abundance and distribution of Mac proximal to the injury site, at the injury site, and distal to the injury site (**Fig. 9A**). In naive WT and *Sarm1-/-* mice, few F4/80^+^ Mac are detected (**Fig. 9B,9C**). Following nerve crush injury, WT and *Sarm1-/-* mice show a rapid increase in F4/80^+^ Mac at the nerve injury site at 1dpc (**Fig. 9D,9E**), 3dpc (**Fig. 9F,9G**) and, 7dpc (**Fig. 9H,9I**). In the distal nerve, however *Sarm1-/-* mice show far fewer F4/80^+^ cells at 3dpc and 7dpc (**Fig. 9F-9I**). At 7dpc, fluoromyelin staining of the proximal WT and *Sarm1-/-* nerves shows intact myelin (**Fig. 9J, 9K**). Myelin ovoids emanating from disintegrated myelinated axons are observed in a ∼3mm segment around the site of nerve injury, both in WT and *Sarm1-/-* mice (**Fig. 9L,9M**). In the distal nerve of the same mice, myelin ovoids are observed only in WT (**Fig. 9N**), but not *Sarm1-/-* mutants showing that at 7dpc fiber fragmentation has not yet occurred (**Fig. 9O).** For an independent assessment of WD elicited nerve inflammation, nerve trunks were isolated from naïve and injured WT and *Sarm1-/-* mice, divided into proximal nerve, the injury site, or distal nerve, and analyzed by Western blotting (**Fig. 9P**). Independently of *Sarm1* genotype, the injury sites show elevated levels of CD11b compared to proximal nerve. In the distal nerve, CD11b was more abundant in WT than in *Sarm1-/-* mice.

**Figure 9.**
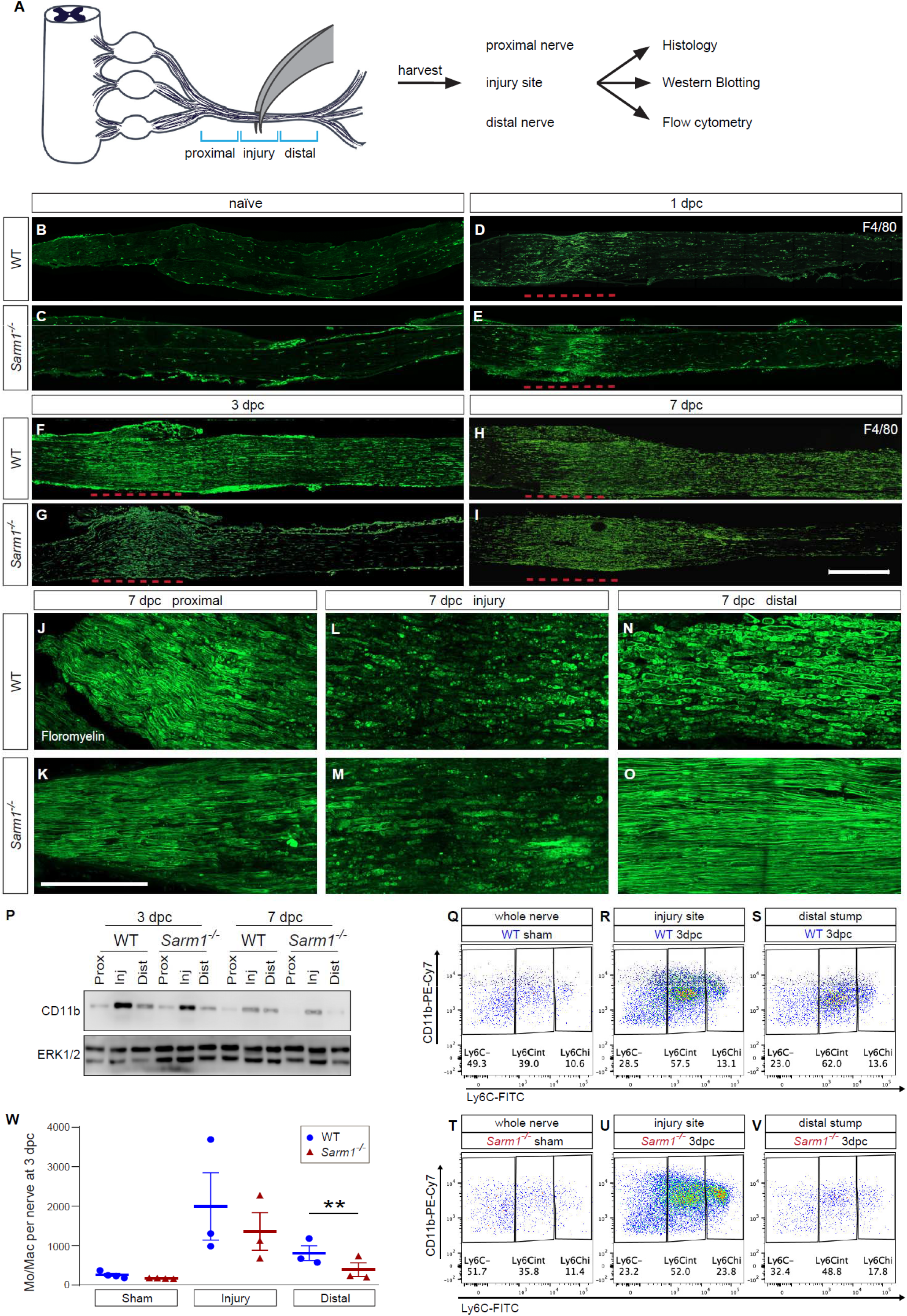
Nerve trauma causes WD independent nerve inflammation. Cartoon of a mouse lumbar spinal cord, dorsal root ganglia (DRGs) and major branches of the sciatic nerve. A nerve injury divides the nerve trunk into a proximal segment, the injury site, and distal segment, each marked with blue brackets. Nerve segments were harvested and subjected to analysis. (**B-I**) Longitudinal sciatic nerve sections from WT and *Sarm1^-/-^* mice, stained with anti-F4/80 for identification of Mac. Representative examples of (**B, C**) naïve nerve, (**D, E**), 1dpc (**F, G**), 3dpc, and at (**H, I**) 7dpc. Injury site is marked with a dashed red line, proximal is to the left. Scale bar, 500 µm. (**J-O**) Longitudinal sciatic nerve sections from WT and *Sarm1^-/-^* mice at 7dpc, stained with fluoromyelin. Representative images of proximal nerve, the injury site and distal nerve are shown. Scale bar, 200 µm. **(P)** Western blots of sciatic nerve segments collected at 3dpc and 7dpc from WT and *Sarm1^-/-^*mice. Nerves were divided into proximal, injury site, and distal segments and blots probed with anti-CD11b, and anti-ERK1/2. **(Q-V)** Flow cytometry dotplots for Mo/Mac of sham operated WT and *Sarm1^-/-^* sciatic nerve trunks, the 3dpc nerve injury site and distal nerve. (**W**) Quantification of Mo/Mac (Ly6C^hi^ + Ly6C^int^ + Ly6C^-^) in sham operated mice, the 3dpc injury site and 3dpc distal nerve of WT and *Sarm1^-/-^* mice. N= 3, with 3-5 mice per genotype per replica. Flow data are represented as mean ± SEM. Statistical analysis was performed in GraphPad Prism (v9) using two-way, paired *t*-test. ** p<0.01.

To quantify immune cell profiles in WT and *Sarm1-/-* mice, we used flow cytometry (**Fig. 9Q-9W)**, the gating strategy is illustrated in **Fig. 9, Suppl. 1**. In sham operated mice, no significant differences in Ly6C-high (Ly6C^hi^), Ly6C-intermediate (Ly6C^int^), or Ly6C-negative (Ly6C^-^) Mo/Mac were observed (**Fig. 9Q,9T**). For quantification of immune cell profiles that respond to traumatic nerve wounding versus WD, we separately harvested the site of nerve injury and distal nerve for analysis by flow cytometry. At the 3dpc injury site, the total number of Mo/Mac is comparable between WT and *Sarm1-/-* mice (**Fig. 9R,9U,9W**). However, within the distal nerve, significantly more Mo/Mac cells are present in WT mice than in *Sarm1-/-* mice (**Fig. 9S,9V,9W**). Taken together, these studies show that nerve trauma causes a highly inflamed wound microenvironment, independently of WD, and a distinct inflammatory response in the distal nerve, that is WD dependent.

**Figure 9, Suppl. 1.**
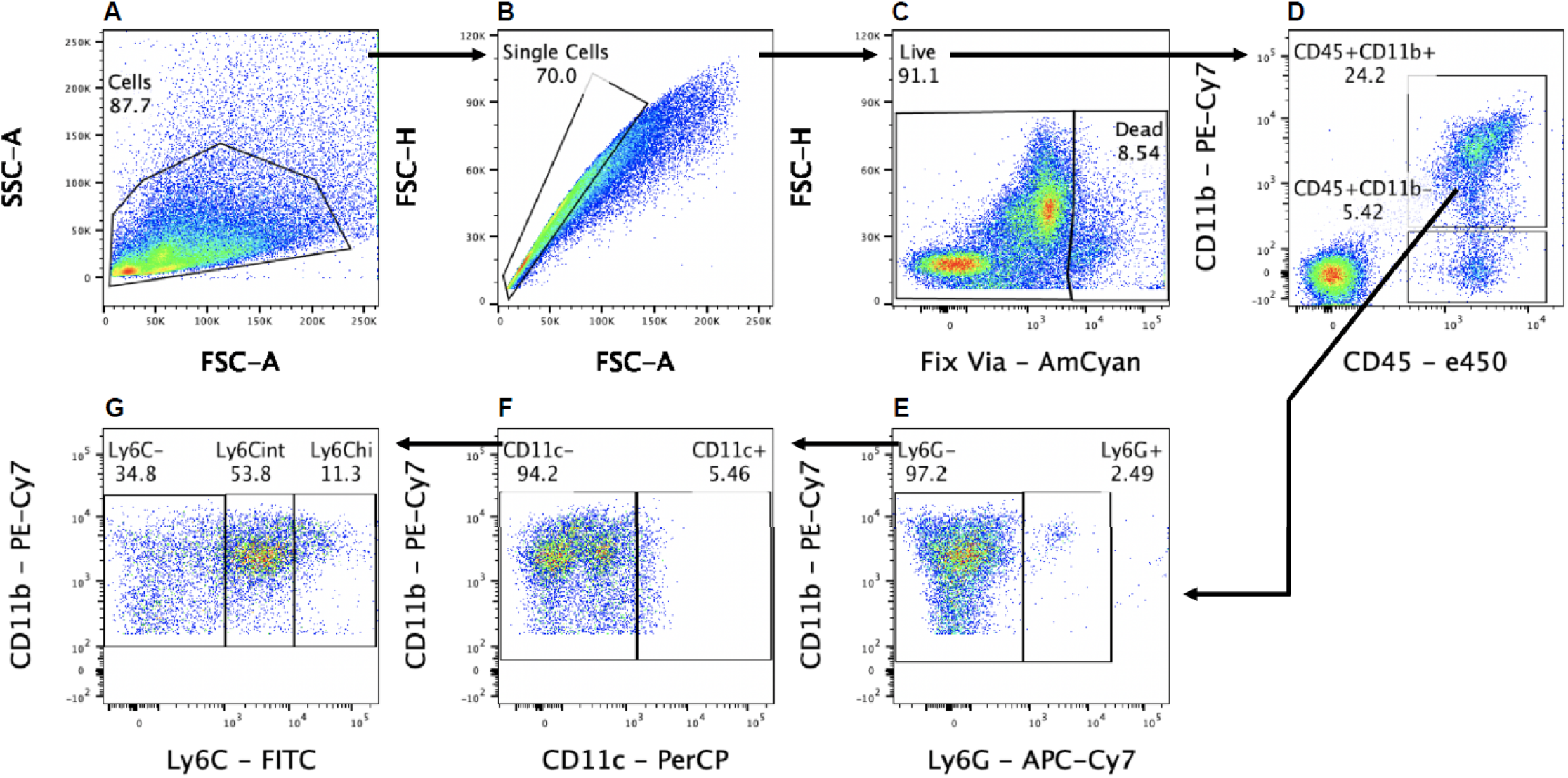
Gating strategy for flow cytometry. **(A)** Cells were first gated with forward scatter (FSC-A) and side scatter (SSC-A) to exclude debris. **(B)** Cells were then gated with forward scatter height (FSC-H) and FSC-A to find single cells and to exclude doublets. **(C)** Live cells were isolated by negative staining for fixed viability dye (Fix Via). **(D-G)** Leukocytes (D) were analyzed as follows: lymphocytes were isolated as CD45^+^, CD11b^-^. Myeloid cells (CD45^+^, CD11b^+^) were further separated into Ly6G^+^ granulocytes (E). The remaining cells (CD45^+^, CD11b^+^, Ly6G^-^) were characterized as DC (F) (CD45^+^, CD11b^+^, CD11c^+^, Ly6G^-^), and Mo/Mac (G) (CD45^+^, CD11b^+^, CD11c^-^, Ly6G^-^).

### In *Sarm1-/-* mice, monocytes enter the distal nerve stump but fail to mature into macrophages

Because we used *Sarm1* global knock-out mice for our studies and *Sarm1* has been shown to function in Mac (Gurtler et al., 2014), a potential confounding effect is *Sarm1* deficiency in Mac. To assess nerve entry of circulating WT immune cells, in a *Sarm1-/-* background, we employed parabiosis (**Fig. 10A**). *Sarm1-/-* host mice were surgically fused to tdTomato (tdT) donor mice and allowed to recover for three weeks. For comparison, WT/tdT parabiosis complexes were generated and processed in parallel. Flow cytometry was used to analyze blood samples of host parabionts for tdT^+^ leukocytes and revealed ∼30% chimerism (**Fig. 10B, 10C**). In each complex, both parabionts were subjected to bilateral nerve crush. At 7dpc, analysis of longitudinal nerve sections of the *Sarm1-/-* parabiont revealed that many tdT^+^ leukocytes entered the site of nerve injury, comparable to injured WT parabionts (**Fig. 10D, 10E, 10F**). In the distal nerve of the *Sarm1-/-* parabiont, at 1mm, 2mm, and 3mm distal to the injury site, some tdT^+^ leukocytes are present, however at significantly reduced numbers when compared to WT parabionts (**Fig. 10F**). Interestingly, only few tdT^+^ cells in the *Sarm1-/-* distal nerve stained for F4/80, a marker for Mac (**Fig. 10G,10H**) or Ly6G, a marker for neutrophils (**Fig. 10, Suppl. 1**). To further investigate the blood-borne immune cells that enter the 7dpc distal nerve of *Sarm1-/-* mice, we separately harvested and analyzed the 7dpc injury site and distal nerve segments from WT and *Sarm1-/-* single mice using flow cytometry. The abundance of Mo and Mac, identified as Ly6C^hi^, Ly6C^int^, and Ly6C^-^ cells, at the nerve injury site is comparable between WT and *Sarm1-/-* mice (**Fig. 10I, 10J**). In the distal nerve however, Ly6C^hi^ cells in *Sarm1-/-* nerves are significantly elevated compared to Ly6C^int^ and Ly6C^-^ populations (**Fig. 10I,10J**). This stands in contrast to WT distal nerves, where Ly6C^-^ and Ly6C^int^ cells outnumber the Ly6C^hi^ population **(Fig. 10K).** Because WT and *Sarm1-/-* mice show similar baselines of Mac in the naïve nerve (**Fig. 9Q,9T**), this shows that in the absence of WD, SNC is sufficient to trigger a strong immune response to nerve wounding, but fails to elicit WD associated nerve inflammation, except for the appearance of Mo.

**Figure 10.**
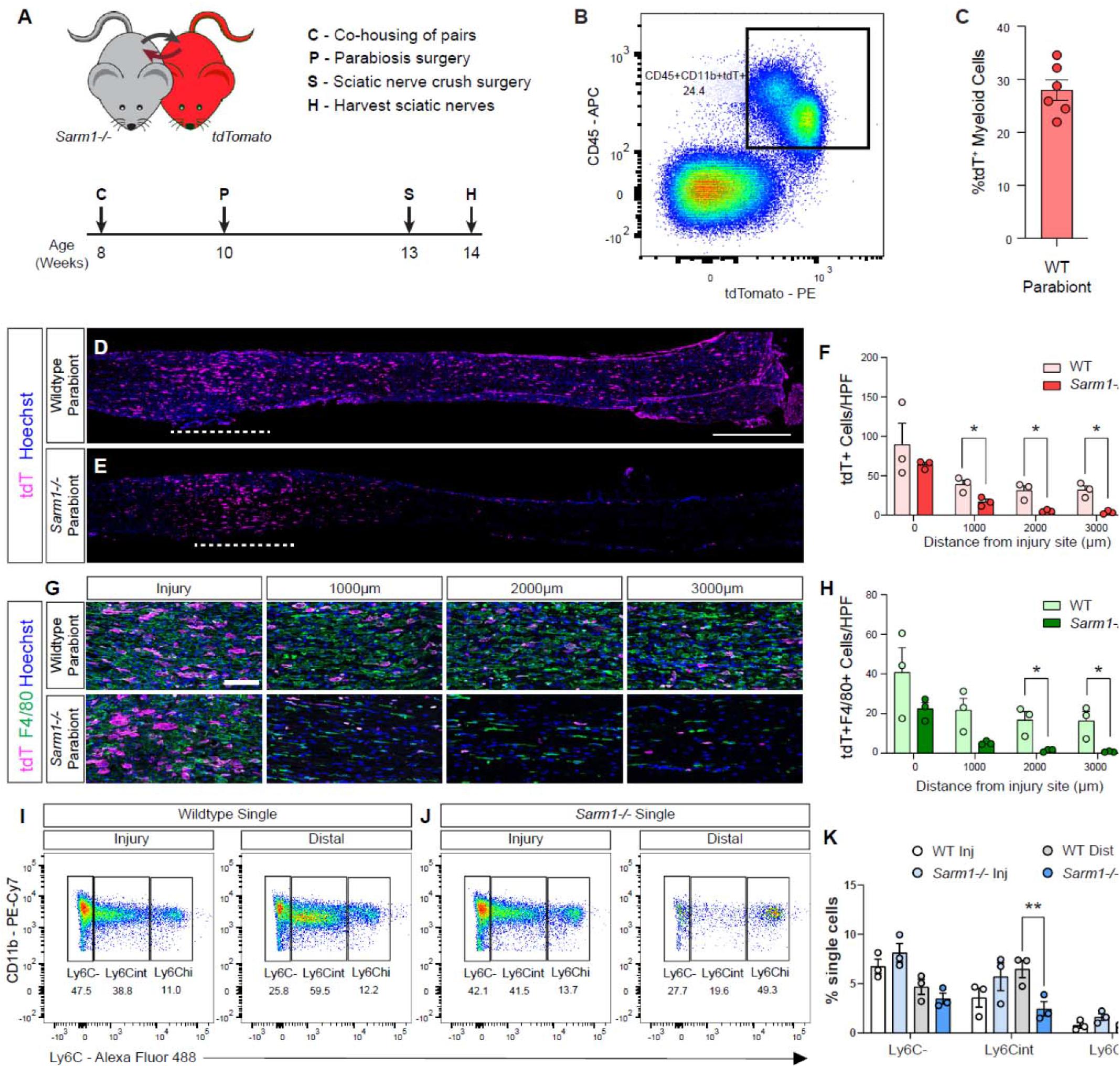
Evidence for WD dependent and WD independent nerve inflammation. (**A**) Timeline for parabiosis experiments. After a two-week co-housing period, 10-week-old WT or *Sarm1^-/-^* and - *tdTomato* mice were surgically paired. (**B**) To assess chimerism, blood was harvested and analyzed by flow cytometry. Dotplot of tdT^+^ myeloid cells (CD45^+^CD11b^+^) is shown. (**C**) Quantification of tdT^+^ myeloid cells in host parabionts (n= 6), revealed chimerism of 28± 2%. (**D, E**) Bilateral SNC was performed 3 weeks after pairing and tissue harvested at 7dpc. Longitudinal sciatic nerve sections from (**D**) WT and (**E**) *Sarm1^-/-^*parabionts showing infiltrating tdT^+^ leukocytes (magenta). Nuclei (blue) were labeled with Hoechst dye. The nerve crush site is marked by the white dashed line, proximal is to the left. Scale bar, 500 μm. (**F**) Quantification of tdT^+^ cells per high power field (HPF, 500 μm x 250 μm) at the injury site (0 μm) and at 1000, 2000, and 3000 μm distal to the injury site. The average cell number ± SEM is shown, n = 3 mice per genotype, average of 4 HPF per 2 nerves. Student’s *t* test, p < 0.05 (*). (**G**) HPF of sciatic nerves from WT and *Sarm1^-/-^* parabionts 7dpc taken from the injury site, 1000, 2000, and 3000 μm distal to the injury site showing infiltrating tdT^+^ leukocytes (magenta), F4/80^+^ macrophages (green), and nuclei (blue). Scale bar, 100 μm. (**H**) Quantification of tdT^+^F4/80^+^ cells per HPF ± SEM at indicated distances distal to the injury site, n = 3 mice, average of 4 HPF per 2 nerves. Student’s *t* test, p < 0.05 (*). (**I, J**) Flow cytometric analysis of sciatic nerves from single (not part of parabiosis complex) WT and *Sarm1^-/-^* mice 7dpc. Sciatic nerve trunks were microdissected and separated in 3 mm injury site and distal nerve segments. Dotplots showing Mo/Mac maturation assessed by Ly6C surface staining, Mo (Ly6C^hi^), Mo/Mac (Ly6C^int^), Mac (Ly6C^-^), previously gated as CD45^+^CD11b^+^Ly6G^-^CD11c^-^ cells. (**K**) Quantification of Mo/Mac shown in panels **I** and **J,** as a percentage of single cells ± SEM, n = 3, injury and distal sites were pooled from 5 mice per genotype per biological replicate. Two-way ANOVA with Tukey’s post-hoc test for multiple comparisons, p < 0.05 (*), p < 0.01 (**).

**Figure 10, Suppl. 1.**
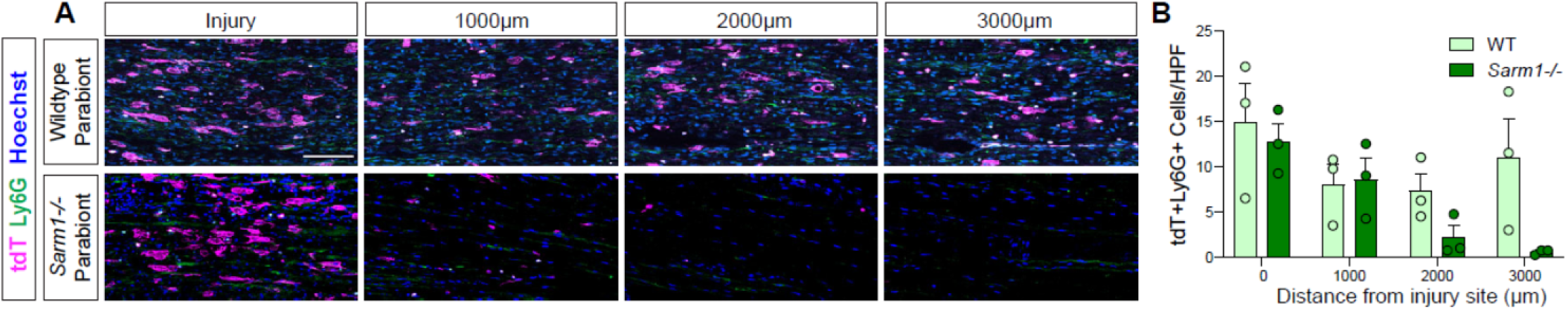
Quantification of tdT,Ly6G double-positive neutrophils in the 7dpc nerve of host parabionts (n= 6). (**A**) High power filed (HFP) of longitudinal sections of a WT and *Sarm1-/-* parabionts at 7dpc. Representative images taken from the injury site, 1000, 2000, and 3000 μm distal to the injury site. Infiltrating tdT^+^ leukocytes (magenta), Ly6G^+^ neutrophils (green), and nuclei (Hoechst) are labeled. Scale bar, 100 μm. (**B**). Quantification of tdT^+^Ly6G^+^ cells per (HPF) ± SEM at indicated distances distal to the injury site, n = 3 mice, an average of 4 HPF per 2 nerves. Student’s *t* test.

Taken together, these data show that in *Sarm1-/-* mice Ly6C^hi^ Mo enter the distal nerve prior to WD but fail to differentiate into Mac. This suggests that chemoattractive signals for Mo are released from severed axons prior to WD and that fiber degeneration in the distal nerve is required for Mo maturation. Moreover, WD is required for rapid, full-blown inflammation of the distal nerve.

## Discussion

The injured adult murine PNS exhibits a remarkable degree of spontaneous axonal regeneration and functional recovery. To better understand the cellular and molecular events associated with PNS repair, we carried out a longitudinal scRNAseq study. Analysis of the immune response to nerve crush injury, during the first week, revealed a highly dynamic microenvironment. The early immune response is pro-inflammatory and dominated by GC and Mo/Mac, metabolically programmed for glycolytic energy production. The elevated expression of nearly all glycolytic enzymes, lactate dehydrogenase, and the lactate export channel MCT4 indicates that a Warburg-like effect is at play, coupling glycolytic energy production with a proinflammatory Mo/Mac phenotype. This stands in marked contrast to the low glycolytic activity of circulating Mo/Mac in blood of naïve mice and indicates a rapid metabolic shift upon nerve entry. The glycolytic burst is short-lived, however, at 3dpc expression of glycolytic enzymes begins to decline, and at 7dpc has reached levels similar to naïve nerve. As glycolytic activity declines, there is evidence for a metabolic shift toward OXPHOS, and this coincides with the appearance of Mac with a resolving phenotype. Separation of the nerve injury site from distal nerve revealed that mechanical nerve injury creates two separate immune microenvironments. A wound repair response at the crush site, and an inflammatory response to WD in the distal nerve. This was further corroborated in injured *Sarm1-/-* mice. At the injury site of *Sarm1-/-* and WT mice, nerve crush results in a strong immune response, dominated by hematogenous leukocytes. In the distal nerve of *Sarm1-/-* mice, full-blown nerve inflammation is delayed, and thus, WD dependent. Hematogenous immune cells are largely missing from the *Sarm1-/-* distal nerve, except for Mo, suggesting that chemotactic signals are released from severed fibers prior to physical disintegration. Taken together, we describe a framework of cell types and single-cell transcriptomes for a neural tissue with a high degree of spontaneous morphological and functional regeneration. The datasets reported provide an essential step toward understanding the dynamic nature of complex biological processes such as neural tissue degeneration and regeneration.

### iSNAT

To facilitate mining of scRNAseq datasets of naïve nerve and at different post-SNC time points, we generated the injured sciatic nerve atlas (iSNAT). The “*Expression Analysis*” function is a search tool to assess which cells in the nerve express a gene of interest. The output is four feature pots (naïve, 1, 3, and 7dpc nerve) shown side-by-side to assess which cells express a gene of interest and whether the gene is regulated by SNC. In addition, any cell type or time point during the first week can be selected to identify the top 50 enriched gene products. The *two Genes* function quantifies co-expression of two genes in the same cell. Higher resolution UMAP plots of select cell types, such as immune cells and structural cells (stromal cells), can be accessed and mined separately for analysis of subcluster specific gene expression. We added single cell transcriptomes of PBMC and used dataset integration to show how immune cell clusters change during the first week following nerve injury. Embedded in iSNAT is the CellChat function, designed for identification of intercellular signaling networks. Families of surface or secreted molecules, e.g. CXCL family members, can be searched for cells in the naïve and injured nerve that express the corresponding receptor(s) and the probability for this interaction to occur is calculated. This provides a powerful tool for understanding interactions among different cell types in the nerve. The *Spatial Distribution* function shows gene expression at the injury site versus distal nerve at 3dpc. We have validated many gene products identified by scRNAseq, using a combination of qRT-PCR, RNAscope, immunofluorescence staining, reporter gene expression, and ELISA. While iSNAT is expected to facilitate data analysis and functional studies in the injured PNS, there are some notable limitations. The reading depth at the single cell level is still limited. We have sequenced and analyzed ∼150,000 high-quality cells. On average we detect ∼2000 unique features per cell, well below the estimated total of 5000-10000. Thus, if a gene of interest is not found in iSNAT, it may either not be expressed, or be expressed below the detection sensitivity of scRNAseq. We acknowledge that enzymatic tissue digestion combined with mechanical tissue dissociation may lead to cell loss or variable capturing efficiency for different cell types. Most notably, mSC are sparse in our naïve nerve dataset. Additional cell types that may be lost include B cells and adipocytes, both of which have previously been detected in the rodent PNS (Chen et al., 2021). Our gene expression atlas is work in progress and we anticipate that future studies will overcome these limitations, allowing us to generate improved, next generations of iSNAT.

### Structural cells in the injured nerve shape the immune microenvironment

A nerve crush injury triggers proliferation of stromal cells, including epineurial Fb, identified as *Mki67^+^Pdgfra^+^* cells that mature into Fb. dMES are abundantly found in the injured nerve. In addition, pMES and eMES begin to proliferate following SNC. In the UMAP plot of 3dpc nerve, clusters with proliferating structural cells are connected and give rise to pMES, eMES, and dMES (**Fig. 2**). This suggests that in addition to damaging the epineurium, SNC damages protective cell layers within the nerve, including the perineurium, a thin cell layer of epithelioid myofibroblasts that surrounds nerve fascicles, and the endoneurium, a delicate layer of connective tissue that covers individual myelinated nerve fibers and contains the endoneurial fluid. Proliferation of eMES and pMES likely reflects damage to the BNB and is supported by the accumulation of some serum proteins in the crushed nerve. In addition to IgG (Vargas et al., 2010), we detected other serum proteins that function in opsonization. These include complement components, soluble C1qR1/CD93, adiponectin (Adipoq), and pentraxins (Nptx2, Crp), molecules that aid in the clearance of cellular debris and apoptotic cells and push Mac toward an anti-inflammatory resolving phenotype (Blackburn et al., 2019; Casals et al., 2019; Guo et al., 2012).

Of interest, structural cells in the injured nerve consistently show the highest number of unique transcripts, indicative of a strong injury response. MES are a major source of immune modulatory factors, shaping the injured nerve microenvironment in a paracrine manner through release of soluble factors. CellChat, identified important roles in chemotaxis, angiogenesis, ECM deposition and remodeling, suggestive of extensive stroma-immune cell communication. In particular, eMES are a major signaling hub and show strong interactions with Mo, Mac, SC, pMES, and EC. The presence of several chemokines, growth factors, and immune proteins identified at the transcriptional level was independently validated by ELISA, providing confidence in the quality of scRNAseq datasets.

### Cellular metabolism and macrophage functional polarization in the injured PNS

Evidence from injured non-neural tissues shows that immune cell metabolism is directly linked to cell plasticity and function, thereby affecting tissue repair and scarring (Eming et al., 2021; Eming et al., 2017). Little is known about the metabolic adaptions associated with successful neural tissue repair. Here we compared immune cell metabolism of bone-marrow derived circulating myeloid cells before and after nerve entry. Once in the injured nerve, neutrophils, Mo, and Mac undergo rapid metabolic reprogramming, greatly increasing gene products that drive glycolysis. This is similar to the metabolic shift observed in non-neural tissues with high regenerative capacities, such as skeletal muscle (Eming et al., 2021). Interestingly, rapid upregulation of glycolytic activity in myeloid cells in the injured nerve is reminiscent of the injury-regulated glycolytic shift in SC, involving the mTORC/Hif1α/c-Myc axis (Babetto et al., 2020). This suggests that enhanced glycolytic flux and lactate extrusion from both, SC and innate immune cells, is axoprotective.

Many cells use aerobic glycolysis during rapid proliferation, since glycolysis provides key metabolites for the biosynthesis of nucleotides and lipids (Lunt and Vander Heiden, 2011). In the 1dpc nerve, myeloid cells show highest levels of glycolytic enzymes, however only few *Mki67* (encoding Ki67^+^) proliferating cells are detected. In immune cells, alterations in metabolic pathways couple to immune cell effector function, most notably the production of different cytokines (O’Neill et al., 2016). Glucose is a main source for cellular energy (ATP) production through two linked biochemical pathways, glycolysis and the mitochondrial TCA (O’Neill et al., 2016; Voss et al., 2021). Glycolysis converts glucose into pyruvate and pyruvate is converted into acetyl-CoA to enter the TCA and fuel OXPHOS in mitochondria as an efficient means of ATP production. Alternatively, pyruvate can be converted into lactate and NAD^+^, creating a favorable redox environment for subsequent rounds of glycolysis. The upregulation of *Ldha* and *Slc16a3*/MCT4 in myeloid cells of the injured nerve is striking and resembles the Warburg effect described for cancer cells (Schuster et al., 2021; Zhu et al., 2015). The transient increase in extracellular lactate may not only be axon protective (Babetto et al., 2020; Funfschilling et al., 2012), but additionally regulate immune cell reprogramming (Morioka et al., 2018) and trigger pain (Rahman et al., 2016). The injury induced increase of *Hif1α* suggests that hypoxia is a main driver of metabolic reprogramming, however, the Hif1α pathway can also be activated by pattern recognition receptors recognizing DAMPs released by injured cells following trauma (Corcoran and O’Neill, 2016).

The molecular basis for Mac reprogramming into an anti-inflammatory state remains incompletely understood. A resolving Mac phenotype may be initiated by Mac mediated engulfment and digestion of apoptotic cell corpses (Boada-Romero et al., 2020; Greenlee-Wacker, 2016). Mac in the injured sciatic nerve are fully equipped with the molecular machinery for efferocytosis, including phagocytic receptors, enzymes, and transporters to cope with elevated cholesterol load and other metabolic challenges (Kalinski et al., 2020). Parabiosis experiments, combined with SNC revealed that clearing of apoptotic leukocytes, through efferocytosis, takes place in the injured PNS (Kalinski et al., 2020). Professional phagocytes that undergo multiple rounds of efferocytosis experience metabolic stress such as accumulation of intracellular lipids (Schif-Zuck et al., 2011). Growing evidence suggests that Mac leverage efferocytotic metabolites for anti-inflammatory reprogramming to promote tissue repair (Zhang et al., 2019). During the resolution phase Mac are equipped with the machinery for fatty acid oxidation and OXPHOS as a means of energy production. The rapid reprogramming of Mac is likely key for wound healing, axon regeneration, and restoration of neural function. Timely resolution of inflammation protects from excessive tissue damage and fibrosis. Interestingly, perineural cells may function as lactate sink in the injured nerve, since pMES express high levels of *Slc16a1*/MCT1 for cellular import, as well as *Ldhb* for conversion of lactate into pyruvate. This suggests that different cell types in the injured nerve employ different strategies to cover their bioenergetic needs. It will be interesting to examine whether Mo/Mac metabolic reprogramming, efferocytosis, and inflammation resolution are altered under conditions where nerve health and axon regeneration are compromised (Sango et al., 2017). Because nerve inflammation has been linked to the development of neuropathic pain (Davies et al., 2020), prolonged and poorly resolving nerve inflammation, due to impaired metabolic reprogramming, may directly contribute to pain syndromes.

### Identification of distinct immune compartments in the injured PNS

Mo/Mac are highly plastic cells that are educated by the microenvironment. Considering the complexity of injured PNS tissue, it is perhaps not surprising that Mac subpopulations were identified that are not uniformly distributed within the injured nerve. At the site of injury, nerve trauma is caused by compression or transection, resulting in cell destruction, release of damage associated molecular patterns (DAMPs), vascular damage, nerve bleeding, and disruption of tissue homeostasis. In the distal nerve, where physical trauma is not directly experienced, severed fibers rapidly undergo WD. Thus, mechanical nerve injury results in temporally and spatially distinct microenvironments. This was demonstrated by single cell RNAseq of immune cells captured at the nerve injury site or the distal nerve. We identified distinct, yet overlapping immune compartments, suggesting functions associated with wound healing and WD. Experiments with *Sarm1-/-* mice demonstrate that SNC triggers a spatially confined inflammatory response and accumulation of blood-borne immune cells independently of WD. Thus, physical nerve wounding and the resulting disruption of tissue homeostasis are sufficient to trigger robust local nerve inflammation. Our findings are reminiscent of a study in zebrafish larvae where laser transection of motor nerves resulted in Mac accumulation at the lesion site prior to axon fragmentation. Moreover, delayed fragmentation of severed zebrafish motor axons expressing the *Wld(s)* transgene did not alter Mac recruitment (Rosenberg et al., 2012). In injured *Sarm1-/-* mice, distal nerves are much less inflamed when compared to parallel processed WT mice. Our observation is consistent with studies in *Wld(s)* mice, where reduced nerve inflammation has been reported (Chen et al., 2015; Coleman and Hoke, 2020; Lindborg et al., 2017; Perry and Brown, 1992). Parabiosis experiments show that hematogenous WT immune cells readily enter the *Sarm1-/-* injury site, and to a much lesser extent, the distal nerve. Interestingly, Mo enter the *Sarm1-/-* distal nerve prior to fiber disintegration, however, fail to mature into Mac. This suggests that severed, but physically intact PNS fibers in the *Sarm1-/-* distal nerve release chemotactic signals for Mo. Our studies show that SNC is sufficient to trigger Mo recruitment to the distal nerve, but WD is required for Mo differentiation and full-blown nerve inflammation.

### What drives WD associated nerve inflammation?

Studies with injured *Sarm1-/-* mice show that full-blown inflammation of the distal nerve requires WD, the underlying molecular signals, however, remain incompletely understood. Because WD results in axon disintegration and simultaneous breakdown of myelin sheaths into ovoids, it is not clear whether myelin debris, SC activation, or axon fragmentation is the main trigger for WD associated nerve inflammation. Of interest, transgenic expression of Raf-kinase in mSC in adult mice is sufficient to drive SC dedifferentiation into p75-positive progenitor-like state without compromising axon integrity. SC dedifferentiation resulted in cytokine expression and nerve inflammation, however analysis of immune cell composition and comparison to SNC triggered nerve inflammation, has not yet been carried out (Napoli et al., 2012). In the healthy PNS, the endoneurial milieu is protected by the BNB, a selectively permeable barrier formed by the specialized EC along with the perineurial barrier. The BNB creates an immunologically and biochemically privileged space harboring nerve fibers and endoneurial fluid (Lim et al., 2014). ELISA of injured nerve tissue revealed that the BNB is at least partially compromised following SNC, resulting in local disturbances of vascular permeability, allowing access of serum proteins that function in opsonization and phagocytosis to the endoneurium. This suggests that in addition to degenerating axons and myelin debris, disruption of the BNB may be a driver of trauma inflicted nerve inflammation. Additional studies are needed to fully define the mechanisms that underly WD-associated nerve inflammation.

Taken together, we carried out a longitudinal analysis of injured mouse PNS, naïve nerve, and PBMC transcriptomes at single cell resolution. The study provides unprecedented insights into the dynamic cellular landscape, cell-cell interaction networks, and immune cell metabolic reprograming during the first week following nerve crush injury. To facilitate dataset mining, we developed the injured sciatic nerve atlas (iSNAT), a novel tool to navigate the cellular and molecular landscape of a neural tissue endowed with a high regenerative capacity.

## Material and Methods

### Mice and genotyping

All procedures involving mice were approved by the Institutional Animal Care and Use Committees (IACUC) of the university of Michigan (PRO 00009851) and performed in accordance with guidelines developed by the National Institutes of Health. Young adult male and female mice (8-16 weeks) on a C57BL/6 background were used throughout the study. Transgenic mice included, *Sarm1/Myd88-5-/-* (Jackson labs Stock No: 018069) and *ROSA26-mTdt/mGFP* (Jackson labs Stock No: 007576). Mice were housed under a 12 h light/dark cycle with regular chow and water ad libitum. For genotyping, ear biopsies were collected, and genomic DNA extracted by boiling in 100 µl alkaline lysis buffer (25 mM NaOH and 0.2 mM EDTA in ddH_2_O) for 30 min. The pH was neutralized with 100 µl of 40 mM Tris-HCI (pH 5.5). For PCR, 1–5 µl of gDNA was mixed with 0.5 µl of 10 mM dNTP mix (Promega, C1141, Madison, WI), 10 µl of 25 mM MgCl_2_, 5 µl of 5X Green GoTaq Buffer (Promega, M791A), 0.2 µl of GoTaq DNA polymerase (Promega, M3005), 1 µl of each PCR primer stock (100 µM each), and ddH_2_O was added to a total volume of 25 µl. The following PCR primers, purchased from *Integrated DNA Technologies,* were used: *Sarm1* WT Fwd: 5’GGG AGA GCC TTC CTC ATA CC 3’; *Sarm1* WT Rev: 5’TAA GAA TGA GCA GGG CCA AG 3’; *Sarm1* KO Fwd: 5’CTT GGG TGG AGA GGC TAT TC 3’; *Sarm1* KO Rev: 5’AGG TGA GAT GAC AGG AGA TC 3’; *Rosa26* WT Fwd: 5’-CGT GAT CTG CAA CTC CAG TC-3’; *Rosa26* WT Rev: 5’-GGA GCG GGA GAA ATG GAT ATG-3’. PCR conditions were as follows: Hot start 94°C 3 minutes; DNA denaturing at 94°C 30 seconds; annealing 60°C 1 minute; extension 72°C 1-minute, total cycles 34. Final extension for 6 min at 72°C.

### Surgical procedures

Mice were deeply anesthetized with a mixture of ketamine (100 mg/kg) and xylazine (10 mg/kg) or with isoflurane (5% induction, 2-3% maintenance, SomnoSuite Kent Scientific). Buprenorphine (0.1 mg/kg) was given as an analgesic. For SNC, thighs were shaved and disinfected with 70% isopropyl alcohol and iodine (PDI Healthcare). A small incision was made on the skin, underlying muscles separated, and the sciatic nerve trunk exposed. For sham operated mice, the nerve was exposed but not touched. For SNC, the nerve was crushed for 15 seconds, using a fine forceps (Dumont #55). The wound was closed using two 7mm reflex clips (Cell Point Scientific). Parabiosis surgery was performed as described (Kalinski et al., 2020). Briefly, before parabiosis surgery, similar aged, same sex mice were housed in the same cage for 1-2 weeks. Mice were anesthetized and their left side (host) or right side (donor) shaved and cleaned with iodine pads. A unilateral skin-deep incision was made from below the elbow to below the knee on the host and donor mouse. Mice were joined at the knee and elbow joints with non-absorbable nylon sutures. Absorbable sutures were used to join the skin around the shoulders and hindlimbs. Reflex wound clips (7 mm) were used to join the remainder of the skin between the two mice. Mice were allowed to recover for 3-4 weeks before use for SNC surgery.

### Histological procedures

Mice were euthanized with an overdose of Xylazine/Ketamine and transracially perfused for 5 min with ice-cold PBS followed by 5 min with freshly prepared ice-cold 4% paraformaldehyde in PBS. Sciatic nerve trunks were harvested and postfixed for 2 hours in ice-cold perfusion solution, followed by incubation in 30% sucrose in PBS solution at 4°C overnight. Nerves were covered with tissue Tek (Electron Microscopy Sciences, 62550– 01) and stored at −80°C. Nerves were cryo-sectioned at 14 µm thickness and mounted on Superfrost^+^ microscope slides, air dried overnight, and stored in a sealed slide box at −80°C. For antibody staining, slides were brought to RT and rinsed in 1x PBS three times, 5 min each. Slides were incubated in 0.3% PBST (1X PBS plus 0.3% Tween 20) for 10 min, followed by incubation for 1h in 5% donkey serum solution in 0.1% PBST (blocking buffer). Primary antibodies at appropriate dilutions were prepared in blocking solution, added to microscope slides, and incubated at 4°C overnight. The next day, sections were rinsed three times in PBS, 5 min each. Appropriate secondary antibodies in blocking buffer were added for 1h at a dilution of 1:1000 at room temperature. Slides were rinsed three times in PBS, 5 min each, once in 0.1% PBST for 5 min, and briefly with milliQ water. Sections were mounted in DAPI containing mounting medium (Southern biotec (Cat. No.0100-20)), and air dried. Images were acquired with a Zeiss Apotome2 microscope equipped with an Axiocam 503 mono camera and ZEN software. Image processing and analysis were conducted using the ZEN software.

For *in situ* mRNA detection, the RNAscope Multiplex Fluorescent Reagent Kit v2 (ACD, 323100) was used. Microscope slides with serially cut nerves were rinsed in 1x PBS for 5 min and air dried by incubation at 60°C for 45 min in an oven (VWR Scientific, Model 1525 incubator). Next, tissue sections were fixed in 4% Paraformaldehyde (PFA)/PBS for 45 mins at RT and dehydrated by incubation in a graded series of 50%, 70%, and 100% ethanol for 5 min each. Sections were air dried for 5 min at RT and one drop of hydrogen peroxide solution (ACD catalog number: PN 322381) was added to each nerve section on each slide and incubated at RT for 10 min. Sections were then submerged in 99°C RNase free water for 1 min, followed by incubation in 99°C 1x antigen retrieval solution (ACD catalog number: 322000) for 20 min. Next, slides were air dried by incubation for 45 min in an oven at 60°C. Protease III solution (ACD catalog number: PN 322381) was applied to tissue sections followed by incubation at 40°C in an ACD hybridization oven (ACD catalog number: 321710) for 45 min. RNA probes were mixed at appropriate ratios and volumes (typically 50:1 for C1:C2) for complex hybridization. For single RNA probe hybridization, RNA probes were diluted with probe dilutant at 1:50-1:100 (ACD catalog number: 300041). Appropriate probes or the probe mixtures were applied to tissue sections and incubated for 2h in the hybridization oven at 40°C. 1X wash buffer was prepared from a 50X stock solution (ACD catalog number: PN 310091) and sections rinsed for 2 min. The slides were then stored in 5X SSC buffer overnight. The next morning, sections were rinsed in 1X wash buffer and amplification probes, corresponding to the primary RNA probes, were applied starting with the C1 probe (ACD catalog number: 323100). Slides were incubated in the hybridization oven for 30 min 40°C and then rinsed twice with 1X wash buffer. Next, the A2 and A3 probes were applied. For development, the TSA system (AKOYA, Cy3: NEL744001KT; Cy5: NEL745001KT; Fluorescein: MEL741001KT) was used. Once the color for probe C1 was selected, HRPC1 solution (ACD catalog number: 323120), it was applied to the appropriate sections and incubated for 15 mins in the hybridization oven at 40°C. The sections were then rinsed in 1x washing buffer. Designated TSA color for probe C1, diluted in the TSA dilutant (ACD catalog number: 322809) between 1:1000-1:2000 was applied to the respective sections and incubated for 30 mins in the ACD hybridization oven at 40°C. Sections were rinsed in 1X wash buffer and then HRP blocker (ACD catalog number: 323120) was applied and incubated for 15 min in the ACD hybridization oven at 40°C. This procedure was repeated for probes C2 and C3 as needed using HRPC2 and HRPC3 respectively. Sections were mounted in DAPI southern biotech mounting media (Cat. No.0100-20), air dried, and imaged or stored at 4°C in the dark. For quantification of labeled cells in nerve tissue sections, a field of view (FoV) was defined, 200µm X 500µm at the injury site and the distal nerve. The FoV in the distal nerve was 2000 µm away from the injury site. The number of labeled cells per FoV was counted. Only labeled cells with a clearly identifiable nucleus were included in the analysis. The number of cells counted per FoV was from n= 2 mice and n= 2 technical replicates per mouse.

### Preparation of single-cell suspensions for flow cytometry and scRNAseq

Mice were euthanized and transcardially perfused with ice-cold PBS to reduce sample contamination with circulating leukocytes. Sciatic nerve trunks from naïve and injured mice were harvested. For injured mice, a segment was collected that includes the injury site and distal nerve just before to the trifurcation of the tibial, sural, and peroneal nerves. Nerves were placed in ice-cold PBS containing actinomycin D [45 µM, Sigma Aldrich, A1410]. Some nerves were further dissected into 3 mm segments, either encompassing the site of nerve injury, ∼1.5 mm proximal to ∼1.5 mm distal of the crush site, or distal nerve, located between +1.5 to +4.5 mm away from the crush site. Nerves from 3 mice (6 mm segments) or 5-6 mice (3 mm segments) were pooled for each biological replicate. Nerves were minced with fine scissors and incubated in 1 ml PBS supplemented with collagenase (4 mg/ml Worthington Biochemical, LS004176) and dispase (2 mg/ml, Sigma-Aldrich, D4693) and incubated for 30–45 min at 37°C in a 15-mL conical tube. For scRNAseq, the digestion buffer also contained actinomycin D (45 µM). Nerves were triturated 20x with a 1000 µl pipette every 10 min and gently agitated every 5 min. Next, nerves were rinsed in DMEM with 10% FBS, spun down at 650 g for 5 min, the resulting pellet resuspended, and fractionated in a 30% Percoll gradient. For flow cytometry, the cell fraction was collected and filtered through a pre-washed 40 µm Falcon filter (Corning, 352340) and cells were pelleted at 650 g for 5 min at 4°C. Immune cell populations were identified with established antibody panels as described (Kalinski et al., 2020). For scRNAseq, the cell suspension was cleared of myelin debris with myelin removal beads (Miltenyi, 130-096-733), and cells resuspended in Hanks balanced salt solution (Gibo, 14025092) supplemented with 0.04% BSA (Fisher Scientific, BP1600). To enrich for immune cells in nerve specimens or from peripheral blood, some samples were run over an anti-CD45 or anti-CD11b column (Miltenyi, 130-052-301). Cells were counted and live/dead ratio determined using propidium iodine staining and a hemocytometer (Kalinski et al., 2020). Blood was collected from adult naïve mice by cardiac puncture and collected into K2 EDTA coated tubes (BD 365974) to prevent coagulation. Approximately 500 µl of blood was passed through a 70 µm cell strainer in 5 ml ACK (Ammonium-Chloride-Potassium) lysis buffer. Blood was incubated at room temperature for 5 min and erythrocyte lysis stopped by addition of 15 mL FACS buffer, followed by leukocyte spin down in a clinical centrifuge. This process was repeated 3 times for complete erythrocyte lysis.

### Barcoding and library preparation

The Chromium Next GEM Single Cell 3’ Reagent kit v3.1 (Dual Index) was used. Barcoding and library preparation was performed following the manufacturer’s protocols. Briefly, to generate single-cell gel-bead-in-emulsion (GEMs) solution, approximately 15,000 cells, in a final volume of 43 µl, were loaded on a Next GEM Chip G (10x Genomics) and processed with the 10x Genomics Chromium Controller. Reverse transcription was performed as follows: 53°C for 45 minutes and 85°C for 5 minutes in a Veriti Therml Cycler (Applied Biosystems). Next, first strand cDNA was cleaned with DynaBeads MyOne SILANE (10x Genomics, 2000048). The amplified cDNA, intermedium products, and final libraries were prepared and cleaned with SPRIselect Regent kit (Beckman Coulter, B23318). A small aliquot of each library was used for quality control to determine fragment size distribution and DNA concentration, using a bioanalyzer. Libraries were pooled for sequencing with a NovaSeq 6000 (Illumina) at an estimated depth of 50,000 reads per cell, yielding 11.3 billion reads. Novaseq control software version 1.6 and Real Time Analysis (RTA) software 3.4.4. were used to generate binary base call (BCL) formatted files.

### Data availability

All scRNA-seq datasets will be made available online through the Gene Expression Omnibus (GEO) database.

### scRNAseq data analysis

Raw scRNAseq data were processed using the 10x Genomics CellRanger software version 3.1.0. The CellRanger “mkfastq” function was used for de-multiplexing and generating FASTQ files from raw BCL. The CellRanger “count” function with default settings was used with the mm10 reference supplied by 10x Genomics, to align reads and generate single cell feature counts. CellRanger filtered cells and counts were used for downstream analysis in Seurat version 4.0.5 implemented in R version 4.1.2. Cells were excluded if they had fewer than 500 features, more than 7500, or the mitochondrial content was more than 15%. For each post-injury time point, reads from multiple samples were integrated and normalized flowing a standard Seurat SCTransform+CCA integration pipeline (Hafemeister and Satija, 2019). The mitochondrial mapping percentage was regressed out during the SCTransform normalization step. Principal component analysis was performed on the top 3000 variable genes and the top 30 principal components were used for downstream analysis. A K-nearest neighbor graph was produced using Euclidean distances. The Louvain algorithm was used with resolution set to 0.5 to group cells together. Non-linear dimensional reduction was done using UMAP. The top 100 genes for each cluster, determined by Seurat’s FindAllMarkers function and the Wilcoxon Rank Sum test, were submitted to Qiagen’s Ingenuity Pathway Analysis (IPA) software – version 70750971 (Qiagen Inc., https://digitalinsights.qiagen.com/IPA) using core analysis of up- and down-regulated expressed genes. Top-scoring enriched pathways, functions, upstream regulators, and networks for these genes were identified utilizing the algorithms developed for Qiagen IPA software (Kramer et al., 2014), based on Qiagen’s IPA database of differentially expressed genes.

### Comparative analysis of cell identities at different post-injury time points

Comparison of cell identities between time points was done using the Seurat technique for classifying Cell Types from an integrated reference. This technique projects the PCA structure of the reference time point onto the query time point. This is similar to Seurat’s implementation of Canonical Correlation Analysis (CCA) in that it creates anchors between the two data sets, but it stops short of modifying the expression values of the query. The output of this technique is a matrix with predicted IDs and a prediction score between 0 and 1. For each reference cell type we used the geometric mean of prediction scores for cells predicted with that type in the query set. This single prediction score was used as a surrogate for confidence in the same cell state existing in the query cells. Alternatively, using the overlapping, top 3,000, highly variable genes from each dataset, we computed the Pearson correlation between the genes’ average expression (log2 of uncorrelated “RNA” assay counts) from each reference cell type and its predicted query cells. As reported in Results, we projected 1dpc cell types onto 3dpc as well as 3dpc onto 7dpc.

### Cell Chat

CellChat version 1.1.3, with its native database, was used to analyze cell-cell communication (Jin et al., 2021) (Suoqin Jin et al., https://doi.org/10.1038/s41467-021-21246-9) A truncated mean of 25% was used for calculating average mean expression, meaning a gene was considered to have no expression in a cell type if fewer than 25% of cells in that cell type expressed the gene.

### SlingShot

Slingshot version 2.2.1 was used to model trajectories in the integrated myeloid dataset. The PCA embeddings from the first 4 principal components were used as input and the beginning of the trajectory anchored at the Monocytes. The Mo to Mac and Mo to MoDC trajectories were selected as interesting. To predict the genes that contribute most to pseudotime we used the tidymodels R package (Hadley Wickham and Max Kuhn www.tidymodels.org); specifically, regression models in random forests with 200 predictors (genes) randomly samples at each split and 1,400 trees. The Impurity method was used to calculate a genes importance in predicting the pseudotime. Heatmaps of the top 30 genes by importance were used to visually examine how the gene changes in cells ordered by their pseudotime.

### Code Availability

iSNAT, an interactive web application was generated with the RStudio’s shiny package (http://github.com/rstudio/shiny). The Dashboard format was generously supplied by the RStudio group https://rstudio.github.io/shinydashboard/ The code for all analysis and Rshiny server is available from GitHub(https://github.com/GigerLab/iSNAT) --Not yet populated).

### Protein analysis

For protein analysis, mice (naïve, and at 1, 3 and 7dpc) were euthanized and transcardially perfused with ice-cold PBS for 5 minutes. Naïve and injured sciatic nerves were dissected, with nerves from 3 mice pooled per time point in ice-cold PBS with 1% protease inhibitor cocktail (Sigma-Aldrich, P8340). Samples were minced, homogenized in 1% Triton X-100 (Sigma-Aldrich, T8787). Samples were frozen at −80°C, thawed, centrifuged at 10,000 x g for 5 min to pellet cellular debris. The protein concentrations of the nerve supernatants and serum were determined using a BCA protein concentration assay. Western blot analysis of sciatic nerve tissue was carried out as described previously (Kalinski et al., 2020). For some WT and *Sarm1-/-* mice, injured nerves were divided into proximal, injury site, and distal segments. For the proteome Profiler Mouse XL Cytokine array, 200 µg of protein was applied to each ELISA membrane and developed according to the manufacturer’s instructions (Proteome Profiler Mouse XL Cytokine Kit, ARY028, R&D Systems, Minneapolis, MN, USA). Cytokine array signals were detected by X-ray film, scanned, and quantified using LI-COR Image Studio software – version 5.2.5. Cytokine signals (pixel-density value) from duplicate spots were averaged then normalized to the reference spots in the upper right and left corners and the lower left corner on the membrane. Representative images of array membranes are shown (n = 1-2 biological replicates per condition).

### qRT-PCR

Quantitative PCR (qPCR) was carried out in triplicate with SYBR Green Fluorescein Master Mix (Thermo Scientific Cat. No. 4364344) on a QuantStudio 3 real-time PCR system (Applied Biosystems Cat. No A28567). The ΔΔCt method was used to determine the relative expression of mRNAs, which was normalized to *Rlp13a* mRNA levels. Mice were transcardially perfused with ice-cold PBS, sciatic nerve trunks harvested, and collected in an RNase free 1.5 mL Eppendorf tube in 1 ml TRIzol solution (Thermofisher Cat. No. 15596026). Nerves were minced into small pieces with RNase free spring scissors, homogenized using a pestle motor mixer (RPI, 299200) and frozen at −80°C overnight. The next day, specimens were thawed, placed on ice, and 0.2 ml of chloroform was added to the TRIzol mix and shaken thoroughly for 5 min. Samples were centrifuged at 12000 x g, 4°C for 15 min in a tabletop centrifuge. The aqueous phase was removed and placed in a new Eppendorf tube. Glycogen (2 µl) and isopropanol (0.5 ml) were added to the aqueous solution and mildly shaken with a shaker at 4°C for 10 mins. Samples were centrifuged at 12000 x g at 4°C for 10 min to precipitate total RNA. The supernatant was discarded, and the pellet resuspended in 1 ml 75% ethanol, vortexed briefly and then centrifuged 4°C for 5 mins at 7500 x g. The supernatant was discarded, the pellet airdried for 1-2 h, and resuspended in 20 µl of RNase free water. RNA yield was quantified with a nanodrop (Thermo Scientific, Nanodrop One), RNA aliquoted, and stored at −80°C.

RNA was reverse transcribed into first stand cDNA using the Invitrogen SuperScript™ III First-Strand Synthesis System kit (Cat. No. 18080051). Briefly, to 250 ng of total RNA, 1 µl of 50 µM oligo(dT)20 primer, 1 µl of 10 mM dNTP mix were added and the final volume adjusted to 10 µl using RNase free water. The RNA mix was incubated at 65°C for 5 min and quickly placed on ice for at least 5 mins. In a separate tube, a master mix was prepared containing 2 µl of 10x RT buffer, 4 µl of 25 mM MgCl_2_ and 2 µl of 0.1M DTT. 8 µl of the master mix was added to 10 µl of RNA mix with 1 µl of SuperScript III reverse transcriptase and 1 µl of RNaseOut enzyme. The reverse transcription reaction (final volume 20 µl) was carried out at 50°C for 50 min, followed by a 5 min incubation at 85°C for 5 mins. RNase H (1 µl) was added to the first stand cDNA and incubated at 37°C for 20 mins. First strand cDNA was quantified using a Nanodrop One and stored in aliquots at −20°C. The cDNA was diluted to 50 ng/ul and used for qPCR in a 96 well plate format. SYBR Green Fluorescein Master Mix 10 µl (Thermo Scientific Cat. No. 4364344), 0.4 µl of forward primer, 0.4 µl of reverse primer, first strand cDNA (50 ng) was mixed, and the final volume adjured to 20 µl per well with milliQ water. The plate was sealed, centrifuged at 2500 x g for 1 min, and then placed in QuantStudio 3 real-time PCR system (Applied Biosystems Cat. No A28567) to run the qPCR. The protocol for running was as follows. Data was exported in Excel format and analyzed in Excel using the ΔΔCt method where the gene control was *Rlp13a* while the reference control was the sciatic nerve in naïve condition.

### Statistical Methods

Data are presented as mean ±SEM. Statistical analysis was performed in GraphPad Prism (v7) using paired or unpaired 2-tailed Student’s t test, or 1-way or 2-way ANOVA with correction for multiple comparisons with Tukey’s post-hoc test, as indicated in the figure legends. A p-value < 0.05 (*) was considered significant. p<0.01 (**), p<0.001 (***), and p<0.0001 (****). Data acquisition and analysis was carried out by individuals blinded to experimental groups.

## Acknowledgments

We thank members of the Giger lab for critical reading of the manuscript and Qing Wang for excellent technical support. This work was supported by NIH T32 NS07222, Training in Clinical and Basic Neuroscience (AK), NIH T32 GM113900, Training Program in Translational Research (LH). The National Institutes of Health, MH119346 (RG), R01DC018500 (GC), the University of Michigan MICHR seed funds (RG and GC), and the Dr. Miriam and Sheldon G Adelson Medical Research Foundation (RK, JT, DG, RG).

